# Hierarchical Interpretation of Out-of-Distribution Cells Using Bottlenecked Transformer

**DOI:** 10.1101/2024.12.17.628533

**Authors:** Qifei Wang, He Zhu, Yiwen Hu, Yanjie Chen, Yuwei Wang, Xuegong Zhang, James Zou, Manolis Kellis, Yue Li, Dianbo Liu, Lan Jiang

## Abstract

Identifying the genetic and molecular drivers of phenotypic heterogeneity among individuals is vital for understanding human health and for diagnosing, monitoring, and treating diseases. To this end, international consortia such as the Human Cell Atlas and the Tabula Sapiens are creating comprehensive cellular references. Due to the massive volume of data generated, machine learning methods, especially transformer architectures, have been widely employed in related studies. However, applying machine learning to cellular data presents several challenges. One such challenge is making the methods interpretable with respect to both the input cellular information and its context. Another less explored challenge is the accurate representation of cells outside existing references, referred to as out-of-distribution (OOD) cells. The out-of-distribution could be attributed to various physiological conditions, such as comparing diseased cells, particularly tumor cells, with healthy reference data, or significant technical variations, such as using transfer learning from single-cell reference to spatial query data. Inspired by the global workspace theory in cognitive neuroscience, we introduce CellMemory, a bottlenecked Transformer with improved generalization capabilities designed for the hierarchical interpretation of OOD cells unseen during reference building. Even without pre-training, it exceeds the performance of large language models pre-trained with tens of millions of cells. In particular, when deciphering spatially resolved single-cell transcriptomics data, CellMemory demonstrates the ability to interpret data at the granule level accurately. Finally, we harness CellMemory’s robust representational capabilities to elucidate malignant cells and their founder cells in different patients, providing reliable characterizations of the cellular changes caused by the disease.

## Introduction

In recent years, single-cell sequencing has revolutionized genomics by providing unprecedented insights into cellular heterogeneity and dynamics. International consortia like Human Cell Atlas and Tabula Sapiens are utilizing population-scale single-cell resources to create a consensus reference^1, 2^ reminiscent of the Human Genome Project^3^ but at a cellular level^4^. However, cells from different individuals, conditions, technologies, or species do not exhibit similar distribution patterns. For example, there are variations in the states of malignant and healthy cells, as well as heterogeneity among malignant cells across different patients. Moreover, significant technical variations are present between single-cell and spatial omics data. Cells that deviate substantially from the established paradigms are defined as out-of-distribution (OOD) cells^5^.

Deciphering of OOD cells requires concurrent identity inference and data integration, a process known as reference mapping^5^. Currently, only a very limited number of methods are capable of doing both^6–9^ simultaneously. Furthermore, these methods often fall short of adequately considering gene interactions, which are crucial for generalizing cell representations in large-scale, high-resolution single-cell atlases. For biological researchers, the insights provided by explainable artificial intelligence (xAI) frequently surpass the value of predictions, as they can enable novel understandings of life processes^10^. The application of xAI to cells outside the reference scope is an area that has not been thoroughly explored^10, 11^.

Large Language Models (LLMs) based on Transformer architectures, capable of understanding the interrelationships among features, present a novel strategy for exploring data across various fields^12–15^. However, current single-cell LLMs^16–18^ struggle with handling long token inputs due to the computational complexity of self-attention mechanisms. This limits their capacity to comprehend the regulatory relationships among genes and to elucidate comprehensive cell representations. In this study, we developed a neuroscience-inspired transformer architecture to specifically solve these problems.

The Global Workspace Theory (GWT) in neuroscience suggests that consciousness emerges from the selective dissemination of information via a shared “global workspace”^19–21^ or “memory space”. This workspace, a network of densely interconnected neurons, facilitates competition between neuron modules to write information to this limited-capacity space^22^. Inspired by this concept, we propose that combining GWT with a Transformer architecture can efficiently manage informative and sparse single-cell data. We have developed CellMemory, which is a bottlenecked architecture within the Transformer that uses a cross-attention mechanism. This unique design allows CellMemory to learn generalized representations from standardized paradigms and perform interpretable inferences for OOD cells. By incorporating various specialist modules into the global workspace transparently, CellMemory allows for a hierarchical interpretation of the model, which can help uncover both commonalities and distinctive representations of these cells. Furthermore, the simulation of the consciousness formation process in the global workspace enhances our understanding of how the deep learning model systematically processes biological information.

A comprehensive benchmarking study compared 18 task-specific methods to CellMemory utilizing data from over 15 million cells. CellMemory demonstrated harmonious integration and accurate label transfer, even outperforming LLMs^16, 17^ that were pre-trained with tens of millions of cells regarding generalization ability and computational efficiency^23^. Remarkably, CellMemory can provide interpretable identity representations for single-cell spatial transcriptomics at a granular level.

Finally, we used CellMemory and healthy references to delineate out-of-distribution (OOD) malignant cells. CellMemory helps us contextualize malignant cells in harmonious embeddings and explain their origins. Our study focused on a transitional stage in the development of lung cancer, revealing that tumor cells in some lung cancer patients originated from different founder cells. This finding indicates that understanding the cell of origin of a tumor in some patients is crucial for comprehending drug resistance. It underscores the importance of leveraging explainable AI (xAI) and existing references to better understand complex diseases by offering valuable insights into OOD disease states.

## Results

### The inspiration for CellMemory

The Global Workspace Theory in neuroscience suggests that consciousness arises from the selective dissemination of information through a shared “global workspace”^19–21^. The brain consists of numerous specialized modules that handle various tasks like vision, hearing, memory, and emotions. The “global workspace” or “Memory space” is a network of densely interconnected neurons that enables these specialized modules to share and communicate information. Due to its limited capacity, not all information can be retained in the global workspace. Specialized modules compete to write information to the global workspace, which then becomes part of conscious awareness. Once stored, the information is broadcasted to diverse specialized neural modules via synchronized neural firing and rapid signaling. This ensures that various brain regions process the information concurrently, contributing to a unified conscious experience.

Inspired by GWT in cognitive neuroscience, we propose that an architecture comprising specialists trained to communicate effectively through the constraint of a global workspace yields a substantial enhancement in computational efficiency and generalization^12^. To achieve this, we have developed CellMemory, which utilizes a bottlenecked architecture within a Transformer-based framework (Fig. 1a). CellMemory is tailored to derive meaningful cellular representations from single-cell datasets and effectively address the challenge of making inferences about out-of-distribution (OOD) cells in an explainable manner. Furthermore, it can integrate population-scale single-cell datasets at a granular level, adeptly managing variations in cell state, technical platform, spatiotemporal conditions, ethnicity, and species (Fig. 1b).

**Fig 1.**
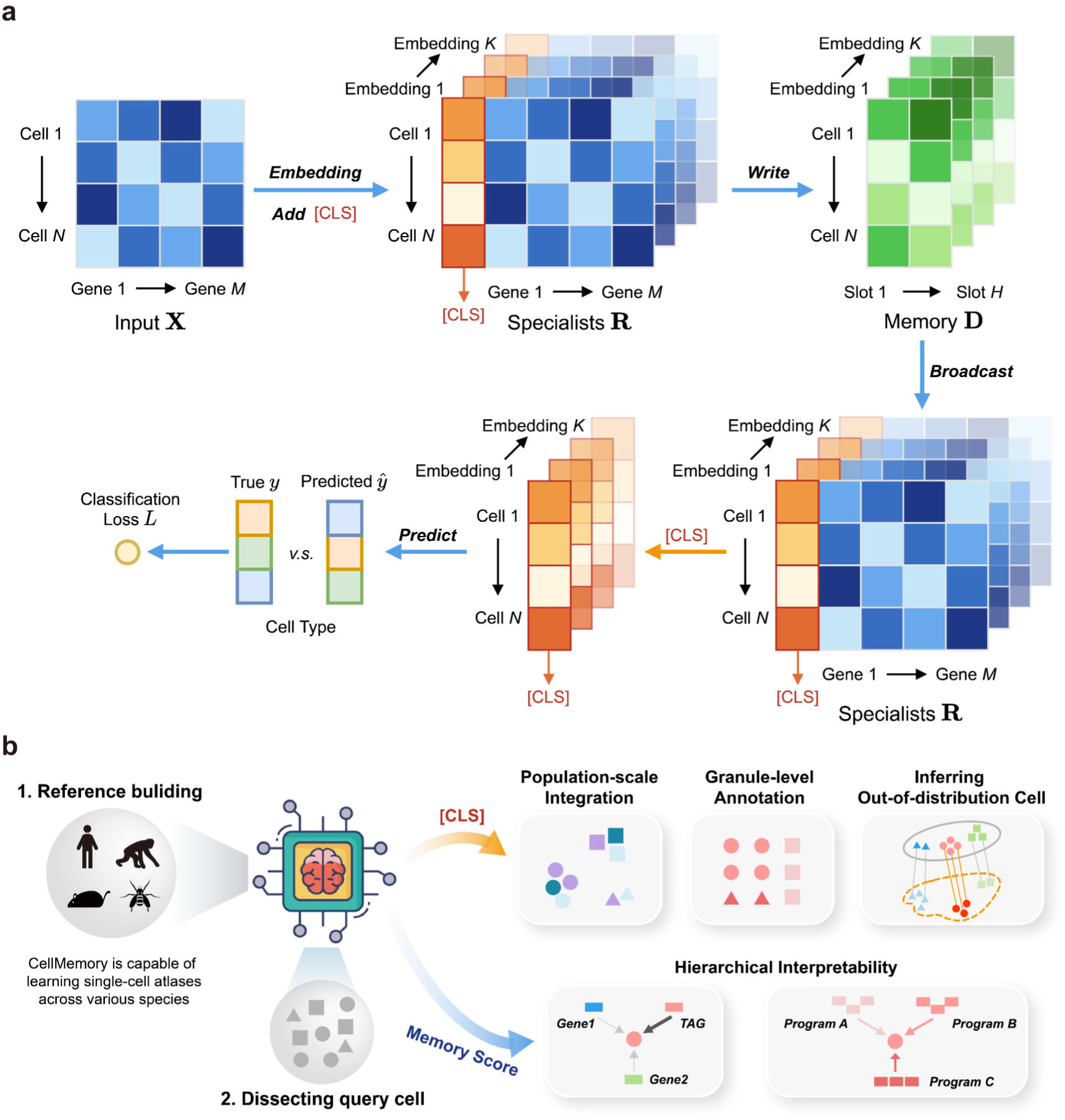
The illustration of CellMemory. **a,** CellMemory learns cell representations from references in a supervised manner. The densely interconnected neurons in CellMemory enable specialized modules to share and communicate information. The “Specialists” (specialized modules) compete with each other to write information to the “Memory”. The information is then broadcasted to “Specialists” through synchronized neural firing and rapid signaling. The processes of writing and broadcasting between “Specialists” and “Memory” are executed by the cross-attention mechanism. The CLS token of CellMemory is used for training the model by comparing predictions with the real labels. **b,** The application of CellMemory. CellMemory is capable of training models using single-cell data from any species without being constrained by a predefined feature set. Upon analysis of the query set with the trained CellMemory, the generated cell embeddings [CLS] enable population-scale integration (in the illustration, shapes represent cell types, and colors represent batch information), granule-level annotation (both shapes and colors represent cellular identity, and similar colors represent similarities in cell identities), and inference of out-of-distribution cells (both shapes and colors represent cellular identity). While performing these analyses with cell embeddings, CellMemory also generates the memory score, which allows for the interpretation of model decisions at the level of specific genes and gene programs. TAG: Top Attention Genes by memory score.

CellMemory employs a strategy of natural language processing (NLP) models for generating token and position embeddings^13^. To characterize features such as genes, position embedding is employed, while gene expression is processed in a bag-of-words manner and characterized by token embedding (Methods). The feature matrix is then transformed by the embedding layer into the “Specialists” module. Concurrently, the CLS token is incorporated into the “Specialists” module as the cell representation (Fig. 1a).

The conventional self-attention mechanism^12, 13^ used by BERT has a quadratic complexity concerning sequence length, resulting in increased time and memory complexity^24^ as sequences become larger. This presents a particularly difficult challenge when dealing with large-scale single-cell omics data. In contrast, CellMemory uses the “Specialists” module (length=*M*) to perform cross-attention with the “Memory” (length= *H*, *H* ≪ *M* ) (Fig. 1). In this process, a significant amount of biological information competes for the limited “Memory space” (Supplementary Fig. 1a). The refined information is then broadcast to all modules by the “Memory” (Supplementary Fig. 1b) (Methods). The bottleneck of “Memory space” acts as a filter, prioritizing the most significant information and ensuring that the most important biological details are efficiently communicated. As a result, the CellMemory architecture not only has impressive generalization capabilities but also substantially reduces computational costs (Methods).

During the inference process, CellMemory empowers the allocation of different weights to various tokens, or genes, within the input based on their determined significance through memory scores. Simultaneously, it ensures the consideration of the cellular context when integrating all scores into memory slots (Methods). This unique architecture equips the model to provide interpretability at both gene and gene program levels (Fig. 1b, Supplementary Fig. 1c), thus enabling a more insightful understanding of specific cellular states.

### CellMemory enables one-stop reference mapping of single-cell atlas across diverse scenarios

CellMemory’s annotation performance was assessed by comparison with task-specific methods^6, 7, 16–18, 25, 26^ using over 4.6 million cells^26–34^ with diverse biological and technological attributes (Supplementary Fig. 2a). The model was trained on cells from one condition and subsequently predicted outcomes from distinct states, covering variations in platform, age, organ, or species. Performance metrics included the F1-score (macro) and accuracy, with the F1-score offering a nuanced evaluation of predictive performance for rare categories in single-cell datasets. Remarkably, CellMemory outperformed LLMs^16, 17^ in various tasks, including identifying rare cell types on most benchmark datasets^35^, without resource-consuming pre-training (Fig. 2a, Supplementary Fig. 2-3). While pre-trained LLMs may demonstrate improved accuracy on small-scale datasets, an imbalance in cell proportions during the pre-training phase likely leads to neglecting low-abundance cell types (Fig.2a, Supplementary Fig. 2b-c). In contrast, CellMemory exhibited excellent generalization capabilities (Fig. 2a), and processed data more effectively than self-attention-based Transformers (Fig. 2b, Supplementary Fig. 2d).

**Fig 2.**
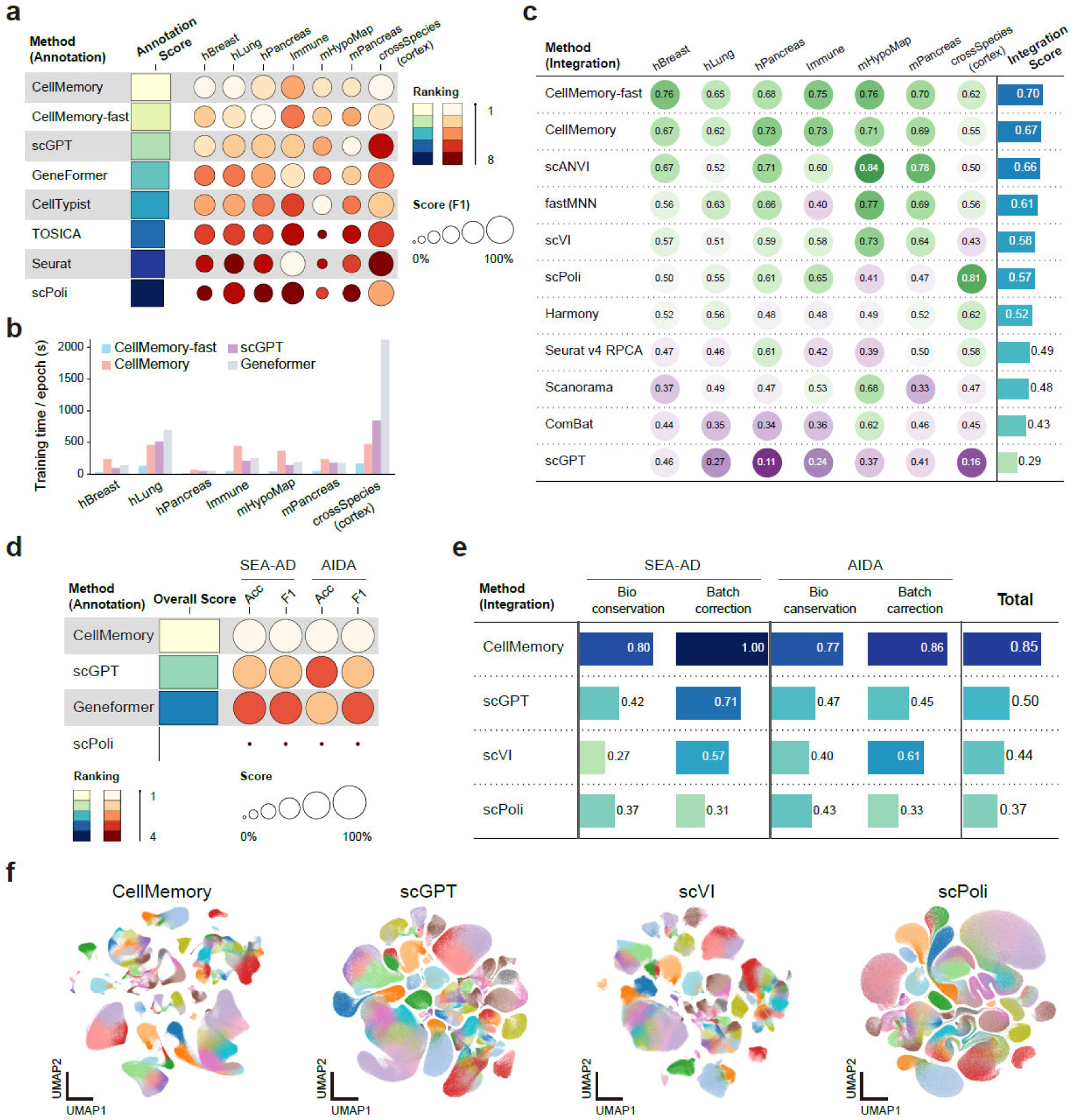
Annotation and integration benchmark. **a,** F1-score (macro) is employed to assess the annotation performance of single-cell datasets. Each circle represents the mean of the five-fold cross-validation for the respective method (except for crossSpecies, which includes data from five species). The overall performance across all datasets corresponds to the Annotation Score. *hBreast*: Prediction of human breast cancer samples using normal breast samples. *hLung*: Prediction of human LUSC samples using normal lung samples. *hPancreas*: Prediction of human Type 1 diabetes samples using pancreatic islets normal control samples. *Immune*: Immune cells from human tissue group 1 were used as the training set, and immune cells from human tissue group 2 were used as the test set. *mHypoMap*: Prediction of mouse hypothalamus samples produced by the Drop-seq platform, using samples produced by the 10x platform. *mPancreas*: Prediction of mouse type 1/2 diabetes samples using normal pancreatic islet samples. *crossSpecies (cortex)*: The human cortex samples were used as the training set, while one human and four non-human primate cortex samples were used as the test set. CellMemory: The model was trained using all input genes, considering genes with zero expression in each cell. CellMemory-fast: Genes with zero expression in each cell were filtered out, and the model was trained using the Mixed Precision Training strategy. **b,** The average time taken to train one epoch is recorded for each dataset for CellMemory, scGPT, and Geneformer. **c,** The integration benchmark for the dataset includes popular integration methods such scPoli, scVI, and scGPT. The Integration Score is comprehensively evaluated based on the retention of biological information and the removal of batch effects. The overall score is calculated as a weighted average of the batch correction score and the biological conservation score, with weights of 0.4 and 0.6, respectively. **d-e,** Annotation and integration benchmarks for the SEA-AD and AIDA datasets. **f**, UMAPs are generated from SEA-AD cell embeddings obtained by four integration methods. Cell colored by original annotation.

The performance of CellMemory in integrating cross-state or cross-platform datasets during reference mapping was evaluated using cell embeddings generated from the datasets to be integrated (The same CLS embedding used as in the annotation results above, details in Methods). Comparisons were conducted with advanced integration tools^36^ such as scVI^37^, LLM scGPT^16^, and scPoli^6^, which is specifically tailored for the integration of population-scale datasets. Benchmark results demonstrate that CellMemory effectively preserves biological representations while minimizing batch effects (Fig. 2c, Supplementary Fig. 4). In seven benchmark datasets, CellMemory outperformed other tools in more than half of them. Recent research has presented discrepancies in the token preprocessing strategies for feeding single-cell data into NLP models. To assess the impact of retaining genes with zero expression and using different token preprocessing strategies on the integration performance of NLP models, we conducted experiments on benchmark datasets (Supplementary Fig. 5). Our evaluations indicate that retaining features with zero expression is beneficial for integration, underscoring the significance of efficiently considering longer tokens for single-cell data integration. Moreover, scaling the expression values of each sample into bins within defined ranges aids in the removal of batch effects. This suggests that a scale-to-bin strategy is more effective for batch correction when integrating data with significant batch effects.

As sequencing technologies become more advanced and affordable, researchers can now gather larger amounts of data from specific tissues. This allows them to explore more subtle cell types or states using detailed single-cell data. Typically, data integration and annotation strategies are iterative^32, 33^, beginning with distinguishing cell types at a broad level and then progressing to finer distinctions between subtypes or even supertypes within each major category. In this investigation, we sought to validate the capacity of CellMemory to achieve one-stop reference mapping for two population-scale datasets characterized by high-resolution cell identities (Fig. 2d-e). The first atlas features the brain cortex of Alzheimer’s patients (SEA-AD), with 131 categories of granule-level annotation^32^. This atlas contains 1,395,601 cells from 84 donors, with normal samples serving as the reference and disease samples as the inference object. Another atlas, the Asian Immune Diversity Atlas (AIDA)^38^, encompasses 1,058,909 peripheral blood mononuclear cells from 503 healthy donors across five Asian ancestries and a European population. We trained the model using cells from the Korean population and integrated data from the other five populations (Methods). In the annotation benchmark, CellMemory demonstrated superior performance in both accuracy and F1-score (Fig. 2d). In the context of the SEA-AD dataset, which contains 131 cell types, CellMemory achieved an accuracy of nearly 90%, while scPoli only reached 60%. The integration results reveal that CellMemory not only removes batch effects but also preserves the most comprehensive biological information, ranking first in 7 out of 8 metrics (Fig. 2e-f, Supplementary Fig. 6a, 7). These results demonstrate that CellMemory surpasses both task-specific methods and pre-trained LLMs in terms of data comprehension capabilities when conducting high-resolution knowledge transfer for population-scale datasets.

### CellMemory facilitates the interpretable characterization of single-cell spatial omics

The deciphering of single-cell spatial omics poses a challenge due to significant technical bias and variations of information richness compared to single-cell omics data. To address this challenge as one of the problems of characterizing OOD cells, we propose the application of CellMemory’s robust representation capabilities for characterizing single-cell spatial transcriptomics (Fig. 3a). A benchmarking was performed to assess the efficacy of spatial annotation using data from CosMx^33, 39^, MERFISH^34, 40^, and Slide-seq^41^. The study compared CellMemory with various spatial methods^42–46^ and two LLMs (Methods). Accuracy was employed as the primary metric for evaluation due to the complexity of spatial data representation. The results demonstrated that CellMemory performs the best, even in the absence of spatial coordinate information (Fig. 3b, Supplementary Fig. 8). The performance of scGPT and Geneformer indicates that pre-training strategies could potentially improve the comprehension of spatial data across species.

**Fig 3.**
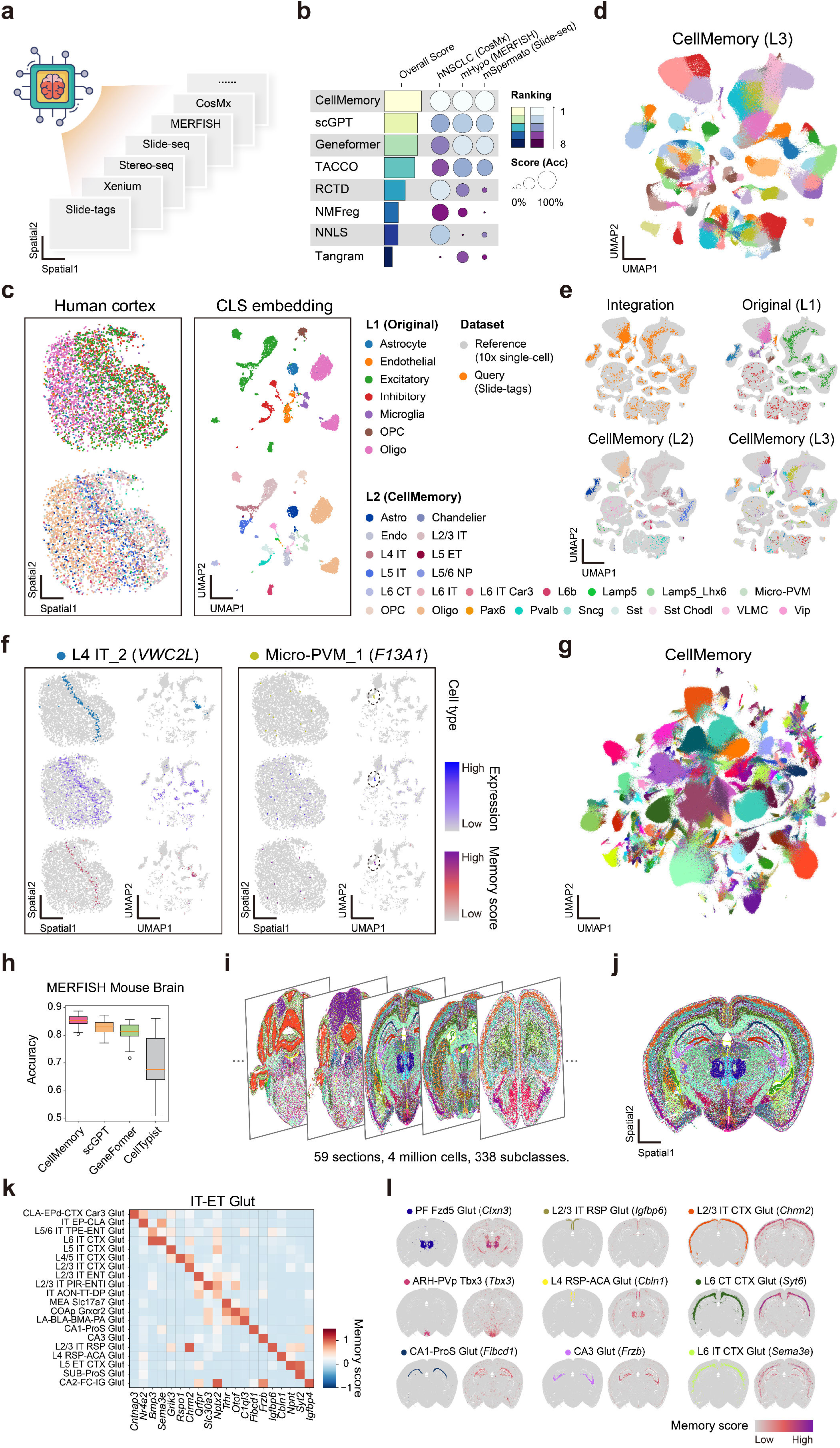
CellMemory interprets single-cell spatial data across omics. **a,** CellMemory can characterize single-cell spatial omics from various sequencing platforms, including CosMx, MERFISH, Slide-seq, Stereo-seq, Xenium, and Slide-tags. **b,** The accuracy of spatial annotation tools is evaluated using datasets from mHypo (mouse hypothalamus measured by MERFISH), hNSCLC (human NSCLC measured by CosMx), and msSermato (mouse spermatogenesis measured by Slide-seq). Each dataset comprised three samples for replication. **c,** CellMemory was trained using single-cell data at the L2 cell type resolution, to generate CLS embeddings and annotations for Slide-tags cells. The right part is plotted by the L1 (original) and L2 (CellMemory) cell types in spatial coordinates. **d,** CellMemory model was built using single-cell data at the L3 resolution, to integrate single-cell and Slide-tags data. **e,** The cells highlighted in the co-embedding are Slide-tags cells, labeled with Slide-tags cell, L1 (original), L2 (CellMemory), and L3 (CellMemory) cell identity. **f,** The identification of Slide-tags cells at L3 resolution by CellMemory (L4 IT_2 and Micro-PVM_1) is displayed (the first line), along with the expression (the second line) and memory score (the third line) of TAGs (*VWC2L, F13A1*). The left half represents the spatial coordinate, and the right half is the UMAP coordinate. **g,** UMAP representation of 4 million mouse whole brain cells from MERFISH, colored by subclasses. **h,** Annotation benchmark comparison of CellMemory with other state-of-art methods, including scGPT, Geneformer, and CellTypist. The reference dataset consists of mouse whole-brain 10x single-cell data, the query set comprises MERFISH cells derived from 59 coronal sections. **i,** Visualization of the 5 (total 59) mouse whole-brain sections, with cells colored by predictions of CellMemory. **j,** Spatial coordinates of mouse brain MERFISH section 36, colored by CellMemory predictions. **k,** Heatmap displaying the memory scores of TAGs for all subclasses of IT-ET Glut in section 36. **l,** Distribution of subclasses and their corresponding TAGs’ memory scores within the spatial coordinates of section 36.

Ductal carcinoma in situ (DCIS) is a non-obligate precursor of invasive ductal carcinoma^47^. Characterization of key markers at invasive boundaries is crucial for understanding the molecular features associated with cancer heterogeneity. The inherent complexities and similarities among cells within DCIS regions pose challenges in achieving accurate interpretation. To overcome this, we utilized CellMemory to model a single-cell breast cancer dataset^48^ and integrate subcellular resolution breast cancer data from 10x Xenium^47^ (Supplementary Fig. 9). The visualization of the memory score demonstrated CellMemory’s interpretable representation of cell groups (Supplementary Fig. 9c, e). In a particular section of breast cancer tissue, CellMemory identified cancer-associated epithelial cells (Epi- Cancer) that different from the inner layer of ductal cells. Interestingly, two ductal regions showed distinct cellular compositions (Supplementary Fig. 9d). The first region (DCIS 1) contained a greater proportion of normal epithelial cells (Epi-Normal), while the second region (DCIS 2) had significantly fewer of these cells. The spatial distribution of Top Attention Genes by memory score (TAGs) like *KRT14*^49^ and *SERHL2*, which are used to identify Epi-Normal and Epi-Cancer cells respectively, exhibited variations (Supplementary Fig. 9d). Previous reports have indicated the presence of *KRT15*^+^ myoepithelial cells around DCIS, which are notably absent in highly invasive tumor regions. The absence of these cells is linked to a poor prognosis^50^. When CellMemory assigned weights to *KRT15* in spatial cells, a significant reduction of *KRT15*^+^ epithelial cells in DCIS 2 was observed (Supplementary Fig. 9f- g), indicating a higher degree of tumor invasion in this region. The confirmation of previous identifications of invasive tumor cells validated the insights provided by CellMemory. By providing spatial cell characterization and identifying critical TAGs associated with invasion, CellMemory contributes to an interpretable inference of cancer biology, thereby enabling the dissection of tumor ecosystems in spatial cancer data.

### CellMemory deciphers single-cell spatial data at the granule level

Characterizing single-cell spatial transcriptomics data at the granule level remains underexplored. We sought to verify whether CellMemory can annotate and integrate single-cell spatial omics data at this fine level of detail. We used human cortex data^51^ generated by Slide-tags as an example. Initially, CellMemory utilized a single-cell cortex dataset^32^ from normal samples to build a model at the L2 level (24 cell types). As illustrated in Fig. 3c, CellMemory produced appropriate embeddings and advanced annotations for the Slide-tags data, showcasing a harmonious integration of two distinct omics datasets (Supplementary Fig. 10a). Moreover, the memory score of TAGs generated by CellMemory offered valuable insights into the decision-making process for integrating spatial omics (Supplementary Fig. 10b). Subsequently, a model was constructed using granule-level cell information (L3, 131 cell types) and integrating single-cell and Slide-tags data using CLS embedding (Fig. 3d, Supplementary Fig. 11).

To assess the performance of integrating spatial data and the reliability of its label assignments, we incorporated Slide-tags information, and L1/2/3 annotations (Fig. 3e). The harmonious integration of data and coherent hierarchical annotations demonstrate that CellMemory can achieve granule-level characterization of single-cell spatial data. The effectiveness of the integration is visually represented in the Sankey diagram, which depicts a transition from coarse to granular categories (Supplementary Fig. 12). The specific expression of *LHFPL3* in oligodendrocyte precursor cells (OPC) is well-known^52^, and CellMemory appropriately assigned greater weight to *LHFPL3* in OPC_2. While *VWC2L* is expressed in several clusters, CellMemory uniquely assigns it higher weights in L4 IT_2 cells after considering all gene correlations. Remarkably, CellMemory can accurately characterize rare cell states, such as Micro-PVM_1 (*F13A1*^+^), which exhibit a very low proportion in the reference set (only 0.43‰) and in the spatial data (3.4‰), providing reasonable explanations for the integration result (Fig. 3f, Supplementary Fig. 13-14). In addition, CellMemory’s performance has been validated using Stereo-seq spatial data from mouse hemi-brain^53, 54^, achieving biologically meaningful interpretations (Supplementary Fig. 15).

The mammalian whole brain is incredibly complex. Understanding the cellular distribution within its spatial coordinates is crucial for comprehending consciousness, memory, emotion, and disease. CellMemory was trained using 781 scRNA-seq libraries, encompassing 4 million single-cell transcriptomes from the mouse brain, to analyze the Allen Institute for Brain Science (AIBS) MERFISH dataset^55^. This dataset includes 59 serial full coronal sections at 200-µm intervals spanning the entire mouse brain, totaling around 4 million cells (Fig. 3g-I, Supplementary Fig. 16). In a scenario with up to 338 subclasses in cross-omics representation, CellMemory outperformed pre-trained LLMs in generalization (Fig. 3h). Compared to a well-known annotation method, CellMemory demonstrated up to 15% improved representation accuracy. This implies that CellMemory accomplishes granule-level knowledge transfer in a single step, unlike previous methods that relied on hierarchical annotation processes. As an example, we present CellMemory’s interpretation of the IT-ET Glut neuronal subclasses in section 36 (Fig. 3j-k). The memory scores of TAGs displayed spatial distributions highly convergent with cell types, partially elucidating the logic behind CellMemory’s representation of spatial omics (Fig. 3l).

Our validation with Slide-tags, Stereo-seq, and MERFISH data conclusively demonstrated that CellMemory delivers ultra-high resolution in the characterization of single-cell spatial omics. This is essential for dissecting spatial cellular heterogeneity in complex biological systems and will significantly enhance our perception of spatial cell states.

### Integrating the Asian Immune Diversity Atlas (AIDA) with hierarchical interpretation

Integrating large-scale single-cell data across diverse populations and technological platforms presents a challenge, often resulting in the loss of low-abundance cells or accurate information due to batch effects and noise. We used PBMC CITE-seq data^7^ to construct a model that integrates AIDA data from 10x 5’ sequencing. The AIDA datasets include 503 healthy donors from five Asian ancestries across Japan, Singapore, and South Korea, as well as European controls (Fig. 4a-b). The cell embeddings generated by CellMemory demonstrated harmonious integration and high consistency with the original manually curated annotations (Supplementary Fig. 17). Notably, CellMemory identified a rare cell type called ASDC, which was not previously recognized in the AIDA dataset and constitutes only about 0.1‰ of the total cell population (Fig. 4b). The distinct expression profiles of *AXL* and *PPP1R14A* confirmed the accuracy of the ASDC characterization (Supplementary Fig. 18).

**Fig 4.**
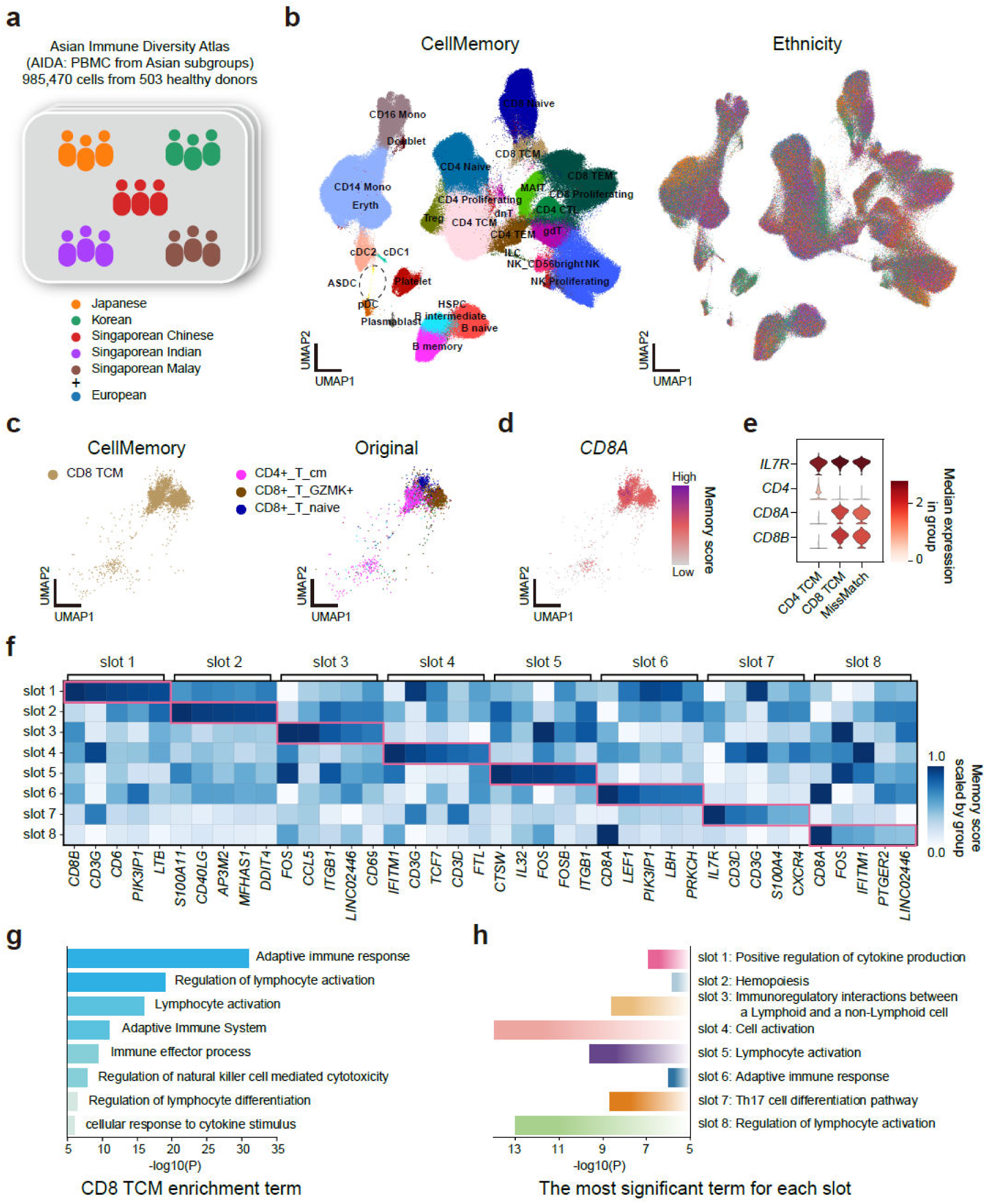
Integration of the Asian Immune Diversity Atlas with hierarchical interpretation. **a,** The Asian Immune Diversity Atlas (AIDA) comprises 985,470 cells from 503 healthy donors. The samples are obtained from Korea, Japan, and Singapore, including five Asian ethnicities, and five European donors as a control group. **b,** The integration of CellMemory with the AIDA dataset generated cell embeddings, which are colored by cell type annotations (left) from CellMemory and ethnicity (right) (randomly selecting 30 donors from each Asian population for fair observation of cell composition and state). **c,** Embeddings of CD8 TCM cells, annotated with CellMemory and original annotations. **d,** Memory scores of TAGs (CD8A) for CD8 TCM cells are displayed. **e,** The expression of IL7R, CD4, and CD8A/B in CD4 TCM, CD8 TCM, and the MissMatch group. **f,** Memory scores for the top five genes of each memory slot in the CD8 TCM group. **g,** Enrichment terms for the top 1-100 TAGs of CD8 TCM. **h,** For the CD8 TCM, the most significant terms in each memory slot (enrichment results for the top 1-50 gene sets in each memory slot).

We have identified a subgroup within CD8 TCM cells through CellMemory’s annotation that was originally mislabeled as CD4+_T_cm cells (Fig. 4c). Upon examining the memory scores of TAGs (*CD8A*) (Fig. 4d) and marker expression for both CD8 and CD4 T cells, we noticed that this Mismatch subgroup closely resembled the CD8 TCM cell phenotype (Fig. 4e). This observation underscores the complexity involved in characterizing immune cells in large-scale, multi-population studies and highlights the pivotal role of CellMemory’ interpretable integration.

We investigated how CellMemory makes decisions, particularly focusing on the hard-to-define CD8 TCM cells. Our research examined how each memory slot processes information within the memory space (Methods). Each slot assigns distinct weights to specific features, as illuminated by the memory scores of TAGs across all slots (Fig. 4f). And the most significant enrichment terms within each slot almost exclusively match a subset of the enrichment results of the cell type specific TAGs (Fig. 4g- h, Supplementary Fig. 19-20). For instance, slot 8 prioritizes the term “Regulation of lymphocyte activation”, while slot 6 focuses on “Adaptive immune response”- both of which are prominent terms in the CD8 TCM TAGs enrichment results. This discovery indicates that within CellMemory, when multiple specialist modules (gene programs) require coordinated management, they are organized into specific memory slots based on identifiable patterns, such as biological functions (Supplementary Fig. 1c). Consequently, each memory slot becomes specialized in comprehending and processing a distinct aspect of information, facilitating efficient collaborative processing. This mechanism is analogous to the cognitive process of driving, wherein the driver must synthesize and decipher a multitude of informational cues, such as road markings, traffic signs, and auditory cues, to navigate effectively. The bottlenecked architecture of CellMemory substantially enhances our understanding of the model’s dynamic approach to processing biological information, imparting an additional dimension of interpretability to its role in orchestrating complex biological processes.

### Studying malignant cells and the heterogeneity among patients using normal cells as a baseline

While it’s feasible to map data sets from healthy individuals to healthy reference atlases, interpreting diseased samples using healthy atlases is highly desirable but comes with distinct challenges^56^. Malignant cells are notably intricate, exhibiting aneuploidy and substantial heterogeneity among patients, thereby amplifying the inherent challenges of the task at hand. With CellMemory, we address this issue by utilizing its strong generalization capability, which allows the characterization of malignant cell states using an established healthy reference.

We focus on Mixed Phenotype Acute Leukemia (MPAL), a high-risk subtype of leukemia characterized by the co-expression of mixed myeloid and lymphoid features. Recent reports have indicated that this high-risk subtype is associated with abnormalities in primitive hematopoietic progenitors^57^. Accurately characterizing the malignant cell components and molecular signatures in MPAL patients is crucial for studying the potential mechanisms driving this complex disease. In our investigation, we trained CellMemory using bone marrow and peripheral blood mononuclear cells (BMMC, PBMC) from healthy individuals. We then applied this model to analyze malignant cells from two MPAL patients^58^, one with B myeloid MPAL (MPAL-a) and the other with T-myeloid MPAL (MPAL-b) (Fig. 5a-b). CellMemory was utilized to define the lineage proportions in these patients (Methods), identifying the progenitor-like state as the predominant original type (Fig. 5c).

**Fig 5.**
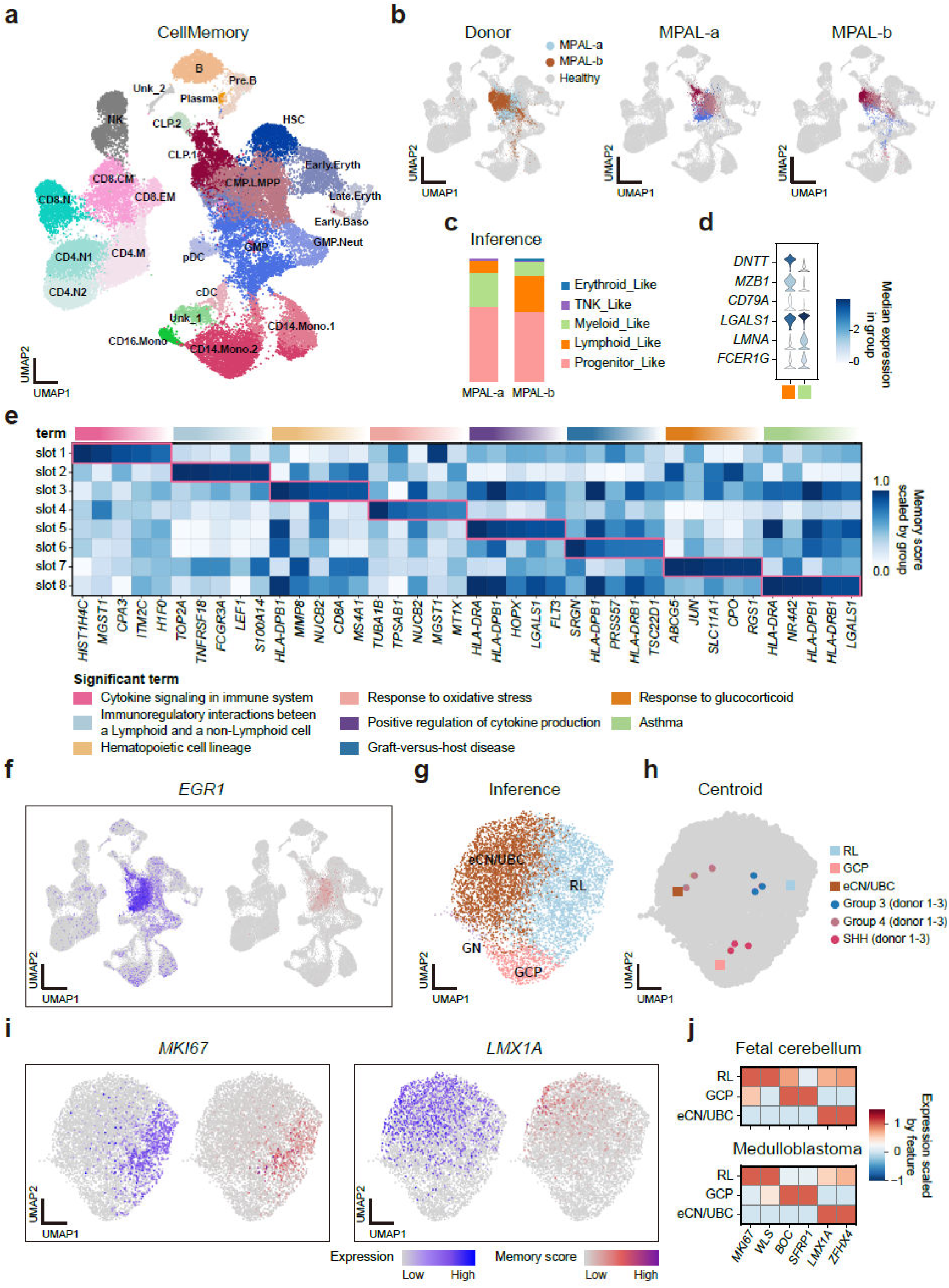
CellMemory interprets the individual heterogeneity of complex diseases. **a,** CellMemory integrated malignant blood cells from MPAL patients with bone marrow and peripheral blood mononuclear cells from healthy samples. **b,** Colored points show the positions of MPAL patient cells (MPAL-a: cells; MPAL-b: cells) in the co-embedding space and their inferred cell identities. **c,** Classify the lineage proportions based on cell types. **d,** The expression of lineage-specific genes in lymphoid and myeloid cells is inferred from MPAL-a patient cells. **e,** The heatmap showing the attention level of progenitor-like cells from MPAL-a patient in memory space. Eight memory slots are displayed, each showing the top five TAGs that it focuses on. The top 1-50 TAGs for each slot were subjected to enrichment analysis, with significant terms displayed above the heatmap. **f,** Expression and memory score of EGR1 in healthy cells and MPAL-a cells. **g,** Cell embedding generated by CellMemory for MB patients, colored by the inferred potential origin types of MB cells. **h,** The centroids of RL, GCP, and eCN/UBC cell populations are compared to centroids of cells from three subgroups of MB patients (each subgroup displaying three patients). **i,** UMAP shows the expression and memory scores of MKI67 and LMX1A. **j,** Heatmap displaying the expression of cell type specific genes in fetal cerebellum cells and malignant cells from MB patients.

After examining the specific features of lymphoid and myeloid cells in MPAL-a, we consistently observed characteristics related to the normal lineage and developmental stage of these malignant cells (Fig. 5d). The gene program level and gene level information stored in the bottlenecked memory space provide biologically meaningful interpretations for these malignant cells with progenitor-like properties. The GO term “Hematopoietic cell lineage” enriched in memory slot 3’s top attention genes suggests the cellular origin, while the term “Response to oxidative stress” in slot 4 reflects the cells’ stress state^59^. The term in slot 6 indicates immune function dysregulation^60^ (Fig. 5e, Supplementary Fig. 21a). *EGR1*, a key feature of the progenitor lineage^61^, was identified as a TAG by CellMemory, highlighting the origin characteristics of MPAL (Fig. 5f). Furthermore, CellMemory provided distinct hierarchical interpretations for MPAL-b, with increased attention to *DNTT*, implying potential differences in lineage or stages for the cell of origin among patients^62^ (Supplementary Fig. 21b-e).

Medulloblastoma (MB) is a malignant pediatric cerebellar tumor that can be molecularly divided into distinct subgroups^63^: WNT, SHH, Group 3, and Group 4. It is crucial to accurately identify the origins of each subgroup to better understand the disease and develop interventions. While it is currently believed that WNT MB originates from mossy fiber neurons in the embryonic dorsal brainstem^64^, the origins of Group 3 and Group 4 MB are still unclear. A recent report has suggested a potential association between the development of embryonic cerebellar glutamatergic neuronal lineages and MB^65^. To explore this, we employed single-cell data from SPLIT-seq^66^, which was generated from developing human embryonic cerebellar glutamatergic neuronal lineages, to construct a model. This model was then used to characterize individuals from various MB subgroups (SHH, Group 3, Group 4) based on data obtained through the Smart-seq2^67^ method, aiming to uncover the diverse origins of MB.

The findings indicate that cell types in MB can be confidently categorized into three lineages: eCN/UBC (excitatory cerebellar interneuron/unipolar brush cell), RL (rhombic lip), and GCP (granule cell progenitor) (Fig. 5f). The centroids of these cell identities (Methods) demonstrated high consistency with the centroids of MB subgroup donor cells (Fig. 5g, Supplementary Fig. 22). It is strongly suggested that Group 3 MB may originate from RL, Group 4 from eCN/UBC, and SHH from GCP. The distribution of memory score of *MKI67* in cell embeddings shows unquestionable similarities between MB cells and RL representations during embryonic cerebellar development^65^, as seen with *LMX1A* (Fig. 5h). The expression of markers for RL, GCP and UBC in MB cells, along with their memory scores, solidly supports the potential origins of different MB subgroups, in agreement with previous reports^65^ (Fig. 5i, Supplementary Fig. 22).

In conclusion, the precise characterization of OOD cells with CellMemory reveals the diverse nature of complex tumor origins. These clear and interpretable representations form a robust foundation for understanding the relationship between normal and disease states, underscoring the potential of CellMemory as an invaluable tool for identifying tumor founder cells.

### Investigating the intermediate stages in the development of lung cancer

Cancer begins with disrupted normal cells and genetic changes allowing tumor cell proliferation^68^. However, accurately annotating malignant cells via label transfer is challenging for computational methods due to the similarities between tumor cells and adjacent normal cells, as well as the considerable heterogeneity of tumors among different individuals. In this study, we aim to demonstrate the effectiveness of CellMemory by constructing a model using a lung cancer atlas^33^. The atlas includes tumor cells (Methods) as a reference, and we use another dataset consisting of 293,432 cells from 52 lung cancer patients^69^ as a query. This query dataset covers lung adenocarcinoma and lung squamous cell carcinoma, with a focus on LUAD results.

First, the capacity of CellMemory and scPoli to identify malignant cells was evaluated using the validation set. In this highly error-intolerant task, CellMemory exhibited robust performance across various tokenization strategies (Fig. 6a) (Methods). The trained CellMemory was used to integrate data from LUAD patients, resulting in the annotation of 24 cell types, including tumor cells (Fig. 6b, Supplementary Fig. 23a-b). The cell embedding revealed that transitional cells (*SCGB3A2*) within the epithelial cell population are most closely related to tumor cells. A subset of AT2 cells was found to be embedded within transitional cells (Fig. 6b), suggesting a similarity between AT2 and transitional cells. These findings support the reports that LUAD may exploit lung plasticity and regeneration mechanisms to generate tumors^69–71^. We named the AT2 cell embedded within transitional cells as AT2-2 (Methods), while those positioned further away were designated as AT2-1 (Fig. 6c, Supplementary Fig. 23c). The analysis of intercellular interactions^72^ indicates that tumor cells exhibit more significant interactions with cells like macrophages compared to transitional cells. The copy number variation (CNV) results^73^ and hierarchical interpretation of tumor cells further emphasize their malignant nature (Supplementary Fig. 23-24). This implies that the tumor cells identified by CellMemory display distinct characteristics associated with malignancy^74^.

**Fig 6.**
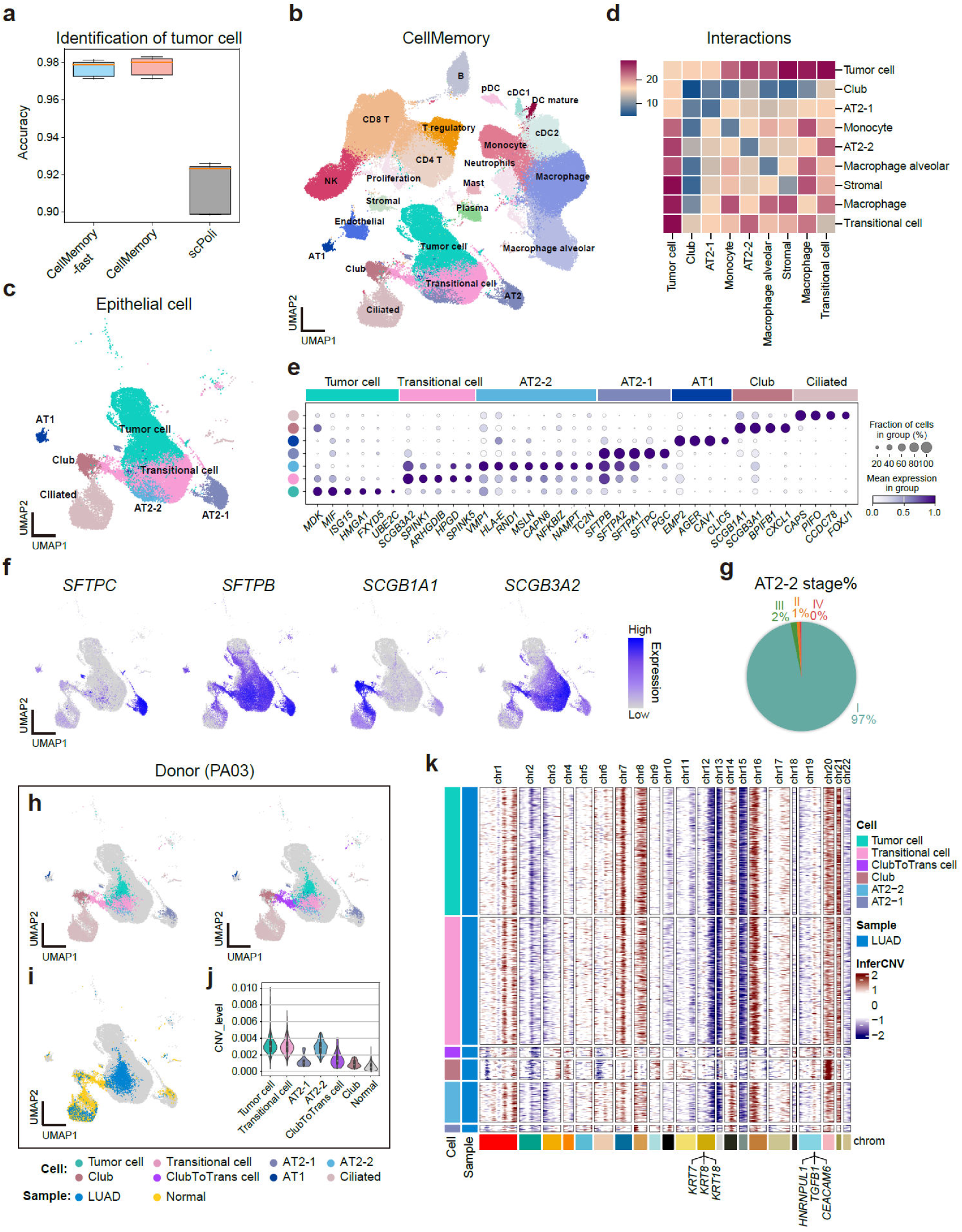
CellMemory accurately characterizes the lung cancer atlas. **a,** The accuracy of malignant cell identification by CellMemory and scPoli was validated through five-fold cross-validation. Filter: filter zero expression genes in cells. Full: consider all input genes. **b,** UMAP generated by CellMemory, integrating and annotating 149,595 cells from LUAD donors. **c,** UMAP of all epithelial cells in the LUAD dataset. **d,** Interactions between tumor-associated epithelial cells and monocytes, macrophages, stromal cells, etc. **e,** Dop plot demonstrating the specific gene expression in epithelial cells. **f,** UMAP displaying the expression of *SFTPC*, *SFTPB*, *SCGB1A1*, and *SCGB3A2* in epithelial cells. **g,** Proportion of stages identified as AT2-2 cells. **h-i,** UMAP displays epithelial cells from PA03, with color indicating cell type (**h**) and sample tissue (**i**). **j,** CNV levels of tumor-associated epithelial cells in PA03. **k,** CNV of tumor-associated epithelial cells in PA03 inferred using inferCNV. The CNV distances between each cell type were generated by inferCNV, which corresponds to the alignment of rows. Tumor cells and Transitional cells are down-sampled to 400 cells.

AT2-2 cells, when compared to AT2-1 cells, show closer associations with tumor cells (Fig. 6d). The specific gene expression trends of AT2-2 cells were found to be between those of AT2-1 and transitional cells. AT2-2 cells lose characteristics associated with AT2-1 (*SFTPC, SFTPA1/2*) while acquiring features typically associated with transitional cells (*SCGB3A2, SPINK*) (Fig. 6e-f, Supplementary Fig. 23d-f). Upon reviewing the interpretation in the memory space, we noticed that in memory slot 2, the gene *SFTPB* and the term “Surfactant metabolism” reflected the identity of AT2 cells, while in slots 1, 7, and 8, the genes and terms indicate that AT2-2 cells are acquiring tumor-related cell states (Supplementary Fig. 24c-d). Furthermore, the interactions, stages, and CNV patterns of these cells suggest that AT2-2 cells represent an intermediate state in the progression of LUAD from AT2 cells (Fig. 6d, g, Supplementary Fig. 23g-i). Genes specifically expressed in AT2-2, including *MSLN*, *CAPN8*, *NAMPT*, and *TC2N*, have been identified as potential therapeutic targets, with the knockout of these genes significantly suppressing LUAD^75–78^. *HLA-E,* known in pancreatic cancer to be exploited by circulating tumor cells (CTCs) to evade immune checkpoints and promote metastasis^79^, is closely related to the most focused GO term “Antigen processing and presentation of exogenous peptide antigen” in memory slot 1 of AT2-2 (Supplementary Fig. 24d). This implies that the AT2-2 state plays a pivotal role in cancer progression and may be a valuable target for therapeutic intervention.

CellMemory confidently creates cell embeddings that thoroughly consider all input gene interactions without being constrained by prior information. This effectively characterizes the cell states in an understandable way for seamless integration and offers a robust method for exploring the continuum of cell states within tumor contexts.

### Dissecting the heterogeneity of founder cells among lung cancer patients

Single-cell transcriptomics enables a detailed understanding of patient-specific disease states. In the LUAD dataset, the cell embeddings generated by CellMemory revealed the similarity between AT2 cells and tumor cells. For example, in patient PA11, we observed that the cell representations and CNV patterns of AT2 cells closely resembled those of tumor cells (Supplementary Fig. 25), consistent with the common view of founder cells in LUAD. Additionally, the cell embeddings indicated that club cells and their neighboring transitional cells follow a trajectory that converges toward tumor cells. In a subset of patients, tumor tissues displayed CNV patterns in club cells that were similar to those in tumor cells (Supplementary Fig. 23g). We are providing personalized characterization for patient PA03. (Fig. 6h-i, Supplementary Fig. 26). The embedding of epithelial cells in PA03 revealed a distinct group that had converted from club cells to transitional cells. Previous reports have indicated that bronchial cells can use intermediate state cells with differentiation potential to replenish alveolar cells^80^. Accordingly, we have designated this cluster of cells as ClubtoTrans cells (*SCGB1A1, SCGB3A2, SFTPB*) (Methods).

In the tumor tissue of the PA03 patient, AT2-2 cells exhibited CNV levels and patterns akin to those observed in tumor cells (Fig. 6j-k). Interestingly, a small number of club and ClubtoTrans cells were identified within the tumor tissue of the PA03. The CNV pattern observed in tumor cells was also evident within ClubtoTrans cells. Based on the clustering results from InferCNV, the CNV profile of ClubtoTrans cells is more closely aligned with that of tumor cells (Fig. 6k). In key oncogenic regions of LUAD (e.g., *KRT8, CEACAM6*), ClubtoTrans cells exhibit significant CNV amplification (Fig. 6k). Recent reports have pointed out that club cells can develop tumors by leveraging the plasticity and regenerative capacity of the lung^70^. During the transition, club cells acquire an AT2-like epigenetic pattern, which is consistent with CellMemory’s characterization of the single-cell transcriptome (Fig. 6h). Importantly, the heterogeneity of founder cells among LUAD patients is closely associated with tumor state transdifferentiation^81^. In the club and ClubToTrans cells of patient PA03, the expression of LUSC markers was observed (Supplementary Fig. 26e-27). This suggests that the patient may be in the early stages of transitioning to lung adenosquamous carcinoma (LUAS)^81^. However, the potential impact of this transition on drug resistance and prognosis was not recognized in the original study. The integration results generated by CellMemory revealed similar state changes in another cohort of LUAD data for certain donors (LUAD dataset 2, Methods) (Supplementary Fig. 28-30).

It can be challenging to focus on rare subpopulations when integrating population-scale data, due to confounding factors such as background noise and batch effects. However, CellMemory’s analysis of lung cancer data shows that it can effectively transfer important prior knowledge without bias (Supplementary Fig. 31-33). Identifying potential founder cells specific to patients will provide valuable insights into understanding individual drug resistance and personalized targeted interventions for tumor progression.

## Discussion

To take advantage of existing reference data for understanding out-of-distribution (OOD) cells^5^, we have developed CellMemory. CellMemory is a Transformer model that employs a cross-attention mechanism inspired by a theory of consciousness. We conducted a comprehensive comparison of 18 task-specific methods for integration and annotation data, utilizing data from over 15 million cells. Due to its generalized representations, CellMemory outperforms other methods including scGPT and Geneformer^23^, especially in cross-omics situations. We believe that CellMemory’s exceptional performance is due to its bottlenecked architecture’s attention and competitive mechanism, which allows it to retain essential information^20^.

We have studied different strategies for processing tokens in single-cell data and observed that they have varying impacts on data integration. When integrating datasets with technical variability, such as single-cell and single-cell spatial data, it is recommended to scale gene token values into bin ranges. Current self-attention-based single-cell LLMs often exclude zero-value genes due to computational cost constraints^16, 17^. However, our evaluation revealed that keeping zero-value genes in cells resulted in enhanced integration performance. This highlights the importance of devising efficient model architectures for analyzing of single-cell omics.

We then focused on scenarios involving cross-omics, ethnic populations, and the comparison of disease versus healthy states to explore interpretable inference for OOD cells. Notably, by analyzing the most focused genes in cell type specific memory slots, we identified significant biological function differences within the gene programs that are targeted by different memory slots. A variety of specialist modules were organized into distinct shared global workspaces within a bottleneck, ensuring consistency in task completion. This process is analogous to the brain coordinating and unifying driving-related visual and auditory information in the consciousness space for safe driving. This hierarchical interpretation facilitates the analysis of OOD objects related to cellular states, further enhancing our understanding of the dynamic manner in which the model processes biological information.

The identification of malignant cells and the characterization of disease states have posed significant challenges in analyzing single-cell datasets. Previous studies have shown that accurately characterizing disease state cells typically requires the inclusion of control groups (usually derived from healthy samples) during model training^56^. CellMemory, without relying on control samples, exhibits exceptional accuracy in distinguishing between normal and malignant cells in lung cancer. It can illustrate the cell representations of disease samples, investigate the intermediate states of LUAD origin, and reveal the heterogeneous founder cells in some individuals. This emphasizes the crucial role of employing a model capable of discerning the finest details with utmost clarity to universally interpret disease states. By assisting researchers in delineating the broader context alongside intricate specifics, CellMemory facilitates the practice of precision medicine at an individualized level.

As advancements in single-cell references continue, CellMemory is anticipated to fulfill a pivotal role in facilitating reference mapping without the need for repeated training. This will enable a deeper understanding of cell representations beyond the current paradigms. Moreover, its bottlenecked architecture is optimally suited for integrating multimodal data, thus enabling comprehensive information exchange within the low-dimensional memory space. Its capability for processing long tokens makes it particularly suitable for sparse, high-dimensional single-cell omics data and reduces pre-training expenses. The objective of the forthcoming phase is to extend the applications of CellMemory in these areas.

## Methods

### The Global Workspace Theory

Our project builds on the Global Workspace Theory (GWT) in neuroscience, which suggests that consciousness arises from the selective dissemination of information through a shared "global workspace"^19–21^.

The brain consists of numerous specialized modules and processes that handle various tasks, such as vision, hearing, memory, and emotions. The "global workspace", a network of densely interconnected neurons, enables these specialized modules to share and communicate information. However, not all pieces of information can reach the global workspace due to its limited capacity. Instead, the specialized modules may compete with each other to write information to the global workspace. When certain information becomes dominant in the global workspace, it enters our conscious awareness, while other, non-dominant information remains in the unconscious realm. Once information occupies the global workspace, it is broadcasted to diverse specialized neural modules through synchronized neural firing and rapid signaling. This broadcast mechanism ensures that various brain regions concurrently process the information, contributing to a unified conscious experience.

Recently, GWT was used to model complex environments in deep learning^22^. Their approach leveraged the shared global workspace as a communication bottleneck among specialists, significantly outperforming the conventional self-attention frameworks across various visual reasoning benchmarks. Inspired by the GWT in neuroscience and the promising results in deep learning, our project tailored GWT for analyzing single-cell omics data. Single-cell data presents a unique challenge due to its notoriously high dimensionality and intricate underlying patterns. We found that the global workspace model is ideally suited to address this issue. Its bottleneck architecture acts as a filter, prioritizing and highlighting the most significant information, ensuring that the most pertinent biological details are efficiently communicated and processed. Furthermore, the conventional self-attention mechanism used by BERT presents a quadratic complexity concerning sequence length, leading to increasing time and memory complexity as sequences expand^12, 13, 24^. This becomes drastically problematic when dealing with lengthy single-cell omics data. Alternatively, we employed cross-attention to implement the shared global workspace, substantially reducing time and space complexity and resulting in a more efficient single-cell omics data analysis.

### The methodology of CellMemory

Assuming we have an input matrix ***X*** ∈ ℝ*^N^* ^×^ *^M^*, where *N* represents the number of cells and *M* stands for the total number of genes.

#### Embedding

During the embedding process, we encoded each gene of each cell using a word embedding strategy. The processed expression values were transformed into discrete values through a bag-of-words approach (rounding up, or scaling to bin ranges), generating token embeddings. The positions of genes were utilized to create position embeddings.

#### Construct specialist matrix

For each cell, *n* ∈ *N* of the input matrix **X**, an extra CLS token is appended to the front of genes. This extended representation is further converted into a *k*-dimensional embedding space, resulting in the specialist matrix **R** ∈ ℝ*^N^* ^×^ ^(1+*M*)^ ^×^ *^k^*, where *K* is a hyperparameter (please refer to “Hyperparameter settings” for details) identifying the number of dimensions essential for processing complex data relationships, and *M* represents the number of specialists (genes).

#### Compete to write

The specialist modules **R** compete to write the information to the shared global workspace (Supplementary Fig. 1a), resulting in the memory matrix **D** ∈ ℝ^N^ ^×^ ^H^ ^×^ ^K^ , with *H* distinct memory slots serving as a hyperparameter. Typically, *H* is much smaller than the original input dimension *M*, reflecting the bottleneck of the global workspace. The writing procedure involves a cross-attention mechanism between the specialist matrix **R** and the memory matrix **D**, detailed as follows.

(1) Initially, we construct the Query, Key, and Value matrices. Here, 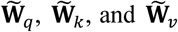 represent the respective weight matrices for the Query, Key, and Value when calculating the cross-attention matrix.

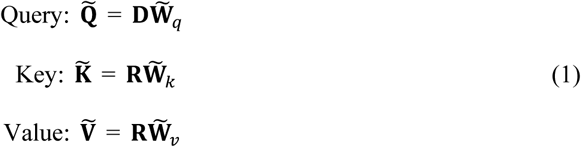

(2) Then, the attention matrix A is calculated by multiplying the Query and Key matrices, scaled by the last dimension of attention weights d, and applying a softmax function for normalization. We select the top k largest attention scores along the 1 + *M* dimension, where specialized modules compete to write relevant information to the global workspace. This algorithm aligns with the principles postulated in the global workspace theory in neuroscience, wherein various cognitive processes compete to access a centralized, conscious processing stage. It’s worth noting that, in our experiments, we observed a slight performance improvement when using the top *k* mechanism, but it impacted the interpretability of the model results. Therefore, in this article application, we did not employ the top *k*. If the focus is solely on cell type prediction and the input tokens are relatively long, we recommend considering the use of the top *k*.

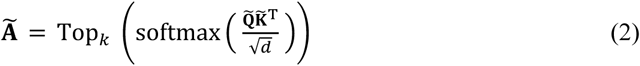

(3) Finally, the attention matrix A is multiplied by the Value matrix V , resulting in the updated memory **D**.

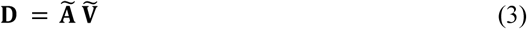

#### Broadcast to specialists

The memory matrix **D** is broadcasted back to the specialists (Supplementary Fig. 1b), leading to the creation of an updated specialist matrix **R**. This broadcasting action also utilizes a cross-attention mechanism, which is demonstrated as follows.

(1) Similar to the writing mechanism, we formulate the Query, Key, and Value matrices when computing the cross-attention for broadcasting. Specifically, the weight matrices associated with the Query, Key, and Value are represented by 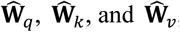, respectively.

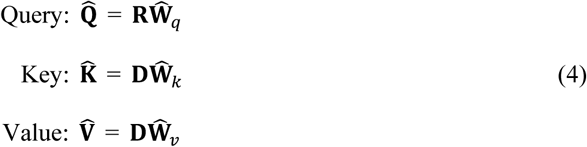

(2) Then, we calculate the attention matrix A by applying softmax over the slots.

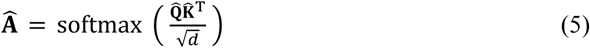

(3) Eventually, we construct the updated specialist matrix R.

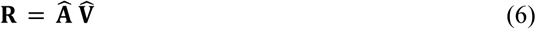

The processes of "Compete to Write" and "Broadcast to Specialists" are not confined to singular iterations. Instead, they can be orchestrated in a recurrent sequence: write → broadcast → write → broadcast, and so forth. By repeatedly updating the global workspace and broadcasting for specialized feedback, the system ensures a comprehensive integration of information, as each iteration enhances the clarity and resolution of the analysis. Therefore, when applied to the single-cell omics data, this repeated cycle promotes a thorough understanding and in-depth analysis of the data.

#### Model training and prediction

In the training process of CellMemory, the CLS token matrix **S** ∈ *ℝ^N^* ^×^ ^k^ is extracted from the first dimension of the revised specialist matrix **R** at index S_n,k_ = R*_n,o,k_* Through the iterative process of writing and broadcasting, the CLS token acquires substantial intracellular information. The CLS token **S** is then projected through a fully connected layer *f* and processed to the probability for each class *c* ∈ **C** using softmax function. The predictions are compared to the true labels, and the cross-entropy loss L_CE_ is calculated as follows.

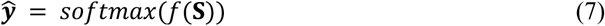

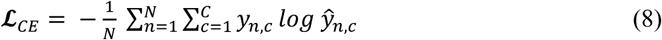

CellMemory is guided to focus on acquiring information related to the identity representation of cells. During the prediction, the softmax function assigns each cell’s prediction results to the probability of each category, with the category having the highest probability (or confidence score) being considered as the cell’s identity.

### Hyperparameter settings

The gene expression matrix, after undergoing embedding, is input into a 4-layered transformer along with the CLS token for classification. We set the embedding dimension (*K*) to 256 and the feed-forward network dimension to 512. The Query and Key sizes are set to 32, and the Value size is set to 64. We employ 4 heads during reading from and writing into the global workspace, which comprises 8 memory slots (*H*). We train the model using an early stopping strategy, stopping training when the validation set’s loss has not decreased for 5 epochs. We use Adam optimizer with a learning rate of 0.0003 and anneal the learning rate using cosine annealing^22^.

### Performing downstream tasks

(1) CLS token: The CLS token **S** ∈ ℝ^N^ ^×^ ^K^ effectively communicates with gene tokens inside the cell through attention mechanisms and the bottleneck architecture in a supervised manner, obtaining a complete cellular representation. It is employed for single-cell and single-cell spatial data reference mapping.

(2) Memory score matrix: The interpretability of CellMemory originates from the cross-attention mechanism’s explicit demonstration of the importance of input features. The gene expression matrix undergoes a writing and broadcasting process with the memory, leading to the generation of two attention score matrices 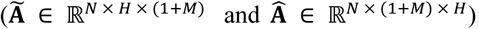 . Matrix multiplication is performed on the two attention matrices derived from the writing and broadcasting steps, yielding a resultant matrix **Z** = AA ∈ ℝ^N^ ^×^ ^(1+M)^ ^×^ ^(1+M)^. From matrix **Z**, we extract matrix 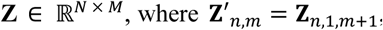 apturing the attention of each CLS token concerning each of the *M* genes across *N* cells. To generate cell-type-specific TAGs, the averaged memory score for specific groups of cells is computed based on the global attention score matrix **Z**. Then, the scores are sorted to obtain the top attention genes by memory score for each group. To generate memory slot specific TAGs, the matrix A produced during broadcasting is used. The mean is calculated across selected cells, then, based on averaged attention scores, the features are sorted to obtain TAGs for each memory slot.

### Theoretical analysis

CellMemory is more computationally efficient than the conventional self-attention mechanisms, especially in single-cell omics data analysis. Specifically, the traditional self-attention approach used in BERT is characterized by quadratic complexity as a function of sequence length. This inherent complexity becomes a substantial impediment as sequences extend, particularly when dealing with voluminous single-cell omics datasets. Using traditional self-attention for such extensive data requires substantial computational resources and elongated processing time, making it unsuitable for real-time applications. In contrast, CellMemory harnesses a cross-attention mechanism to implement the shared global workspace, significantly reducing time and space complexity.

### Time complexity

Our proposed method, CellMemory, outperforms traditional self-attention mechanisms in terms of computational time complexity. Consider the "Compete to Write" procedure. For each cell n ∈ N , we have Query 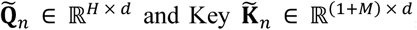, and the total time complexity to calculate cross-attention matrix 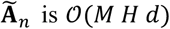 in our proposed method. Similarly, due to the consistent use of cross-attention, the time complexity to calculate 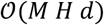 in the Broadcast to Specialists" procedure. In comparison, the traditional self-attention mechanism exhibits a total time complexity of 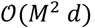 . Specifically, *H*, the hyperparameter representing memory slots, is typically chosen to be much smaller than the input dimension *M* to reflect the bottleneck of the shared global workspace.

### Space complexity

Furthermore, CellMemory also surpasses the traditional self-attention mechanisms regarding space complexity. For each cell n ∈ N, the "Compete to Write" procedure yields a cross- attention matrix A_n_ with dimensions (1+M) × H, while the "Broadcast to Specialists" procedure results in A_n_ with dimensions (1 + M) × H. Consequently, the space complexity of our method is O(MH). In contrast, the traditional self-attention mechanisms operate with a quadratic space complexity of O(M^2^). Given H ≪ M, it is evident that CellMemory offers a more memory-efficient alternative, saving valuable computational resources compared to the conventional self-attention techniques.

### Relation with other Transformer variants

Transformers, with their attention-based architecture, have become a cornerstone in deep learning. However, traditional transformers face challenges like quadratic time and memory complexity and a large parameter count, impeding scalability. To address these issues, numerous transformer variants have emerged in recent years. Our method aligns with variants incorporating a ’Neural Memory’ and implementing down-sampling within the attention mechanism^82^. This approach is directly inspired by the transformer shared global workspace^22^ and bears similarities to the Perceiver model^83^. Both strategies refine the pairwise self-attention mechanism by introducing global coordination and a unified memory. This integrated memory acts as a constrained communication channel, distinct from other transformer bottleneck techniques like discrete bottleneck^84^ and restricted cross-modal attention^85^. In our model, bottleneck effects are achieved through a small, integrated memory, facilitating information integration and dimension reduction. These features are particularly advantageous for processing complex and noisy biological data, as in our study.

### Single-cell dataset

#### hBreast

We collected a single cell atlas of the adult human breast from the CELLxGENE database. This dataset includes 244,285 cells (100,000 cells from normal samples by down-sampling; 144,285 cells from breast cancer samples), identifying 58 biological cell states. The data was subset to the 3,000 highly variable genes (HVGs) to benchmark. We established an annotation scenario using normal cells as the reference for annotating cells from breast cancer samples. The CLS embedding of the query set was evaluated in the integration benchmark. Batch information was denoted using the ‘library_uuid’.

#### hLung

The Lung atlas is available in the CELLxGENE database, comprising both normal and LUSC samples It contains 305,319 cells (212,889 cells from normal samples and 92,430 cells from LUSC samples) and identifies 24 cell types. We restricted the reference dataset to data exclusively from the 10x platform to predict the cell type of the query set of LUSC. The query set incorporated data from 5 platforms. In the integration benchmark, the term ‘dataset’ was utilized as the batch label. A total of 3,000 HVGs were used for benchmarking.

#### hPancreas

The human pancreas dataset includes 47,644 cells (26,197 cells from control samples; 21,447 cells from T1D patients, and 13 cell types). We removed the hybrid cell type from the dataset, and cells from control donors were used as the reference while cells from T1D patients were analyzed as the query set. A total of 3,000 HVGs were used for benchmarking.

#### Immune

The training set was composed of 141,524 immune cells derived from 8 tissues, including the lung and liver. The test set consisted of 188,238 immune cells from 9 tissues, such as the spleen and thymus. A total of 4,000 HVGs were used for benchmarking.

#### mPancreas

The mPancreas contains scRNA-seq data from 239,679 mouse pancreatic islet cells (with the reference down-sampled to 100,000). The normal cells within the dataset served as a reference to train the model, and the T1/2D samples were utilized as the query set (139,679 cells). This dataset comprises 20 cell types. The term ‘batch_integration’ was utilized as the batch label. A total of 3,000 HVGs were used for benchmarking.

#### mHypoMap

The mouse hypothalamic atlas from HypoMap contains 66 cell populations derived from the transcriptional profiling of 161,903 cells. We used the data from the Drop-seq platform as the query set (61,903 cells) and the data from the 10x platform as the train set (downsampled to 100,000 cells). The term ‘Sample_ID’ was utilized as the batch label. A total of 4,000 HVGs were used for benchmarking.

#### crossSpecies (cortex)

In the cross-species benchmark, we utilized human cerebral cortex data, comprising 200,000 cells, as the training set. The query set was constituted by a dataset that includes one human and four non-human cerebral cortex datasets (human: 156,285 cells, Gorilla: 139,945 cells; Chimpanzee: 112,929 cells; Macaque: 89,136 cells; Marmoset: 75,861 cells). This part ensured the retention of a set of homologous genes across all species. The performance of the integration was evaluated using chimpanzee cortex data. A total of 3,000 HVGs were used for benchmarking.

#### Drosophila

The Drosophila dataset (body) comprises 265,979 cells (17 cell types), categorized into four age groups (5, 30, 50, 70 days), aiming to investigate aging modifications in Drosophila at the single-cell level. The younger samples of 5 days and 30 days served as the reference (173,181 cells), and the older samples of 50 days and 70 days were used as the query (92,798 cells). In the benchmark comparison, we omitted Geneformer and scGPT from the Drosophila dataset due to substantial disparities between the gene features used for their pre-training and those of Drosophila’s gene features. A total of 3,000 HVGs were used for benchmarking.

#### Breast cancer atlas

Using the breast cancer single-cell data as a reference, we constructed a model to integrate the 10x Xenium data. We retained the overlapping gene set (301 genes) with the Xenium data and included cells with counts more than 20, and genes more than 10, totaling 94,535 cells. The model was built at the “celltype_major” resolution, integrating the Xenium and 10x 3’/5’ single-cell data (from the same sample). Gene expression values were scaled to the bin range.

#### Brain cell atlas of Alzheimer’s disease (SEA-AD)

This atlas was sampled from the dorsolateral prefrontal cortex of Alzheimer’s disease patients, comprising 1,395,601 cells. Among them, 796,768 cells originated from normal samples and 598,833 cells from disease samples. It includes annotations at both subclass (24) and supertype (131) resolutions. In the integration assessment, cells from normal samples were used to construct a reference, which was then integrated with cells from disease samples, while retaining the 3,000 highly variable genes.

#### The Asian Immune Diversity Atlas (AIDA)

AIDA is an immune dataset comprising 985,470 PBMC cells from 503 healthy donors across five Asian ancestries from Japan, Singapore, and South Korea, along with European controls (5 donors, 73,439 cells) via 10x 5’ single-cell sequencing.

#### Adolescent mouse nervous system

The single-cell mouse brain atlas of cell types from the Linnarsson Lab is employed as the reference dataset to characterize Stereo-seq adult mouse coronal hemibrain dataset. Cell types consistent with the organization of the query dataset were retained, and the model was trained using over 120 cell types. This work provided identified marker genes, which were used to train the model for reference mapping.

#### Mouse whole-brain scRNA-seq atlas

A scRNA-seq whole adult mouse brain dataset of 4,057,701 cells was generated by the 10x platform (v2, v3) with 338 subclasses. This dataset was used to build a model to infer mouse whole-brain MERFISH data. The gene set was intersected with MERFISH data to retain 500 genes, and cells with more than 30 captured genes were kept.

#### CITE-seq PBMC

We used PBMC cells produced by CITE-seq (161,764 cells, 36 classes) to integrate and annotate the AIDA dataset. A total of 4,000 highly variable genes were utilized.

#### MPAL lineage inference

Bone marrow and peripheral blood mononuclear cells from healthy samples were used, totaling 35,582 cells (MBBCs=12,602; CD34+- enriched BMMCs=8,176; PBMCs=14,804). A model was constructed using 1,000 highly variable genes retained from the reference. MPAL-a (3,435 cells) corresponds to B-myeloid MPAL (MPAL4) in the original study, and MPAL-b (2,947 cells) corresponds to T-myeloid MPAL (MPAL5). Cells labeled as healthy-like in the original study were excluded.

#### Origin inference for different MB subgroups

Glutamatergic neuronal lineages from the fetal cerebellum were used as the reference, with 500 highly variable genes. Malignant MB cells produced by Smart-seq2 (including SHH, Group 3, and Group 4 subgroups from 31 donors) were used as the query set.

#### Integration of lung cancer dataset

We used all healthy samples, LUSC samples, and LUAD samples (downsampled to 200k cells) from **hLung** dataset, and retained 3,000 highly variable genes. We then integrated a lung cancer dataset comprising 293,432 cells from 52 lung cancer patients (including LUAD and LUSC) using the trained CellMemory. For tumor cell similarity inference, all tumor cell types were removed from the reference, retaining 5,000 highly variable genes to train the model.

#### Integration of LUAD dataset 2

A model was constructed using the same reference as the **Integration of lung cancer dataset**. This model was then integrated into the LUAD dataset comprising 488,236 cells from 58 LUAD donors.

### Single-cell spatial dataset

#### mHypo

The mouse hypothalamic spatial dataset measured by MERFISH, comprising three samples with naïve behavior, totaling 180,754 cells. The single-cell data from **mHypoMap** was used as the reference. Aligning genes (154) were captured by MERFISH resulting in 8 categories.

#### hNSCLC

The scRNA-seq dataset of **hLung** served as the reference to predict NSCLS spatial dataset. The NSCLC spatial dataset includes Lung-6, Lung-9, and Lung-13, consisting of 256,581 cells from 11 cell types. These samples were generated from a Formalin-Fixed Paraffin-Embedded (FFPE) sample by the CosMx SMI platform. A shared gene set of 952 genes from the reference dataset was selected to align with the query dataset.

#### mSpermato

The spatial dataset of mouse spermatogenesis (measured by Slide-seq) was acquired from three wild-type (WT) mice and three leptin-deficient diabetic (ob/ob) mice, including 207,335 cells and 9 cell types. The WT dataset served as the reference, and the ob/ob dataset served as the query set.

#### Xenium Breast cancer

This dataset consists of two replicate samples from a breast cancer patient, as well as single-cell 3’ and 5’ data. Cells with counts of more than 20 and more than 10 genes were retained, totaling 305,631 cells and 301 genes.

#### Slide-tags human cortex

The spatial transcriptome data generated by Slide-tags sequencing consists of 4,065 cells belonging to 7 cell types. Cells from normal samples in the brain cell atlas of Alzheimer’s disease were utilized as a reference for integrating the Slide-tags spatial data.

#### Stereo-seq adult mouse coronal hemibrain

The single-cell spatial dataset of the mouse brain hemisphere (including 50,140 cells) was measured by Stereo-seq. We utilized segmented data as the query set. It maintained the subset of genes that align with the reference (**Adolescent mouse nervous system**).

#### Mouse whole-brain MERFISH atlas

A MERFISH dataset, encompassing 4,334,174 cells across 59 sections from one whole male mouse brain with a 500-gene panel. This dataset includes a hierarchical cell type annotation, organized into four levels of cellular identities. To evaluate the performance of benchmark models, we used the subclass information. Cells with more than 30 captured genes were kept.

### Comparison with annotation methods

CellTypist (1.5.3, python) emerged as an efficient annotation tool that outperforms most methods. Seurat (4.3.0.1, R) is a widely employed method in single-cell data analysis. In this study, we utilized the implementation of the cell type annotation pipeline as described in the “Mapping and annotating query datasets” chapter. TOSICA (1.0.0, python) is a self-attention Transformer-based model that computes attention at the pathway level and performs cell type annotation. scPoli (python) used the default set.

### Comparison with the pre-trained Transformer model

#### Geneformer

Geneformer is an LLM that leverages pretraining with a dataset of 30 million cells, allowing it to perform tasks such as cell type annotation and the inference of key network regulators. The cells used for pre-training are sourced from human data. Geneformer requires pre-trained weights as input.

#### scGPT

scGPT is an LLM that leverages pre-training with a dataset of 33 million cells. The cells used for pre-training are sourced from human data.

#### scBERT

scBERT adopts the advanced strategy derived from BERT, tailored to implement single-cell data analysis. scBERT is exclusively pre-trained on a human dataset (Panglao dataset, 1,126,580 cells), there are no pre-trained models available for other species.

### Comparison with spatial annotation methods

CellMemory seeks to label individual cells and cell-like objects, such as Stereo-seq spots or Slide-seq beads, through discrete annotations. To enhance the interpretability of spatial single-cell data, understanding the most critical features for each cell is essential. Compositional annotation refers to probabilities associated with discrete categories, indicating objects closely related to the assigned category (the class with the maximum compositions). Default parameters were utilized for all other spatial annotation methods (Tangram (“clusters” level), TACCO, RCTD, NMFreg, NNLS).

### Comparison with integration methods

In benchmark integration tasks, aside from scGPT and scPoli, all other integration methods were performed using scIB^36^. scGPT performs the integration process, and scPoli performs integration and annotation using default steps. The CellMemory CLS embedding used by the integration benchmark is the same as that used by the annotation. The original cell types are applied to biological conservation evaluation.

### Training strategy of the CellMemory

In the process of model training, the reference dataset is divided into a training set and a validation set with an 80:20 ratio. In the annotation benchmark validation, we employed a five-fold cross-validation to ensure the reliability of the results.

### Evaluation metrics

We employed accuracy and macro f1-score as evaluation metrics to assess the classification performance. These metrics were computed using the Python library sklearn. We assessed the data integration using metrics from scIB package. The overall integration score is a weighted average of the average batch mixing score and the average biological conservation score, with weights of 0.4 and 0.6 respectively.

### Enrichment analysis

The enrichment analysis was performed using the Metascape^86^ web-based platform. The analysis was conducted using the TAGs, with enriched terms including GO terms and pathways.

### Centroid determination for MB

When inferring the identity of origin cells, we first applied Leiden clustering to all MB cells, resulting in 13 clusters. We identified the cluster with the highest mean confidence score for each cell type, then averaged the embedding coordinates of all cells in that cluster to determine its centroid. The embedding coordinates of all cells from a specific donor were used to determine the donor centroid, while the embedding coordinates of all cells from a specific subgroup were used to determine the subgroup centroid.

### Identification of Tumor cells

A lung cancer atlas (including tumor cells) was employed as a reference to construct the model. The accuracy of tumor cell identification was validated using tumor cells from the validation set.

### Cell-cell interaction

CellPhoneDB^72^ was used to analyze the interactions between LUAD epithelial cells and other cells such as macrophages.

### CNV inference

inferCNV^73^ was employed to infer CNV patterns and levels in lung cancer patients.

### Definition of AT2 subgroups

The Leiden clustering algorithm was employed to analyze the cell embeddings generated by CellMemory. AT2 cells that overlapped with cluster 5 were designated as AT2-1, while AT2 cells that were not included in cluster 5 were labeled as AT2-2.

## Data availability

All single-cell and single-cell spatial omics data in this study were published previously, and their availability is described in Supplementary Table 2.

## Code availability

The software is available at https://github.com/QifeiWang4Bio/CellMemory.

## Supporting information

Hierarchical Interpretation of Out-of-Distribution Cells Using Bottlenecked Transformer-bioRxiv

## Acknowledgments

We appreciate the user-friendly data provided by the CELLxGENE database. L.J. was supported by the National Key Research and Development Program of China (2019YFA0801702), the Strategic Priority Research Program of the Chinese Academy of Sciences (XDB38020500), the National Natural Science Foundation of China (31970760), and the International Partnership Program of the Chinese Academy of Sciences (153F11KYSB20210006).

## Author contributions

D.L., L.J., and Q.W. designed the project. Q.W. and H.Z. built the model. Q.W., Y.H., and Y.C. conducted the model application, benchmark, and analysis. Q.W., L.J., D.L., Y.W., and H.Z. wrote the manuscript. Y.L., J.Z., X.Z., and M.K. provided comments and application suggestions for the project. All authors discussed the results and commented on the manuscript.

## Reference

1. Sikkema, L. et al. An integrated cell atlas of the lung in health and disease. Nature medicine 29, 1563–1577 (2023).

2. Jones, R.C. et al. The Tabula Sapiens: A multiple-organ, single-cell transcriptomic atlas of humans. *Science (New York*, N.Y.) 376, eabl4896 (2022).

3. Lander, E.S. et al. Initial sequencing and analysis of the human genome. Nature 409, 860–921 (2001).

4. Sandberg, R. Entering the era of single-cell transcriptomics in biology and medicine. Nature methods 11, 22–24 (2014).

5. Lotfollahi, M., Yuhan, H., Theis, F.J. & Satija, R. The future of rapid and automated single-cell data analysis using reference mapping. Cell 187, 2343–2358 (2024).

6. De Donno, C. et al. Population-level integration of single-cell datasets enables multi-scale analysis across samples. Nature methods 20, 1683–1692 (2023).

7. Hao, Y. et al. Integrated analysis of multimodal single-cell data. Cell 184, 3573–3587.e3529 (2021).

8. Xu, C. et al. Probabilistic harmonization and annotation of single-cell transcriptomics data with deep generative models. Molecular systems biology 17, e9620 (2021).

9. Lotfollahi, M. et al. Mapping single-cell data to reference atlases by transfer learning. Nature biotechnology 40, 121–130 (2022).

10. Novakovsky, G., Dexter, N., Libbrecht, M.W., Wasserman, W.W. & Mostafavi, S. Obtaining genetics insights from deep learning via explainable artificial intelligence. Nature reviews. Genetics 24, 125–137 (2023).

11. Ma, Q., Jiang, Y., Cheng, H. & Xu, D. Harnessing the deep learning power of foundation models in single-cell omics. Nature reviews. Molecular cell biology (2024).

12. Vaswani, A. et al. Attention is All you Need. Neural Information Processing Systems (2017).

13. Devlin, J. BERT: Pre-training of Deep Bidirectional Transformers for Language Understanding. arXiv 1810.04805 (2017).

14. Szałata, A. et al. Transformers in single-cell omics: a review and new perspectives. Nature methods 21, 1430–1443 (2024).

15. Chen, Z., Wei, L. & Gao, G. Foundation models for bioinformatics. *Quantitative Biology* n/a (2024).

16. Cui, H. et al. scGPT: toward building a foundation model for single-cell multi-omics using generative AI. Nature methods (2024).

17. Theodoris, C.V. et al. Transfer learning enables predictions in network biology. Nature 618, 616–624 (2023).

18. Yang, F. et al. scBERT as a large-scale pretrained deep language model for cell type annotation of single-cell RNA-seq data. Nature Machine Intelligence 4, 852–866 (2022).

19. Baars, B.J. A cognitive theory of consciousness. (1988).

20. Baars, B.J. In the theatre of consciousness: Global workspace theory, a rigorous scientific theory of consciousness. Journal of Consciousness Studies 4, 292–309 (1997).

21. Dehaene, S., Kerszberg, M. & Changeux, J.P. A neuronal model of a global workspace in effortful cognitive tasks. Proceedings of the National Academy of Sciences of the United States of America 95, 14529–14534 (1998).

22. Goyal, A., et al. Coordination Among Neural Modules Through a Shared Global Workspace. *ICLR* (2021).

23. 23. Rebecca, B., Nalini, S., Alejandro, B., Gad, G. & David, S. A Deep Dive into Single-Cell RNA Sequencing Foundation Models. *bioRxiv*, 2023.2010.2019.563100 (2023).

24. Duman Keles, F. On The Computational Complexity of Self-Attention. *arXiv* 2209.04881 (2022).

25. Chen, J. et al. Transformer for one stop interpretable cell type annotation. Nat Commun 14, 223 (2023).

26. Domínguez Conde, C., et al. Cross-tissue immune cell analysis reveals tissue-specific features in humans. *Science (New York*, N.Y.) 376, eabl5197 (2022).

27. Fasolino, M. et al. Single-cell multi-omics analysis of human pancreatic islets reveals novel cellular states in type 1 diabetes. Nature metabolism 4, 284–299 (2022).

28. Hrovatin, K. et al. Delineating mouse β-cell identity during lifetime and in diabetes with a single cell atlas. Nature metabolism 5, 1615–1637 (2023).

29. Jorstad, N.L. et al. Comparative transcriptomics reveals human-specific cortical features. *Science (New York*, N.Y.) 382, eade9516 (2023).

30. Kumar, T. et al. A spatially resolved single-cell genomic atlas of the adult human breast. Nature 620, 181–191 (2023).

31. Lu, T.C. et al. Aging Fly Cell Atlas identifies exhaustive aging features at cellular resolution. *Science (New York*, N.Y.) 380, eadg0934 (2023).

32. 32. Mariano, I.G. et al. Integrated multimodal cell atlas of Alzheimer’s disease. *bioRxiv*, 2023.2005.2008.539485 (2023).

33. Salcher, S. et al. High-resolution single-cell atlas reveals diversity and plasticity of tissue-resident neutrophils in non-small cell lung cancer. Cancer cell 40, 1503–1520.e1508 (2022).

34. Steuernagel, L. et al. HypoMap-a unified single-cell gene expression atlas of the murine hypothalamus. Nature metabolism 4, 1402–1419 (2022).

35. 35. Tianyu, L., Kexing, L., Yuge, W., Hongyu, L. & Hongyu, Z. Evaluating the Utilities of Large Language Models in Single-cell Data Analysis. *bioRxiv*, 2023.2009.2008.555192 (2023).

36. Luecken, M.D. et al. Benchmarking atlas-level data integration in single-cell genomics. Nature methods 19, 41–50 (2022).

37. Lopez, R., Regier, J., Cole, M.B., Jordan, M.I. & Yosef, N. Deep generative modeling for single-cell transcriptomics. Nature methods 15, 1053–1058 (2018).

38. Kock, K.H. et al. Single-cell analysis of human diversity in circulating immune cells. *bioRxiv*, 2024.2006.2030.601119 (2024).

39. He, S. et al. High-plex imaging of RNA and proteins at subcellular resolution in fixed tissue by spatial molecular imaging. Nature biotechnology 40, 1794–1806 (2022).

40. Moffitt, J.R. et al. Molecular, spatial, and functional single-cell profiling of the hypothalamic preoptic region. *Science (New York*, N.Y.) 362 (2018).

41. Chen, H. et al. Dissecting mammalian spermatogenesis using spatial transcriptomics. Cell reports 37, 109915 (2021).

42. Aliee, H. & Theis, F.J. AutoGeneS: Automatic gene selection using multi-objective optimization for RNA-seq deconvolution. Cell systems 12, 706–715.e704 (2021).

43. Biancalani, T. et al. Deep learning and alignment of spatially resolved single-cell transcriptomes with Tangram. Nature methods 18, 1352–1362 (2021).

44. Cable, D.M. et al. Robust decomposition of cell type mixtures in spatial transcriptomics. Nature biotechnology 40, 517–526 (2022).

45. Mages, S. et al. TACCO unifies annotation transfer and decomposition of cell identities for single-cell and spatial omics. Nature biotechnology 41, 1465–1473 (2023).

46. Rodriques, S.G. et al. Slide-seq: A scalable technology for measuring genome-wide expression at high spatial resolution. *Science (New York*, N.Y.) 363, 1463–1467 (2019).

47. Janesick, A. et al. High resolution mapping of the tumor microenvironment using integrated single-cell, spatial and in situ analysis. Nat Commun 14, 8353 (2023).

48. Wu, S.Z. et al. A single-cell and spatially resolved atlas of human breast cancers. Nature genetics 53, 1334–1347 (2021).

49. Nguyen, Q.H. et al. Profiling human breast epithelial cells using single cell RNA sequencing identifies cell diversity. Nat Commun 9, 2028 (2018).

50. Zhong, P. et al. Low KRT15 expression is associated with poor prognosis in patients with breast invasive carcinoma. Experimental and therapeutic medicine 21, 305 (2021).

51. Russell, A.J.C. et al. Slide-tags enables single-nucleus barcoding for multimodal spatial genomics. Nature 625, 101–109 (2024).

52. van Bruggen, D. et al. Developmental landscape of human forebrain at a single-cell level identifies early waves of oligodendrogenesis. Developmental cell 57, 1421–1436.e1425 (2022).

53. Chen, A. et al. Spatiotemporal transcriptomic atlas of mouse organogenesis using DNA nanoball-patterned arrays. Cell 185, 1777–1792.e1721 (2022).

54. Zeisel, A. et al. Molecular Architecture of the Mouse Nervous System. Cell 174, 999–1014.e1022 (2018).

55. Yao, Z. et al. A high-resolution transcriptomic and spatial atlas of cell types in the whole mouse brain. Nature 624, 317–332 (2023).

56. Dann, E. et al. Precise identification of cell states altered in disease using healthy single-cell references. Nature genetics 55, 1998–2008 (2023).

57. Alexander, T.B. et al. The genetic basis and cell of origin of mixed phenotype acute leukaemia. Nature 562, 373–379 (2018).

58. Granja, J.M. et al. Single-cell multiomic analysis identifies regulatory programs in mixed-phenotype acute leukemia. Nature biotechnology 37, 1458–1465 (2019).

59. Chen, Y., Liang, Y., Luo, X. & Hu, Q. Oxidative resistance of leukemic stem cells and oxidative damage to hematopoietic stem cells under pro-oxidative therapy. Cell death & disease 11, 291 (2020).

60. Negrin, R.S. Graft-versus-host disease versus graft-versus-leukemia. Hematology. American Society of Hematology. Education Program 2015, 225–230 (2015).

61. van Galen, P. et al. Single-Cell RNA-Seq Reveals AML Hierarchies Relevant to Disease Progression and Immunity. Cell 176, 1265–1281.e1224 (2019).

62. Klein, F. et al. Dntt expression reveals developmental hierarchy and lineage specification of hematopoietic progenitors. Nature immunology 23, 505–517 (2022).

63. Northcott, P.A., et al. Medulloblastoma. Nature reviews. Disease primers 5, 11 (2019).

64. Jessa, S. et al. Stalled developmental programs at the root of pediatric brain tumors. Nature genetics 51, 1702–1713 (2019).

65. Smith, K.S. et al. Unified rhombic lip origins of group 3 and group 4 medulloblastoma. Nature 609, 1012–1020 (2022).

66. Aldinger, K.A. et al. Spatial and cell type transcriptional landscape of human cerebellar development. Nature neuroscience 24, 1163–1175 (2021).

67. Hovestadt, V. et al. Resolving medulloblastoma cellular architecture by single-cell genomics. Nature 572, 74–79 (2019).

68. Dohmen, J. et al. Identifying tumor cells at the single-cell level using machine learning. Genome biology 23, 123 (2022).

69. Zhang, L. et al. Integrated single-cell RNA sequencing analysis reveals distinct cellular and transcriptional modules associated with survival in lung cancer. Signal transduction and targeted therapy 7, 9 (2022).

70. Chen, Y. et al. Club cells employ regeneration mechanisms during lung tumorigenesis. Nat Commun 13, 4557 (2022).

71. Wang, Z. et al. Deciphering cell lineage specification of human lung adenocarcinoma with single-cell RNA sequencing. Nat Commun 12, 6500 (2021).

72. Garcia-Alonso, L. et al. Single-cell roadmap of human gonadal development. Nature 607, 540–547 (2022).

73. Patel, A.P. et al. Single-cell RNA-seq highlights intratumoral heterogeneity in primary glioblastoma. *Science (New York*, N.Y.) 344, 1396–1401 (2014).

74. Deng, Y. et al. Multicellular ecotypes shape progression of lung adenocarcinoma from ground-glass opacity toward advanced stages. *Cell reports*. Medicine 5, 101489 (2024).

75. Hao, X.L. et al. TC2N, a novel oncogene, accelerates tumor progression by suppressing p53 signaling pathway in lung cancer. Cell death and differentiation 26, 1235–1250 (2019).

76. Liang, J. et al. Signatures of malignant cells and novel therapeutic targets revealed by single-cell sequencing in lung adenocarcinoma. Cancer medicine 11, 2244–2258 (2022).

77. Liu, X., Chan, A., Tai, C.H., Andresson, T. & Pastan, I. Multiple proteases are involved in mesothelin shedding by cancer cells. Communications biology 3, 728 (2020).

78. Nomura, M. et al. Niacin restriction with NAMPT-inhibition is synthetic lethal to neuroendocrine carcinoma. Nat Commun 14, 8095 (2023).

79. Liu, X. et al. Immune checkpoint HLA-E:CD94-NKG2A mediates evasion of circulating tumor cells from NK cell surveillance. Cancer cell 41, 272–287.e279 (2023).

80. Liu, K. et al. Tracing the origin of alveolar stem cells in lung repair and regeneration. Cell 187, 2428–2445.e2420 (2024).

81. Qin, Z. et al. EML4-ALK fusions drive lung adeno-to-squamous transition through JAK-STAT activation. The Journal of experimental medicine 221 (2024).

82. Lin, T., Wang, Y., Liu, X. & Qiu, X. A Survey of Transformers. *arXiv e-prints*, arXiv:2106.04554 (2021).

83. Jaegle, A. et al. Perceiver: General Perception with Iterative Attention. *arXiv e-prints*, arXiv:2103.03206 (2021).

84. Liu, D., et al. Discrete-Valued Neural Communication. *arXiv e-prints*, arXiv:2107.02367 (2021).

85. Nagrani, A., et al. Attention Bottlenecks for Multimodal Fusion. *arXiv e-prints*, arXiv:2107.00135 (2021).

86. Zhou, Y. et al. Metascape provides a biologist-oriented resource for the analysis of systems-level datasets. Nat Commun 10, 1523 (2019).

