## Supplementary material for "Hierarchical Interpretation of Out-of-Distribution Cells Using Bottlenecked Transformer": Hierarchical Interpretation of Out-of-Distribution Cells Using Bottlenecked Transformer-bioRxiv

### Supplementary Information

#### Supplementary Figure 1

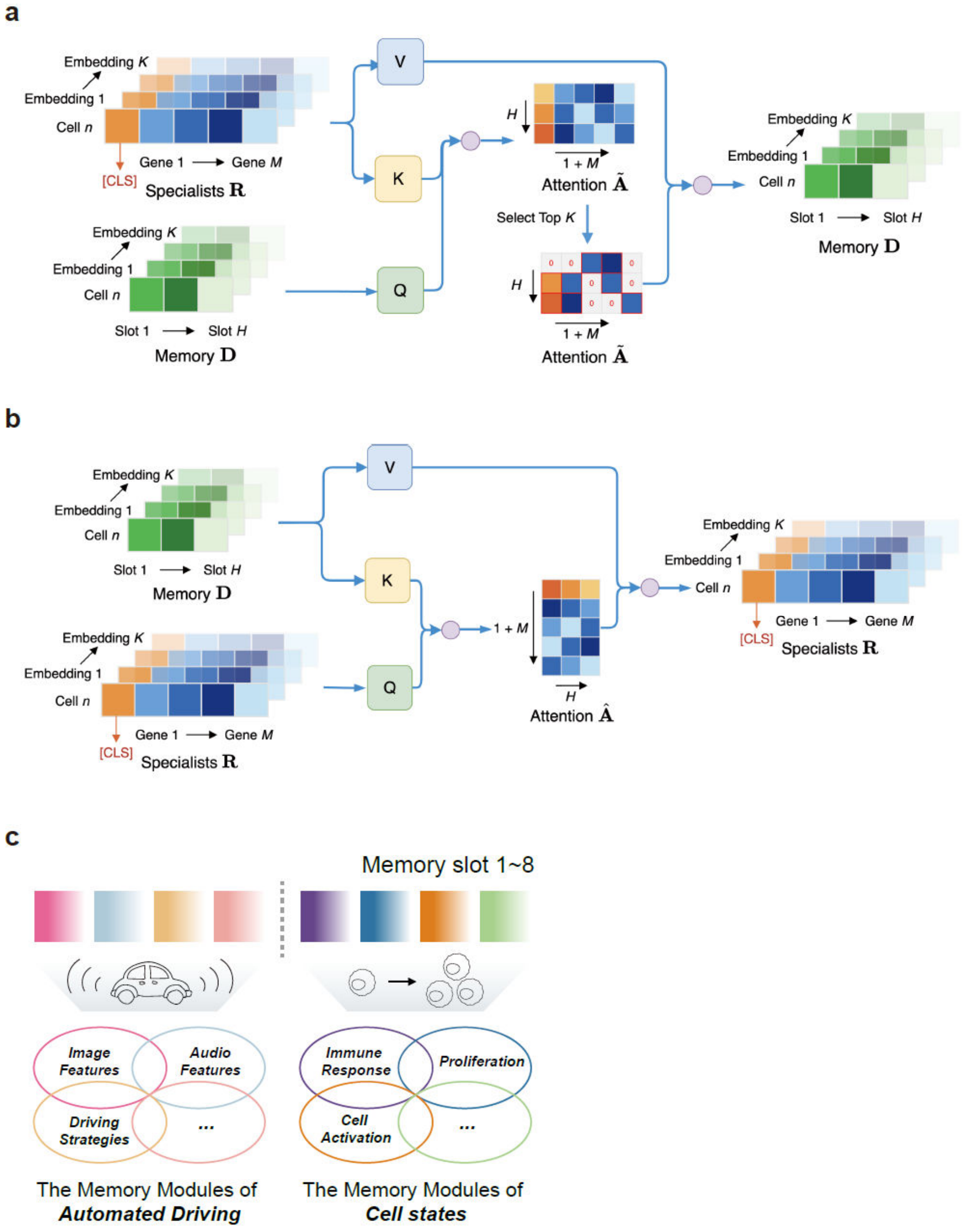


**Supplementary Figure 1. Illustration of CellMemory's writing and broadcasting mechanism.**

**a.** The specialist modules $\mathbf{R}$ will compete to write the information to the shared global workspace, resulting in the memory matrix $\mathbf{D} \in \mathbb{R}^{N \times H \times K}$, with $H$ distinct memory slots serving as a hyper-parameter in our model. Typically, $H$ is chosen to be much smaller than the original input dimension $M$, reflecting the bottleneck of the shared global workspace. The writing procedure involves a cross-attention mechanism between the specialist matrix $\mathbf{R}$ and the memory matrix $\mathbf{D}$.

**b.** The memory matrix $\mathbf{D}$ is broadcasted back to the specialists, leading to the creation of an updated specialist matrix $\mathbf{R}$. This broadcasting action also utilizes a cross-attention mechanism between the specialists and the memory.

**c.** In CellMemory, the memory slots work in concert to accomplish a given task. However, there may be variations among different slots, with each focusing on distinct information, a process that is automatically managed by the model. For instance, in automated driving tasks, different slots might concentrate on visual and auditory features, respectively. Similarly, when inferring cellular representations, such as cell proliferation, various slots could prioritize different functional information, such as immune responses or cell activation.

#### Supplementary Figure 2

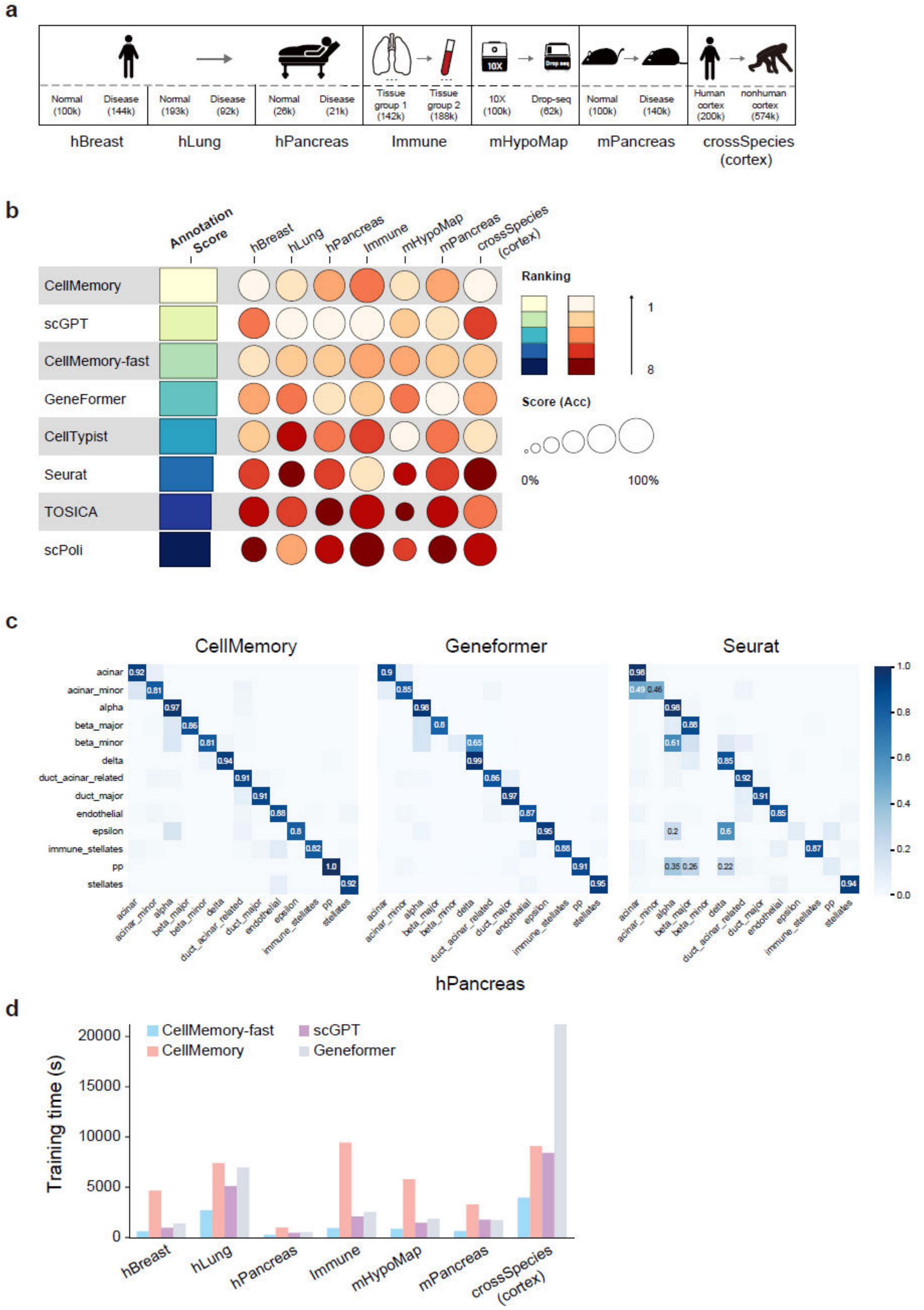


**Supplementary Figure 2. Single-cell annotation benchmark.**

**a.** The number of cells and the states of each dataset's reference and query sets are provided, with cell counts in parentheses. Arrows indicate the direction of prediction.

**b.** Accuracy is employed to assess the performance of single-cell data annotation. The individual scores are then scaled from 0 to 1 using the minimum-maximum scaling approach. The overall score is calculated as the average of the five-fold validation results.

**c.** The hPancreas dataset includes a rare cell type (beta_minor). Although beta_minor constitutes only 0.3% of the query dataset, CellMemory accurately annotates 81% of beta_minor cells, whereas Geneformer and Seurat achieve less than 11% accuracy.

**d.** For each dataset, the average total training time is recorded for CellMemory, scGPT, and Geneformer.

#### Supplementary Figure 3

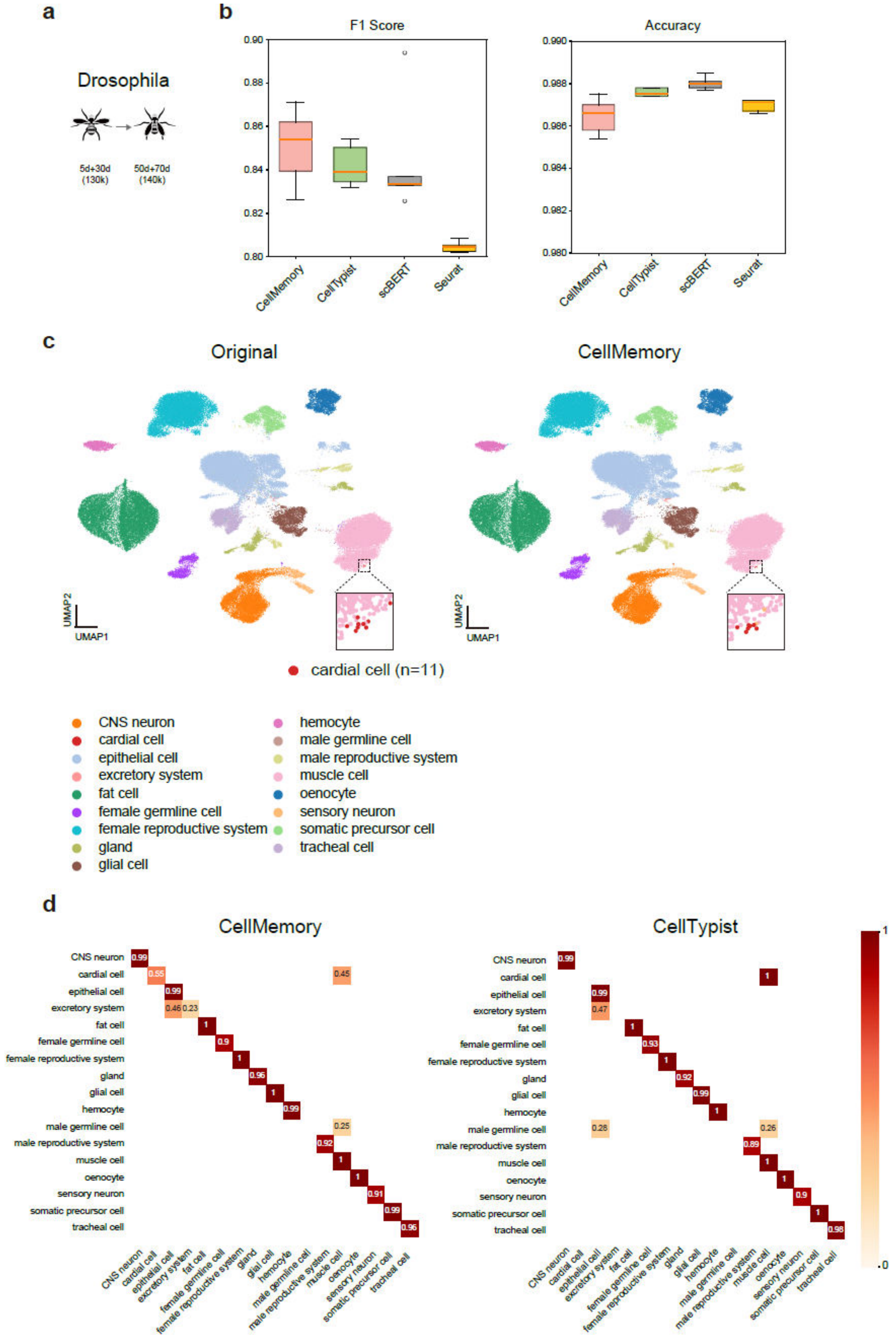


**Supplementary Figure 3. The identification of rare cell types in Drosophila with CellMemory.**

**a.** The models are trained using single-cell data from 5-day and 30-day old Drosophila, then integrated with data from 50-day and 70-day old Drosophila.

**b.** Annotation benchmark. Geneformer and scGPT, which rely on pre-training with gene IDs, are not included in the comparison.
**c.** UMAP generated from integrating 50-day and 70-day old Drosophila cells. The UMAP plots display the original (left) and predicted (right) cell types (n=92,798 cells). In the query dataset, there are only 11 cardial cells (0.012%), a very low proportion. Black dotted boxes indicate the location of cardial cells.
**d.** A comparison with the CellTypist annotation results reveals that CellTypist identified all cardial cells as muscle cells, which are highly similar to cardial cells.

### Supplementary Figure 4

**a**

| Method | NMI | ARI | Cell type ASW | CC conservation | isolated label F1 | isolated label silhouette | Batch ASW | graph connectivity | kBET | Bio conservation | Batch correction | Total |
|---|---|---|---|---|---|---|---|---|---|---|---|---|
| | Bio conservation | | | | | | Batch correction | | | Aggregate score | | |
| CellMemory-fast | 0.71 | 0.74 | 0.78 | 0.52 | 0.52 | 0.62 | 0.95 | 0.99 | 0.80 | 0.65 | 0.92 | 0.76 |
| CellMemory | 0.69 | 0.56 | 0.80 | 0.36 | 0.45 | 0.26 | 0.94 | 0.96 | 0.82 | 0.52 | 0.91 | 0.67 |
| scANVI | 1.00 | 1.00 | 0.67 | 0.24 | 0.29 | 0.11 | 0.81 | 1.00 | 0.71 | 0.55 | 0.84 | 0.67 |
| scVI | 0.47 | 0.37 | 0.19 | 0.20 | 1.00 | 0.12 | 0.92 | 0.98 | 0.58 | 0.39 | 0.83 | 0.57 |
| fastMNN | 0.49 | 0.46 | 0.32 | 0.90 | 0.05 | 0.18 | 0.89 | 0.55 | 0.94 | 0.40 | 0.79 | 0.56 |
| Harmony | 0.52 | 0.41 | 0.36 | 0.00 | 0.00 | 0.88 | 0.75 | 0.61 | 0.91 | 0.36 | 0.76 | 0.52 |
| scPoli | 0.59 | 0.38 | 1.00 | 0.29 | 0.48 | 0.96 | 0.00 | 0.01 | 1.00 | 0.62 | 0.34 | 0.50 |
| Seurat v4 RPCA | 0.61 | 0.64 | 0.35 | 0.91 | 0.34 | 0.00 | 0.49 | 0.53 | 0.35 | 0.48 | 0.45 | 0.47 |
| scGPT | 0.33 | 0.23 | 0.47 | 0.73 | 0.10 | 0.33 | 0.69 | 0.71 | 0.42 | 0.37 | 0.60 | 0.46 |
| ComBat | 0.39 | 0.36 | 0.00 | 0.76 | 0.05 | 1.00 | 0.98 | 0.41 | 0.00 | 0.43 | 0.46 | 0.44 |
| Scanorama | 0.00 | 0.00 | 0.11 | 1.00 | 0.35 | 0.37 | 1.00 | 0.00 | 0.38 | 0.31 | 0.46 | 0.37 |

hBreast

**b**

| Method | NMI | ARI | Cell type ASW | CC conservation | isolated label F1 | isolated label silhouette | Batch ASW | graph connectivity | kBET | Bio conservation | Batch correction | Total |
|---|---|---|---|---|---|---|---|---|---|---|---|---|
| | Bio conservation | | | | | | Batch correction | | | Aggregate score | | |
| CellMemory-fast | 1.00 | 0.98 | 0.50 | 0.00 | 0.96 | 0.16 | 1.00 | 0.87 | 0.30 | 0.60 | 0.72 | 0.65 |
| fastMNN | 0.85 | 0.89 | 0.40 | 0.74 | 0.30 | 0.00 | 0.91 | 0.69 | 0.72 | 0.53 | 0.77 | 0.63 |
| CellMemory | 0.75 | 0.60 | 0.65 | 0.23 | 0.96 | 0.06 | 0.94 | 0.91 | 0.40 | 0.54 | 0.75 | 0.62 |
| Harmony | 0.68 | 0.76 | 0.40 | 0.40 | 0.00 | 1.00 | 0.75 | 0.20 | 0.80 | 0.54 | 0.59 | 0.56 |
| scPoli | 0.99 | 1.00 | 1.00 | 0.14 | 0.36 | 0.69 | 0.00 | 0.00 | 1.00 | 0.70 | 0.33 | 0.55 |
| scANVI | 0.36 | 0.15 | 0.40 | 0.46 | 0.40 | 0.61 | 0.85 | 1.00 | 0.24 | 0.40 | 0.70 | 0.52 |
| scVI | 0.19 | 0.07 | 0.07 | 0.62 | 0.55 | 0.79 | 0.91 | 0.99 | 0.22 | 0.38 | 0.71 | 0.51 |
| Scanorama | 0.22 | 0.07 | 0.21 | 0.90 | 1.00 | 0.15 | 0.97 | 0.76 | 0.05 | 0.42 | 0.59 | 0.49 |
| Seurat v4 RPCA | 0.05 | 0.00 | 0.17 | 1.00 | 0.68 | 0.27 | 0.76 | 0.51 | 0.56 | 0.36 | 0.61 | 0.46 |
| ComBat | 0.00 | 0.10 | 0.00 | 0.70 | 0.37 | 0.66 | 0.77 | 0.52 | 0.00 | 0.31 | 0.43 | 0.35 |
| scGPT | 0.16 | 0.04 | 0.23 | 0.31 | 0.65 | 0.19 | 0.63 | 0.03 | 0.18 | 0.26 | 0.28 | 0.27 |

hLung

**c**

| Method | Bio conservation | | | | | | Batch correction | | | Aggregate score | | |
|---|---|---|---|---|---|---|---|---|---|---|---|---|
| | NMI | ARI | Cell type ASW | CC conservation | isolated label F1 | isolated label silhouette | Batch ASW | graph connectivity | kBET | Bio conservation | Batch correction | Total |
| CellMemory | 0.75 | 0.60 | 0.77 | 0.38 | 1.00 | 0.42 | 0.71 | 0.96 | 0.91 | 0.65 | 0.86 | 0.73 |
| scANVI | 1.00 | 1.00 | 0.19 | 0.27 | 0.19 | 1.00 | 0.96 | 0.83 | 0.79 | 0.61 | 0.86 | 0.71 |
| CellMemory-fast | 0.44 | 0.31 | 0.67 | 0.37 | 0.69 | 0.41 | 0.91 | 1.00 | 1.00 | 0.48 | 0.97 | 0.68 |
| fastMNN | 0.77 | 0.69 | 0.32 | 0.82 | 0.15 | 0.42 | 0.97 | 0.76 | 0.85 | 0.53 | 0.86 | 0.66 |
| scPoli | 0.85 | 0.97 | 1.00 | 0.44 | 0.14 | 0.89 | 0.00 | 0.57 | 0.79 | 0.72 | 0.46 | 0.61 |
| Seurat v4 RPCA | 0.68 | 0.67 | 0.13 | 1.00 | 0.13 | 0.08 | 1.00 | 0.68 | 0.86 | 0.45 | 0.85 | 0.61 |
| scVI | 0.71 | 0.62 | 0.00 | 0.00 | 0.16 | 0.87 | 0.99 | 0.83 | 0.85 | 0.39 | 0.89 | 0.59 |
| Harmony | 0.44 | 0.56 | 0.09 | 0.56 | 0.03 | 0.00 | 0.99 | 0.41 | 0.96 | 0.28 | 0.79 | 0.48 |
| Scanorama | 0.50 | 0.38 | 0.35 | 0.77 | 0.06 | 0.36 | 0.83 | 0.70 | 0.21 | 0.40 | 0.58 | 0.47 |
| ComBat | 0.43 | 0.49 | 0.11 | 0.96 | 0.04 | 0.16 | 0.43 | 0.49 | 0.00 | 0.36 | 0.31 | 0.34 |
| scGPT | 0.00 | 0.00 | 0.17 | 0.45 | 0.00 | 0.27 | 0.03 | 0.00 | 0.14 | 0.15 | 0.06 | 0.11 |

hPancreas

**d**

| Method | Bio conservation | | | | | | Batch correction | | | Aggregate score | | |
|---|---|---|---|---|---|---|---|---|---|---|---|---|
| | NMI | ARI | Cell type ASW | CC conservation | isolated label F1 | isolated label silhouette | Batch ASW | graph connectivity | kBET | Bio conservation | Batch correction | Total |
| CellMemory-fast | 0.84 | 0.92 | 0.81 | 0.72 | 0.29 | 0.38 | 0.91 | 1.00 | 0.71 | 0.66 | 0.87 | 0.75 |
| CellMemory | 1.00 | 1.00 | 0.70 | 0.66 | 0.14 | 0.31 | 0.96 | 0.89 | 0.75 | 0.63 | 0.87 | 0.73 |
| scPoli | 0.73 | 0.71 | 1.00 | 0.53 | 0.24 | 1.00 | 0.00 | 0.75 | 1.00 | 0.70 | 0.58 | 0.65 |
| scANVI | 0.68 | 0.60 | 0.55 | 0.69 | 0.58 | 0.70 | 0.65 | 0.72 | 0.28 | 0.63 | 0.55 | 0.60 |
| scVI | 0.64 | 0.66 | 0.08 | 0.50 | 1.00 | 0.73 | 0.85 | 0.34 | 0.47 | 0.60 | 0.55 | 0.58 |
| Scanorama | 0.70 | 0.67 | 0.07 | 0.87 | 0.15 | 0.00 | 1.00 | 0.68 | 0.48 | 0.41 | 0.72 | 0.53 |
| Harmony | 0.55 | 0.59 | 0.49 | 0.00 | 0.00 | 0.68 | 0.68 | 0.82 | 0.35 | 0.38 | 0.62 | 0.48 |
| Seurat v4 RPCA | 0.58 | 0.65 | 0.22 | 1.00 | 0.06 | 0.02 | 0.76 | 0.21 | 0.29 | 0.42 | 0.42 | 0.42 |
| fastMNN | 0.55 | 0.64 | 0.06 | 0.91 | 0.02 | 0.06 | 0.89 | 0.07 | 0.34 | 0.37 | 0.44 | 0.40 |
| ComBat | 0.55 | 0.66 | 0.15 | 0.81 | 0.05 | 0.08 | 0.95 | 0.00 | 0.00 | 0.38 | 0.32 | 0.36 |
| scGPT | 0.00 | 0.00 | 0.00 | 0.26 | 0.49 | 0.00 | 0.50 | 0.39 | 0.32 | 0.13 | 0.40 | 0.24 |

Immune

**e**

| Method | Bio conservation | | | | | | Batch correction | | | Aggregate score | | |
|---|---|---|---|---|---|---|---|---|---|---|---|---|
| | NMI | ARI | Cell type ASW | CC conservation | isolated label F1 | isolated label silhouette | Batch ASW | graph connectivity | kBET | Bio conservation | Batch correction | Total |
| scANVI | 1.00 | 0.94 | 0.69 | 0.49 | 0.89 | 0.80 | 0.76 | 0.99 | 0.92 | 0.80 | 0.89 | 0.84 |
| fastMNN | 0.50 | 0.52 | 0.66 | 0.91 | 0.92 | 0.64 | 0.85 | 0.81 | 1.00 | 0.69 | 0.89 | 0.77 |
| CellMemory-fast | 0.61 | 0.78 | 0.60 | 0.37 | 0.64 | 0.88 | 0.90 | 0.95 | 0.97 | 0.65 | 0.94 | 0.76 |
| scVI | 0.90 | 1.00 | 0.00 | 0.00 | 0.77 | 0.91 | 0.85 | 1.00 | 0.91 | 0.60 | 0.92 | 0.73 |
| CellMemory | 0.64 | 0.59 | 0.63 | 0.33 | 0.32 | 0.99 | 0.90 | 0.94 | 0.85 | 0.58 | 0.90 | 0.71 |
| Scanorama | 0.49 | 0.61 | 0.25 | 0.83 | 0.89 | 0.75 | 1.00 | 0.69 | 0.52 | 0.64 | 0.73 | 0.68 |
| ComBat | 0.18 | 0.74 | 0.07 | 0.83 | 0.83 | 0.62 | 1.00 | 0.72 | 0.48 | 0.55 | 0.73 | 0.62 |
| Harmony | 0.46 | 0.46 | 0.44 | 0.26 | 0.00 | 0.32 | 0.72 | 0.78 | 0.70 | 0.32 | 0.73 | 0.49 |
| scPoli | 0.33 | 0.91 | 1.00 | 0.38 | 0.22 | 0.00 | 0.00 | 0.27 | 0.67 | 0.47 | 0.31 | 0.41 |
| Seurat v4 RPCA | 0.42 | 0.44 | 0.83 | 1.00 | 0.20 | 0.39 | 0.46 | 0.00 | 0.00 | 0.55 | 0.15 | 0.39 |
| scGPT | 0.00 | 0.00 | 0.31 | 0.10 | 1.00 | 1.00 | 0.24 | 0.28 | 0.42 | 0.40 | 0.31 | 0.37 |

mHypoMap

**f**

| Method | Bio conservation | | | | | | Batch correction | | | Aggregate score | | |
|---|---|---|---|---|---|---|---|---|---|---|---|---|
| | NMI | ARI | Cell type ASW | CC conservation | isolated label F1 | isolated label silhouette | Batch ASW | graph connectivity | kBET | Bio conservation | Batch correction | Total |
| scANVI | 0.83 | 0.86 | 0.37 | 0.76 | 1.00 | 0.74 | 0.70 | 1.00 | 0.74 | 0.76 | 0.81 | 0.78 |
| CellMemory-fast | 0.96 | 0.96 | 0.45 | 0.43 | 0.37 | 0.64 | 0.86 | 0.56 | 0.99 | 0.63 | 0.81 | 0.70 |
| CellMemory | 1.00 | 0.99 | 0.52 | 0.44 | 0.00 | 0.60 | 0.82 | 0.73 | 0.99 | 0.59 | 0.85 | 0.69 |
| fastMNN | 0.86 | 0.87 | 0.26 | 0.83 | 0.12 | 0.55 | 0.84 | 0.83 | 0.92 | 0.58 | 0.86 | 0.69 |
| scVI | 0.85 | 0.92 | 0.00 | 0.26 | 0.12 | 1.00 | 0.90 | 0.88 | 0.65 | 0.52 | 0.81 | 0.64 |
| Harmony | 0.74 | 0.80 | 0.16 | 0.47 | 0.04 | 0.00 | 0.66 | 0.67 | 0.91 | 0.37 | 0.75 | 0.52 |
| Seurat v4 RPCA | 0.76 | 0.86 | 0.31 | 1.00 | 0.21 | 0.31 | 0.59 | 0.51 | 0.05 | 0.57 | 0.38 | 0.50 |
| scPoli | 0.95 | 1.00 | 1.00 | 0.10 | 0.22 | 0.13 | 0.00 | 0.00 | 1.00 | 0.57 | 0.33 | 0.47 |
| ComBat | 0.22 | 0.27 | 0.07 | 0.69 | 0.09 | 0.26 | 0.92 | 0.37 | 0.93 | 0.27 | 0.74 | 0.46 |
| scGPT | 0.16 | 0.12 | 0.09 | 0.00 | 0.15 | 0.85 | 0.65 | 0.51 | 0.92 | 0.23 | 0.69 | 0.41 |
| Scanorama | 0.00 | 0.00 | 0.19 | 0.65 | 0.04 | 0.63 | 1.00 | 0.32 | 0.00 | 0.25 | 0.44 | 0.33 |

mPancreas

**g**

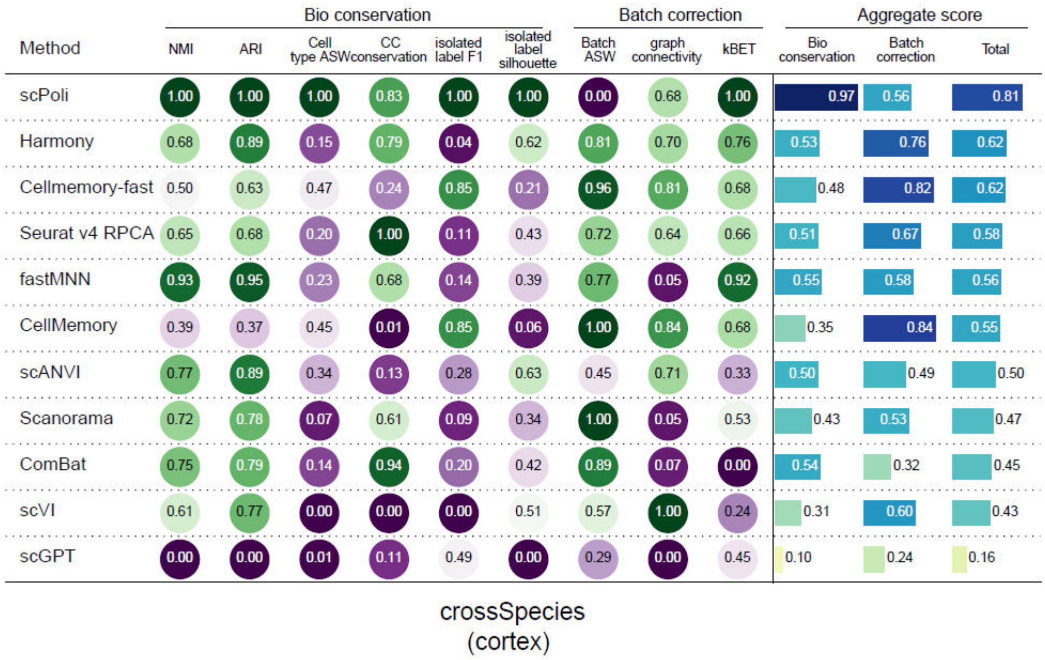


**Supplementary Figure 4. Integration benchmark.**

**a-g.** Evaluation of the integration process for all datasets using the CLS embedding of the query set. The individual scores are scaled from 0 to 1 using the minimum-maximum scaling approach. The overall score is calculated by applying a 40:60 weighted average to the batch correction score and the biological conservation score.

**Supplementary Figure 5**

**a**

| Method | Bio conservation | | | | | | Batch correction | | | Aggregate score | | |
|---|---|---|---|---|---|---|---|---|---|---|---|---|
| | NMI | ARI | Cell type ASW | CC conservation | isolated label F1 | isolated label silhouette | Batch ASW | graph connectivity | kBET | Bio conservation | Batch correction | Total |
| CellMemory-fast | 0.69 | 0.69 | 0.90 | 0.51 | 1.00 | 1.00 | 0.45 | 1.00 | 0.00 | 0.80 | 0.48 | 0.67 |
| CellMemory-fast-bin | 1.00 | 1.00 | 0.00 | 1.00 | 0.00 | 0.19 | 1.00 | 0.00 | 0.90 | 0.53 | 0.63 | 0.57 |
| CellMemory-bin | 0.00 | 0.00 | 0.47 | 0.76 | 0.07 | 0.00 | 0.96 | 0.37 | 1.00 | 0.22 | 0.77 | 0.44 |
| CellMemory | 0.56 | 0.12 | 1.00 | 0.00 | 0.20 | 0.24 | 0.00 | 0.84 | 0.15 | 0.35 | 0.33 | 0.34 |

hBreast

**b**

| Method | Bio conservation | | | | | | Batch correction | | | Aggregate score | | |
|---|---|---|---|---|---|---|---|---|---|---|---|---|
| | NMI | ARI | Cell type ASW | CC conservation | isolated label F1 | isolated label silhouette | Batch ASW | graph connectivity | kBET | Bio conservation | Batch correction | Total |
| CellMemory-fast | 0.83 | 0.90 | 0.23 | 0.75 | 1.00 | 0.68 | 0.70 | 0.76 | 0.00 | 0.73 | 0.48 | 0.63 |
| CellMemory-bin | 0.39 | 0.80 | 0.32 | 0.00 | 0.90 | 1.00 | 1.00 | 0.00 | 1.00 | 0.57 | 0.67 | 0.61 |
| CellMemory-fast-bin | 1.00 | 1.00 | 0.00 | 0.32 | 0.00 | 0.72 | 0.89 | 0.16 | 0.52 | 0.51 | 0.52 | 0.51 |
| CellMemory | 0.00 | 0.00 | 1.00 | 1.00 | 0.99 | 0.00 | 0.00 | 1.00 | 0.44 | 0.50 | 0.48 | 0.49 |

hLung

**c**

| Method | Bio conservation | | | | | | Batch correction | | | Aggregate score | | |
|---|---|---|---|---|---|---|---|---|---|---|---|---|
| | NMI | ARI | Cell type ASW | CC conservation | isolated label F1 | isolated label silhouette | Batch ASW | graph connectivity | kBET | Bio conservation | Batch correction | Total |
| CellMemory-bin | 0.28 | 0.51 | 0.48 | 1.00 | 0.96 | 1.00 | 0.71 | 0.00 | 1.00 | 0.70 | 0.57 | 0.65 |
| CellMemory | 1.00 | 1.00 | 1.00 | 0.01 | 1.00 | 0.02 | 0.00 | 0.53 | 0.00 | 0.67 | 0.18 | 0.47 |
| CellMemory-fast-bin | 0.00 | 0.00 | 0.00 | 0.15 | 0.67 | 0.61 | 1.00 | 1.00 | 0.18 | 0.24 | 0.73 | 0.43 |
| CellMemory-fast | 0.26 | 0.10 | 0.55 | 0.00 | 0.00 | 0.00 | 0.46 | 0.97 | 0.21 | 0.15 | 0.55 | 0.31 |

hPancreas

**d**

| Method | Bio conservation | | | | | | Batch correction | | | Aggregate score | | |
|---|---|---|---|---|---|---|---|---|---|---|---|---|
| | NMI | ARI | Cell type ASW | CC conservation | isolated label F1 | isolated label silhouette | Batch ASW | graph connectivity | kBET | Bio conservation | Batch correction | Total |
| CellMemory-bin | 0.84 | 0.87 | 1.00 | 0.00 | 1.00 | 0.53 | 0.21 | 0.38 | 1.00 | 0.71 | 0.53 | 0.64 |
| CellMemory-fast-bin | 0.79 | 0.86 | 0.00 | 0.45 | 0.51 | 1.00 | 1.00 | 0.00 | 0.64 | 0.60 | 0.55 | 0.58 |
| CellMemory-fast | 0.00 | 0.00 | 0.93 | 1.00 | 0.39 | 0.58 | 0.00 | 1.00 | 0.00 | 0.48 | 0.33 | 0.42 |
| CellMemory | 1.00 | 1.00 | 0.03 | 0.61 | 0.00 | 0.00 | 0.46 | 0.47 | 0.19 | 0.44 | 0.37 | 0.41 |

Immune

**e**

mHypoMap

| Method | Bio conservation: NMI | ARI | Cell type ASW | CC conservation | isolated label F1 | isolated label silhouette | Batch correction: Batch ASW | graph connectivity | kBET | Aggregate score: Bio conservation | Batch correction | Total |
|---|---|---|---|---|---|---|---|---|---|---|---|---|
| CellMemory-fast | 0.00 | 0.78 | 0.86 | 0.28 | 1.00 | 0.00 | 0.00 | 1.00 | 1.00 | 0.49 | 0.67 | 0.56 |
| CellMemory-bin | 0.10 | 1.00 | 0.68 | 0.05 | 0.86 | 0.23 | 1.00 | 0.77 | 0.00 | 0.49 | 0.59 | 0.53 |
| CellMemory | 0.58 | 0.00 | 1.00 | 0.00 | 0.00 | 1.00 | 0.16 | 0.93 | 0.20 | 0.43 | 0.43 | 0.43 |
| CellMemory-fast-bin | 1.00 | 0.60 | 0.00 | 1.00 | 0.60 | 0.23 | 0.05 | 0.00 | 0.56 | 0.57 | 0.20 | 0.42 |

**f**

mPancreas

| Method | Bio conservation: NMI | ARI | Cell type ASW | CC conservation | isolated label F1 | isolated label silhouette | Batch correction: Batch ASW | graph connectivity | kBET | Aggregate score: Bio conservation | Batch correction | Total |
|---|---|---|---|---|---|---|---|---|---|---|---|---|
| CellMemory-bin | 1.00 | 1.00 | 0.99 | 0.00 | 0.15 | 0.00 | 0.65 | 0.11 | 0.88 | 0.52 | 0.55 | 0.53 |
| CellMemory-fast-bin | 0.00 | 0.29 | 0.00 | 1.00 | 0.17 | 1.00 | 1.00 | 0.00 | 1.00 | 0.41 | 0.67 | 0.51 |
| CellMemory-fast | 0.55 | 0.00 | 0.32 | 0.38 | 1.00 | 0.82 | 0.96 | 0.58 | 0.01 | 0.51 | 0.52 | 0.51 |
| CellMemory | 0.76 | 0.18 | 1.00 | 0.44 | 0.00 | 0.24 | 0.00 | 1.00 | 0.00 | 0.44 | 0.33 | 0.39 |

**g**

crossSpecies (cortex)

| Method | Bio conservation: NMI | ARI | Cell type ASW | CC conservation | isolated label F1 | isolated label silhouette | Batch correction: Batch ASW | graph connectivity | kBET | Aggregate score: Bio conservation | Batch correction | Total |
|---|---|---|---|---|---|---|---|---|---|---|---|---|
| Cellmemory-fast-bin | 1.00 | 1.00 | 0.21 | 0.37 | 0.18 | 1.00 | 0.84 | 0.00 | 1.00 | 0.63 | 0.61 | 0.62 |
| CellMemory-bin | 0.13 | 0.35 | 0.00 | 0.50 | 1.00 | 0.32 | 1.00 | 0.20 | 0.98 | 0.38 | 0.73 | 0.52 |
| CellMemory-fast | 0.25 | 0.45 | 1.00 | 1.00 | 0.00 | 0.82 | 0.00 | 0.79 | 0.00 | 0.59 | 0.26 | 0.46 |
| CellMemory | 0.00 | 0.00 | 0.74 | 0.00 | 0.00 | 0.00 | 0.96 | 1.00 | 0.17 | 0.12 | 0.71 | 0.36 |

**h**

Summary

| Method | hBreast | hLung | hPancreas | immune | mHypoMap | mPancreas | crossSpecies (cortex) | Integration Score (Total) |
|---|---|---|---|---|---|---|---|---|
| CellMemory-bin | 0.44 | 0.61 | 0.65 | 0.64 | 0.53 | 0.53 | 0.52 | 0.56 |
| CellMemory-fast-bin | 0.57 | 0.51 | 0.43 | 0.58 | 0.42 | 0.51 | 0.62 | 0.52 |
| CellMemory-fast | 0.67 | 0.63 | 0.31 | 0.42 | 0.56 | 0.51 | 0.46 | 0.51 |
| CellMemory | 0.34 | 0.49 | 0.47 | 0.41 | 0.43 | 0.39 | 0.36 | 0.42 |

**Supplementary Figure 5. Evaluation of different token processing strategies on integration.**

**a-g,** Impact of different token handling strategies on integration for each dataset. CellMemory: All input genes included; CellMemory-fast: Filtering out features with zero expression in each sample; CellMemory-bin: Grouping non-zero expression features of each sample into the integer ranges of 1-bins.

**h.** Impact of different token processing strategies on the comprehensive integration performance across all datasets.

#### Supplementary Figure 6

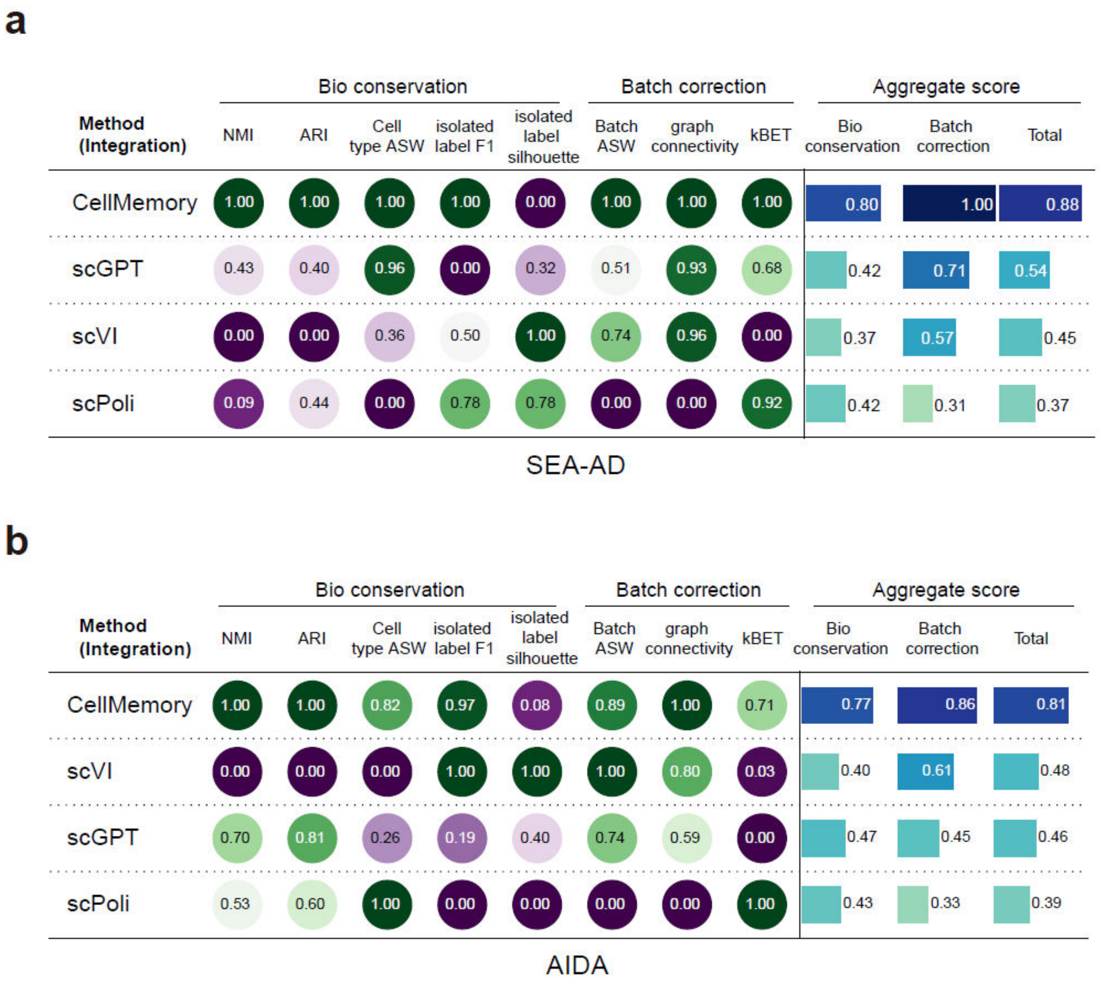


**Supplementary Figure 6. Integration benchmark of population-scale datasets.**

**a-b.** Evaluation of the integration process for the SEA-AD (**a**) and AIDA (**b**) datasets using the CLS embedding of the query set. The individual scores are scaled from 0 to 1 using the minimum-maximum scaling approach. The overall score is calculated by applying a 40:60 weighted average to the batch correction score and the biological conservation score.

#### Supplementary Figure 7

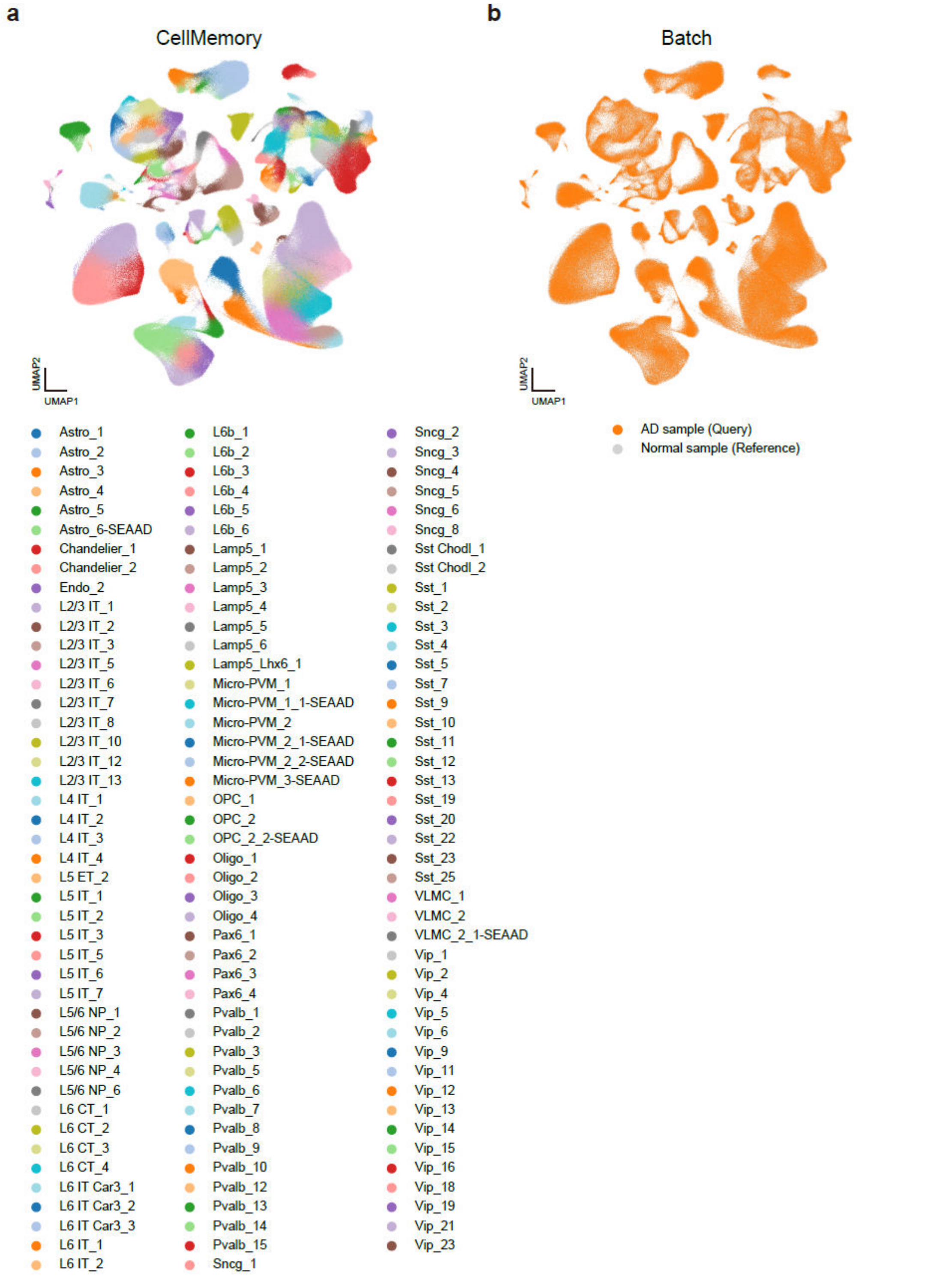


**Supplementary Figure 7. CellMemory ensures granule-level reference mapping for population-scale single-cell Alzheimer's patient cortex dataset.**

**a-b.** UMAP generated by CellMemory for the AD dataset, labeled with original annotations (a) and data information (b).

#### Supplementary Figure 8

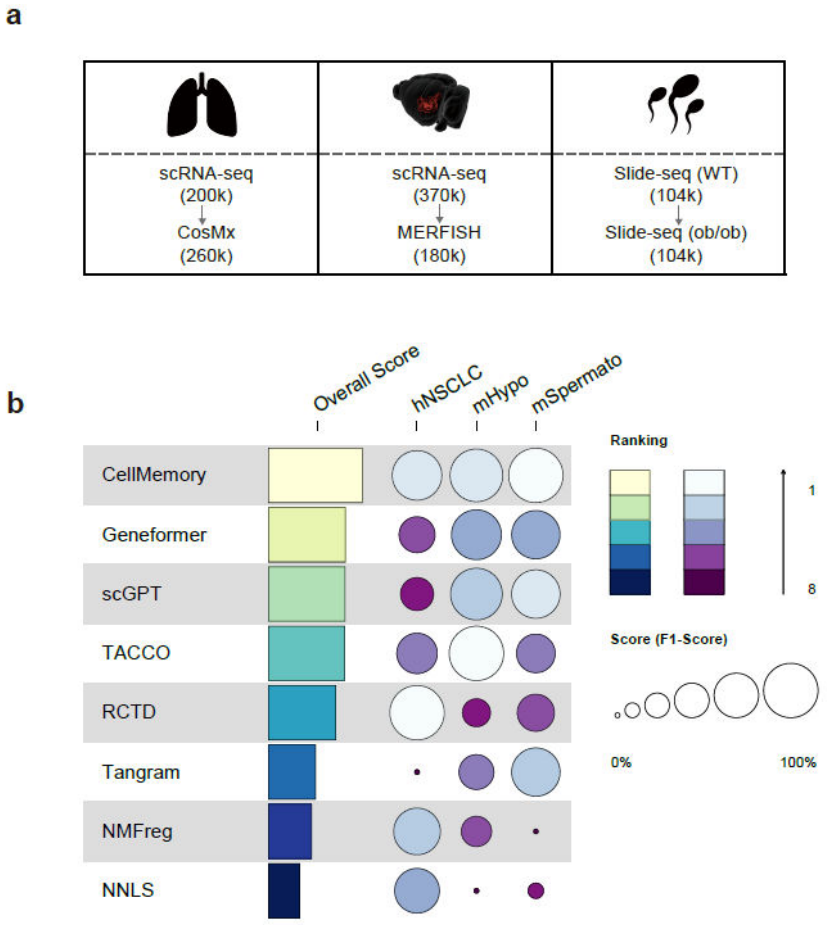


**Supplementary Figure 8. Spatial data annotation benchmark.**

**a.** Sample sizes and sequencing platforms for each prediction scenario's reference and query sets. Arrows indicate the direction of prediction, with cell counts in parentheses.

**b.** Spatial annotation tools were assessed for F1-score using the mHypo, hNSCLC, and mSpermato datasets.

#### Supplementary Figure 9

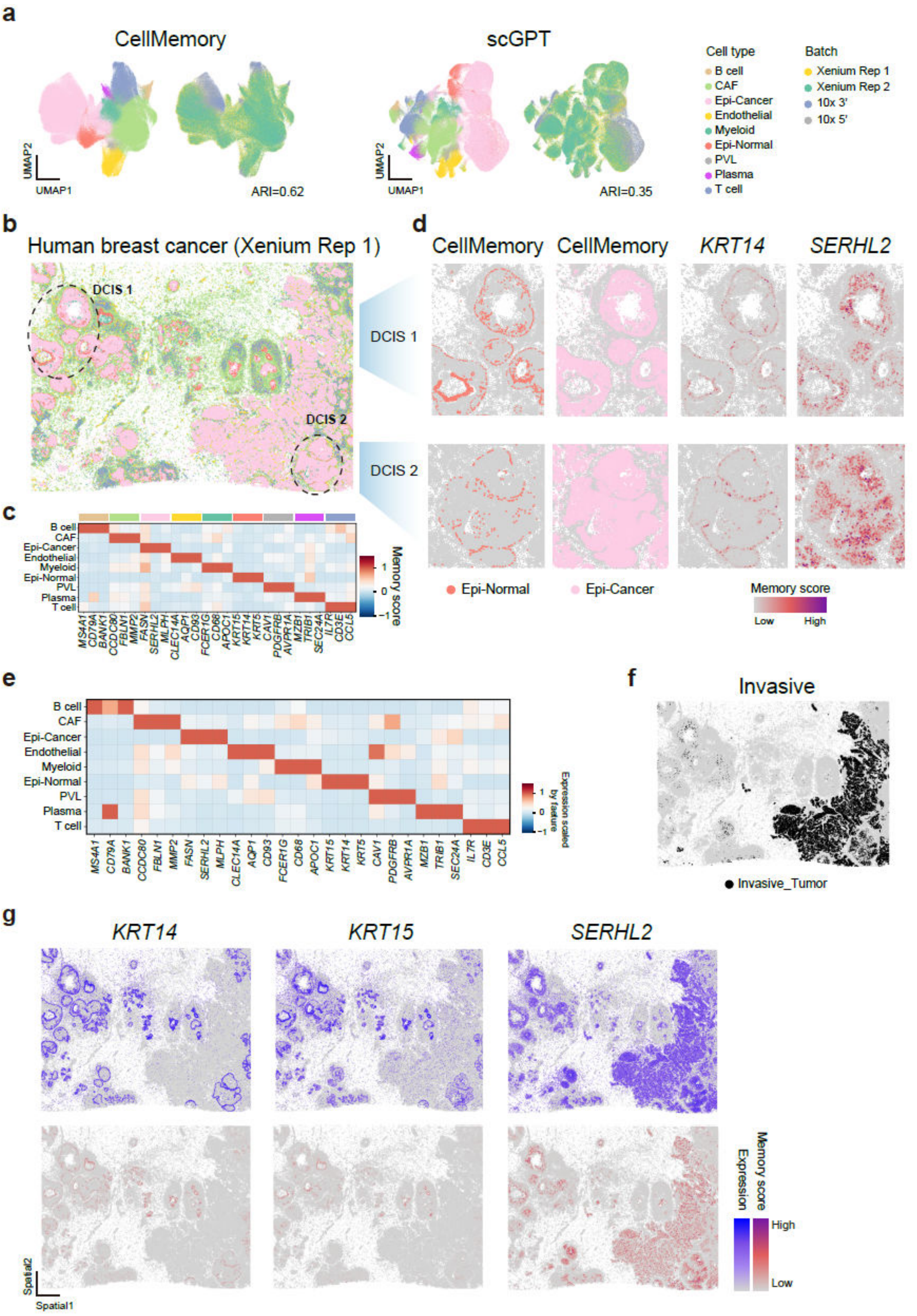


**Supplementary Figure 9. CellMemory accurately characterizes 10x Xenium single-cell spatial data.**

**a.** UMAP visualization of 10x Xenium data, representing human breast cancer samples with two replicates, alongside 10x 3'/5' data, integrated by CellMemory and scGPT. The color is with their respective transferred labels and batches.

**b.** Spatial coordinates of the human breast cancer sample, with cell types transferred by CellMemory, are depicted.

**c.** The heatmap displays the expression of TAGs for each cell type in the Xenium data.

**d.** Spatial coordinates identified by CellMemory for Epi-Normal and Epi-Cancer cells, along with memory scores for *KRT14* and *SERHL2*.

**e.** Heatmap showing the expression of TAGs for each cell type emphasized by CellMemory.

**f.** The spatial coordinates of a human breast cancer sample annotated Invasive tumor cells (**d**) from the original study.

**g.** The expression values and memory scores for *ACTA2*, *KRT15*, and *SERHL2* in the spatial coordinates as displayed.

#### Supplementary Figure 10

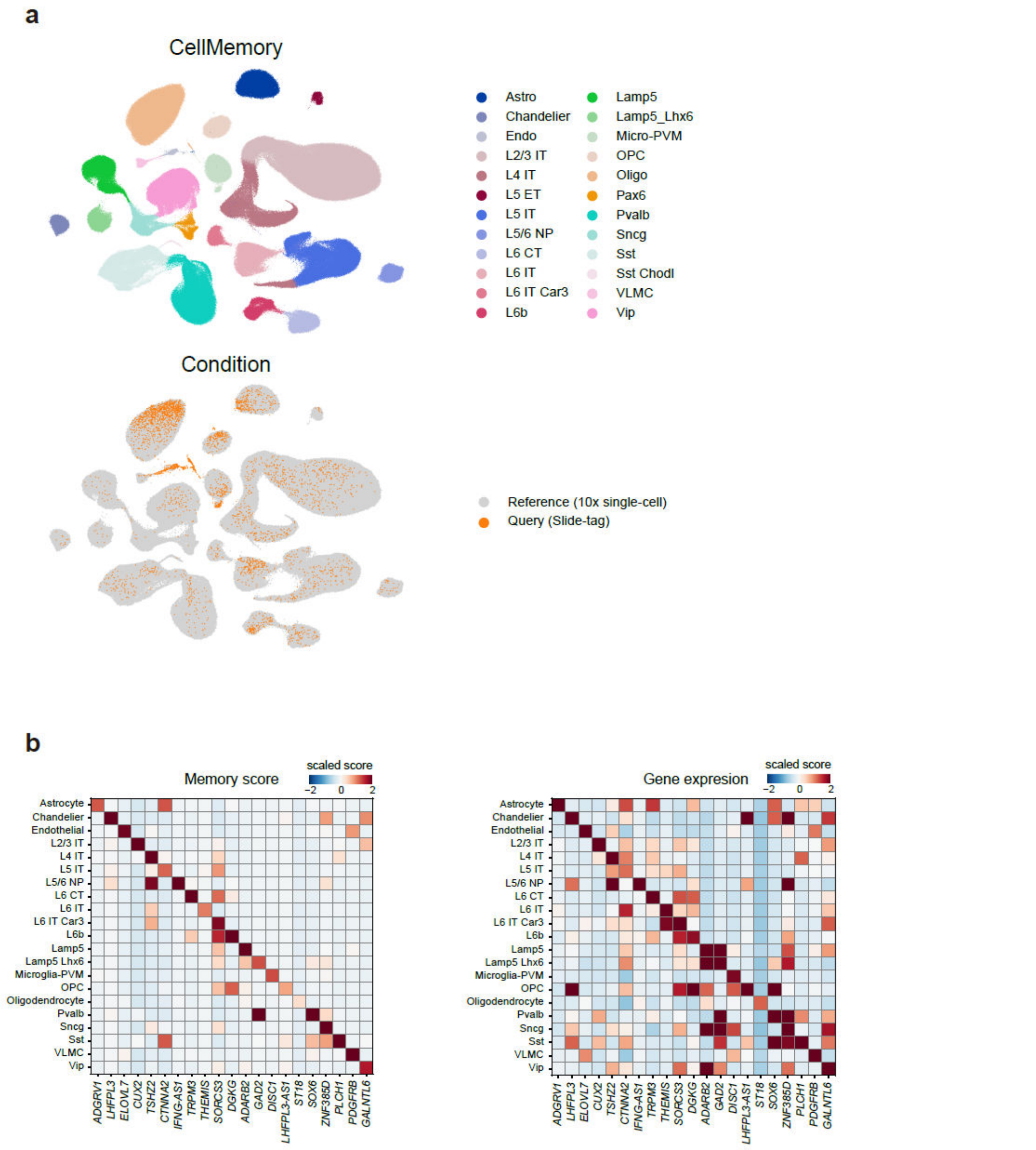


**Supplementary Figure 10. CellMemory integrates single-cell spatial transcriptome data with interpretability (L2 resolution).**

**a.** Construction of a reference with L2 resolution using single-cell data and integration of spatial data from Slide-tags. The top figure is colored according to CellMemory's predictions, while the bottom figure shows reference cells in light gray and query cells colored by condition.

**b.** The memory score/expression of TAGs for each cell type in Slide-tags data (remaining cell types with more than five cells).

#### Supplementary Figure 11

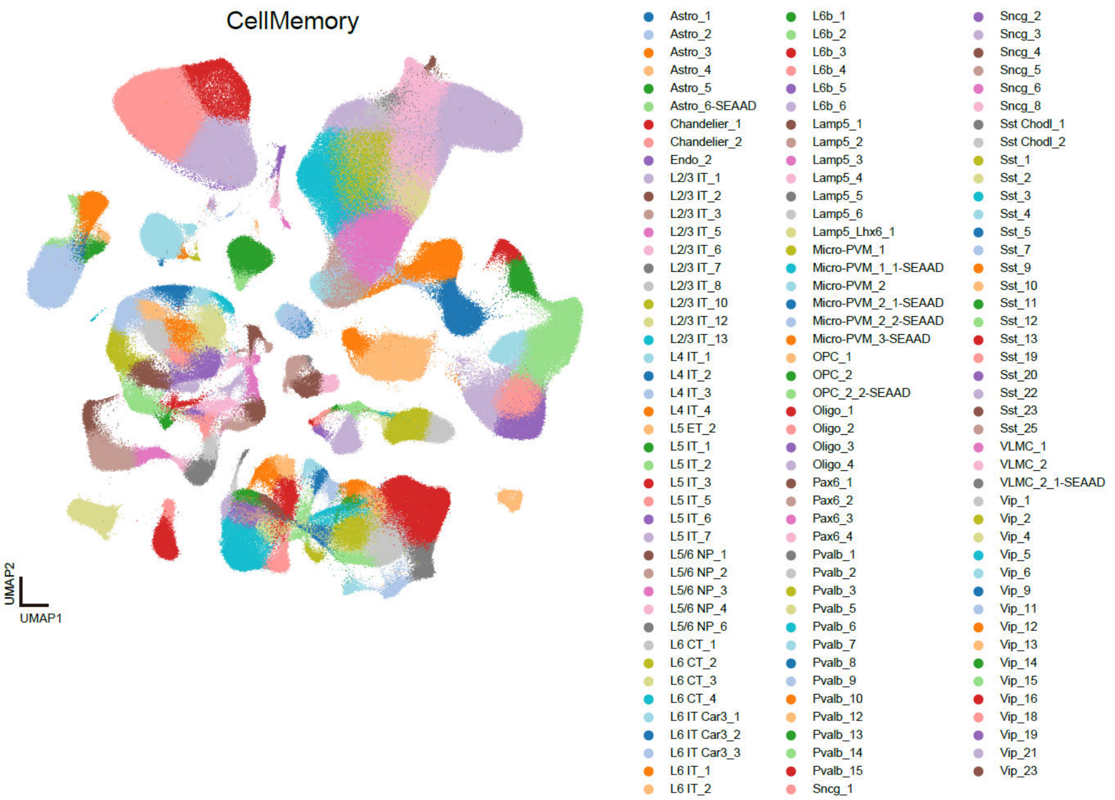


**Supplementary Figure 11. Integration of Slide-tags data using a single-cell dataset containing 131 cell types (L3) generated from the human cortex.**

#### Supplementary Figure 12

a

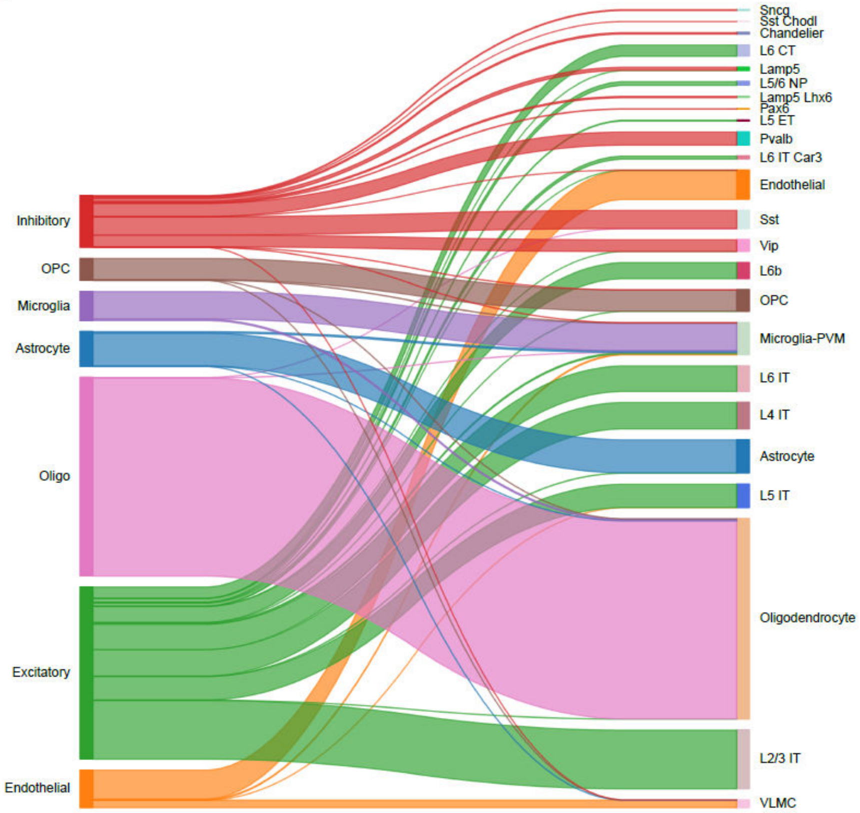

b

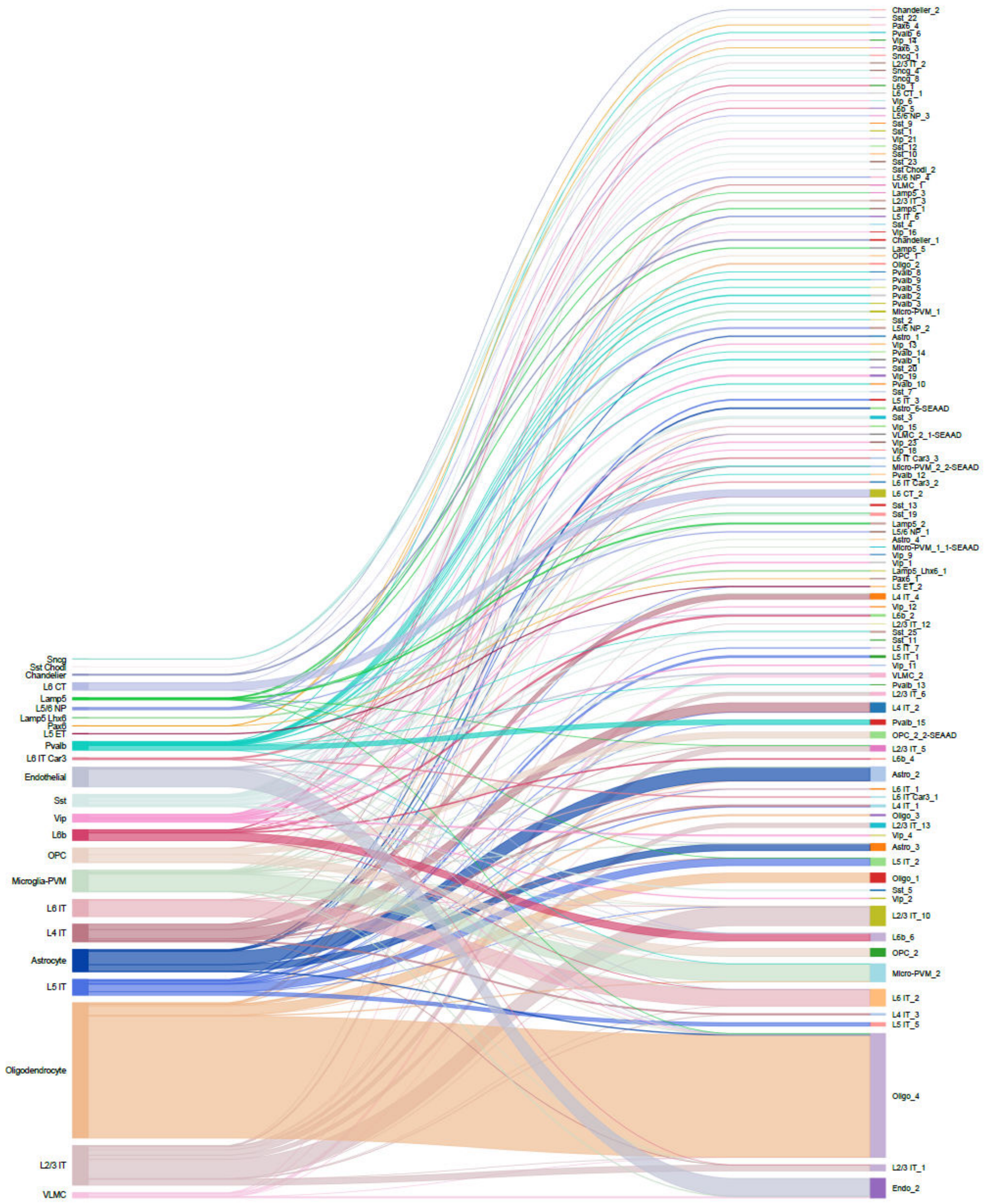


**Supplementary Figure 12. The coherence and consistency of cell type annotation at different resolutions.**

**a.** Sankey diagram linking original labels (L1, 7 categories) with CellMemory's predicted level 2 labels (L2, 24 categories). The colors in the blocks represent the color of cells in the UMAP, with connecting lines matching the original labels on the left. This demonstrates the contribution of broad categories to high-resolution cell types.

**b.** Sankey diagram connecting CellMemory's predicted level 2 (24 categories) and level 3 (112 categories) labels. The colors in the blocks represent the color of cells in the UMAP, with connecting lines consistent with L2 labels on the left, indicating the predominant connections of terms on the right to single broad cell type.

**Supplementary Figure 13**

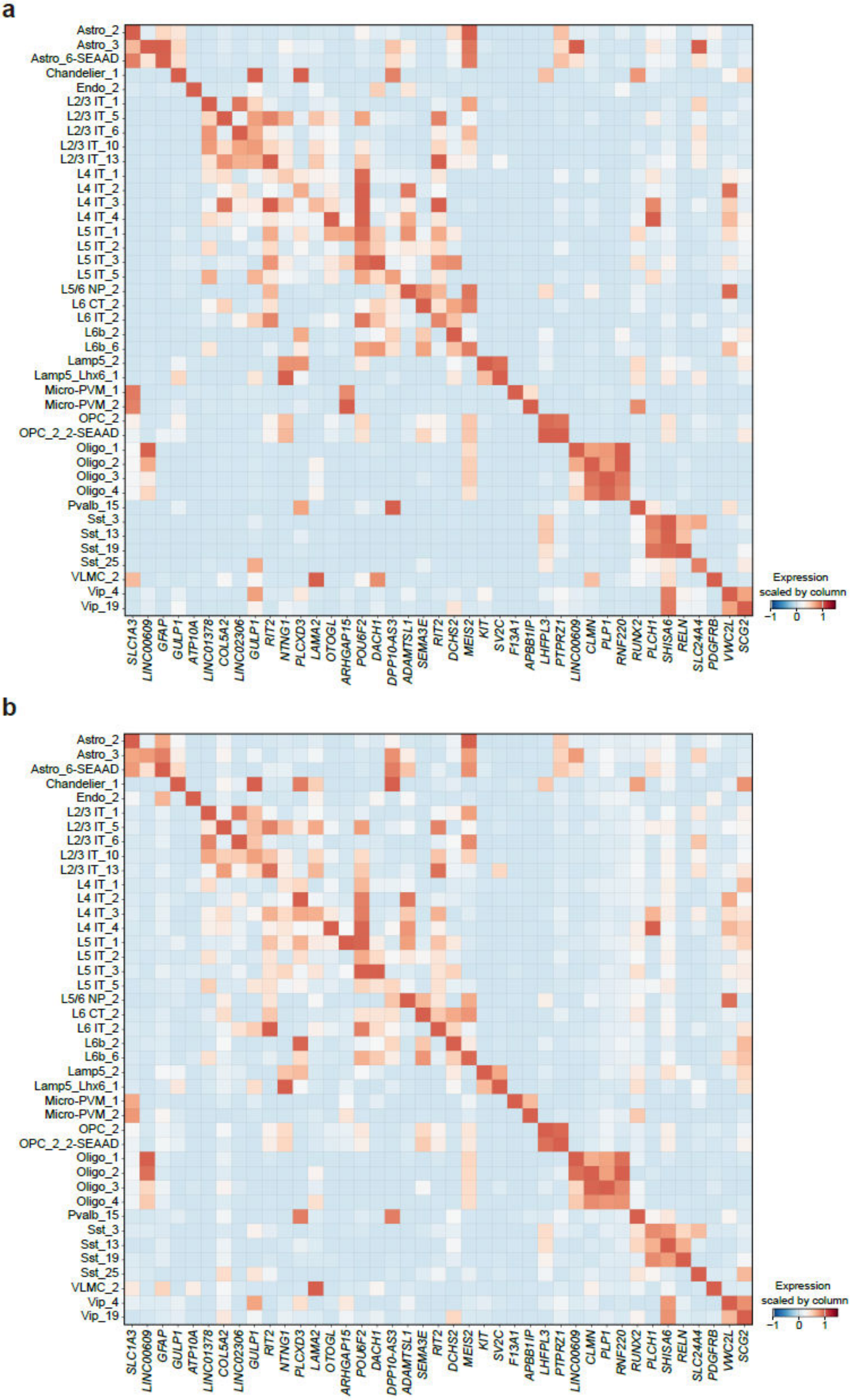


**Supplementary Figure 13. Marker expression of cell states at L3 resolution.**

**a-b.** This figure shows cell states (total 41) with more than 10 cells in Slide-tags data, comparing their expression in 10x single-cell (**a**, reference) and Slide-tags (**b**, query) data. It demonstrates consistent expression trends.

#### Supplementary Figure 14

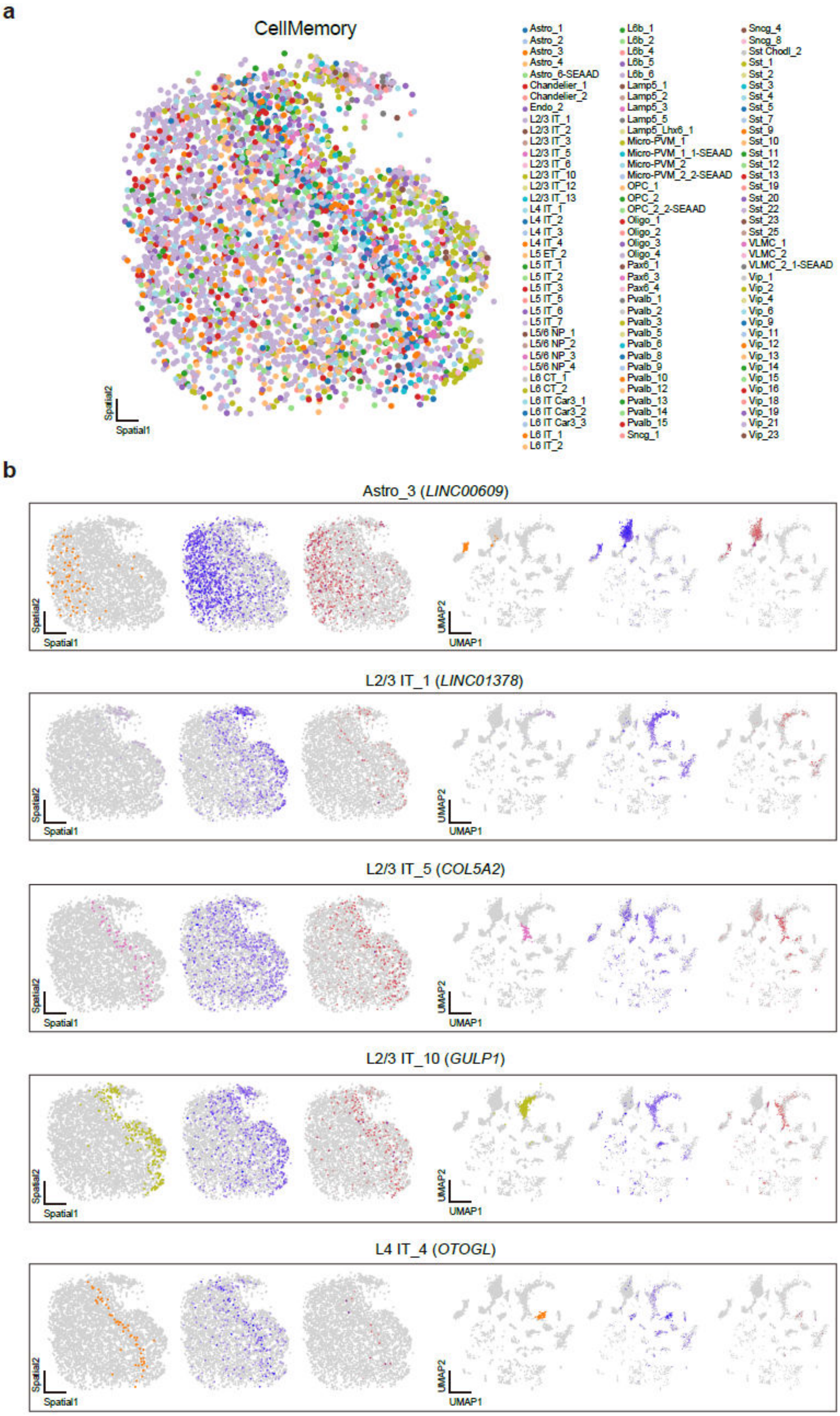

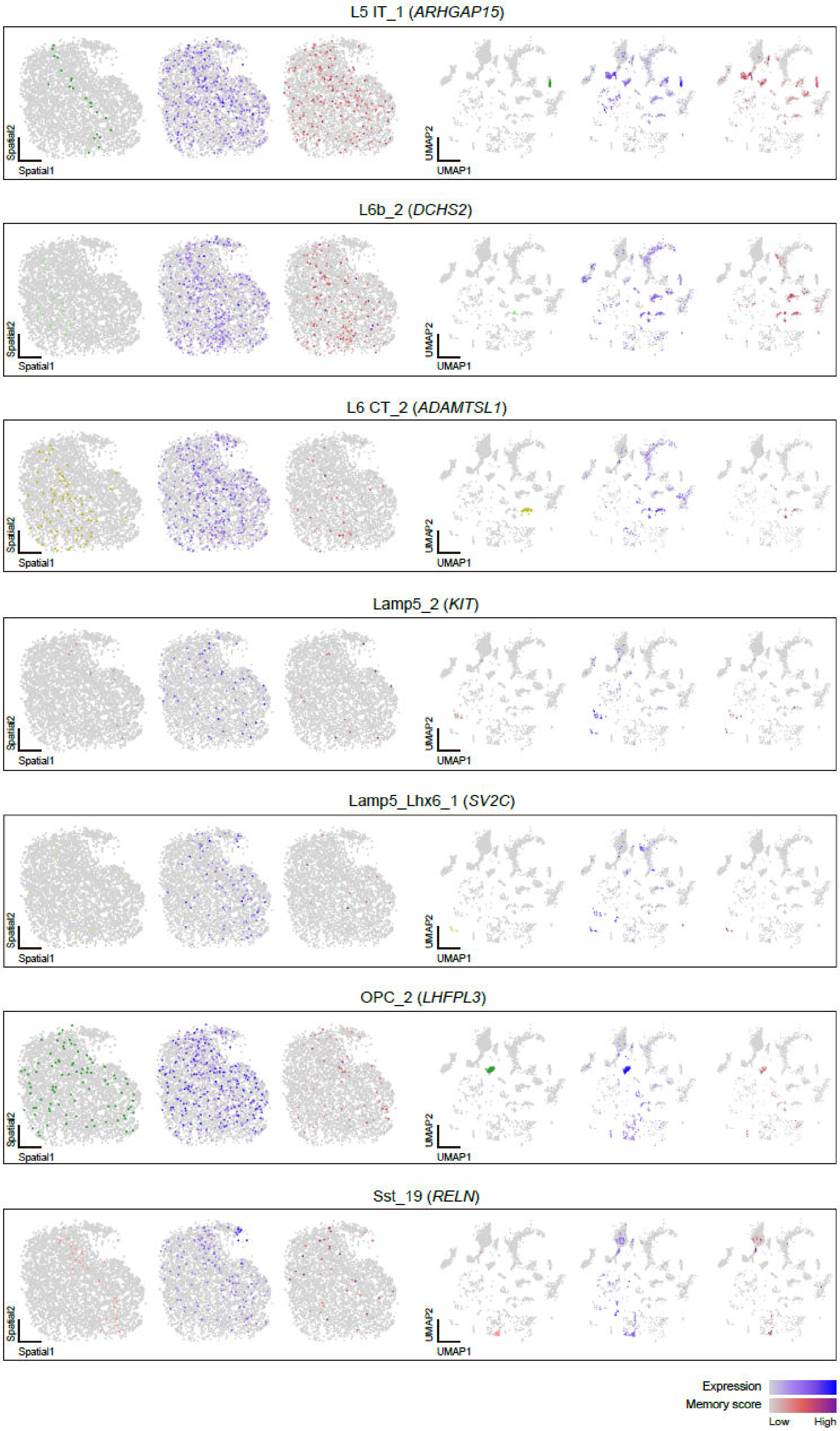

L5 IT_1 (ARHGAP15)
L6b_2 (DCHS2)
L6 CT_2 (ADAMTSL1)
Lamp5_2 (KIT)
Lamp5_Lhx6_1 (SV2C)
OPC_2 (LHFPL3)
Sst_19 (RELN)
Spatial1
Spatial2
UMAP1
UMAP2
Expression
Memory score
Low
High

**Supplementary Figure 14. The interpretability of single-cell spatial transcriptome data integration at the granule-level with CellMemory.**

**a.** Spatial coordinates of the human cortex with L3 annotations from CellMemory.

**b.** Spatial coordinates and UMAP embedding of Astro_3, L2/3 IT_1, L2/3 IT_5, L2/3 IT_10, L4 IT_4, L5 IT_1, L6b_2, L6 CT_2, Lamp5_2, Lamp5_Lhx6_1, OPC_2, Sst_19 in the sample, along with TAGs expression and memory scores.

**Supplementary Figure 15**

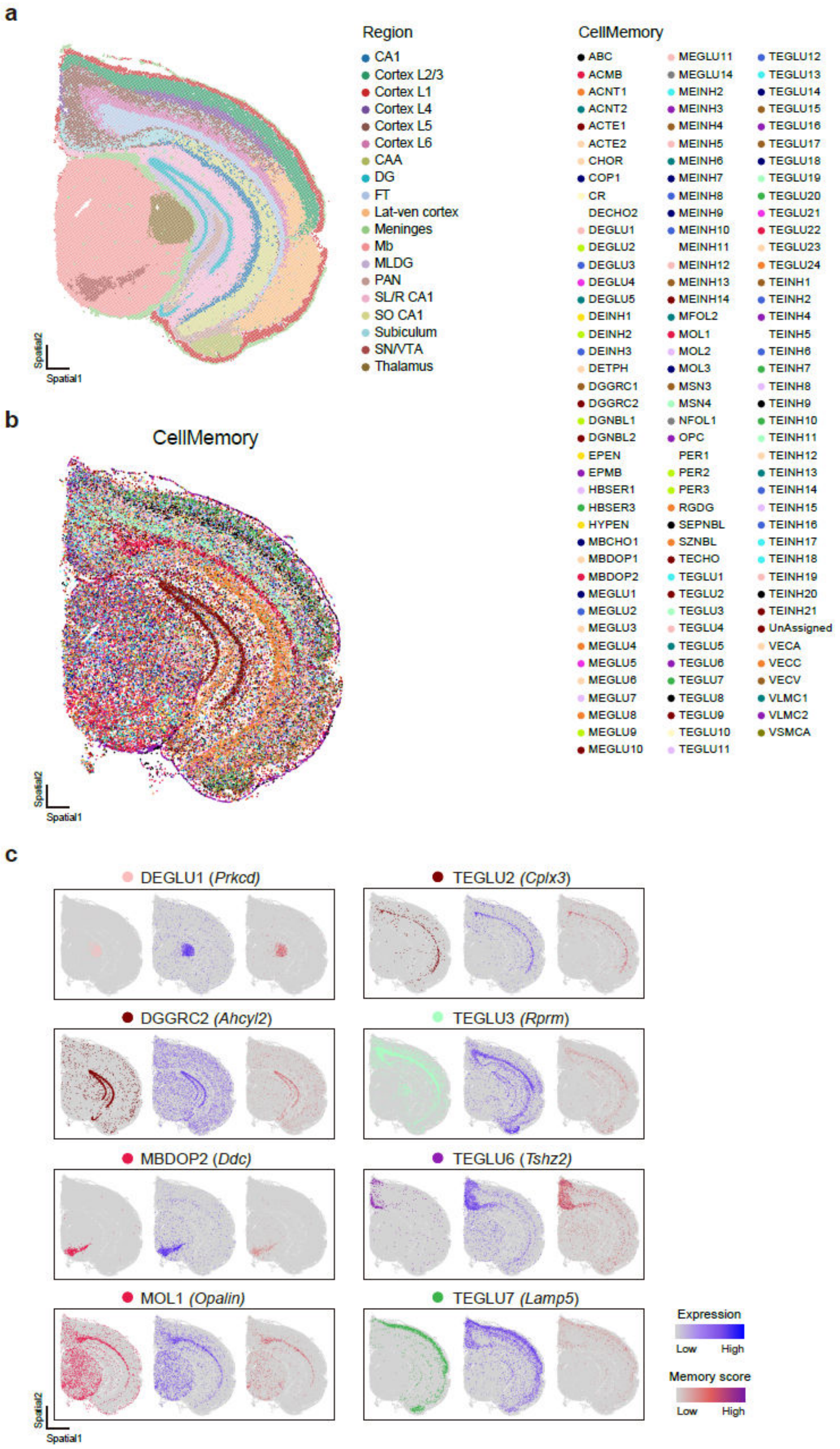

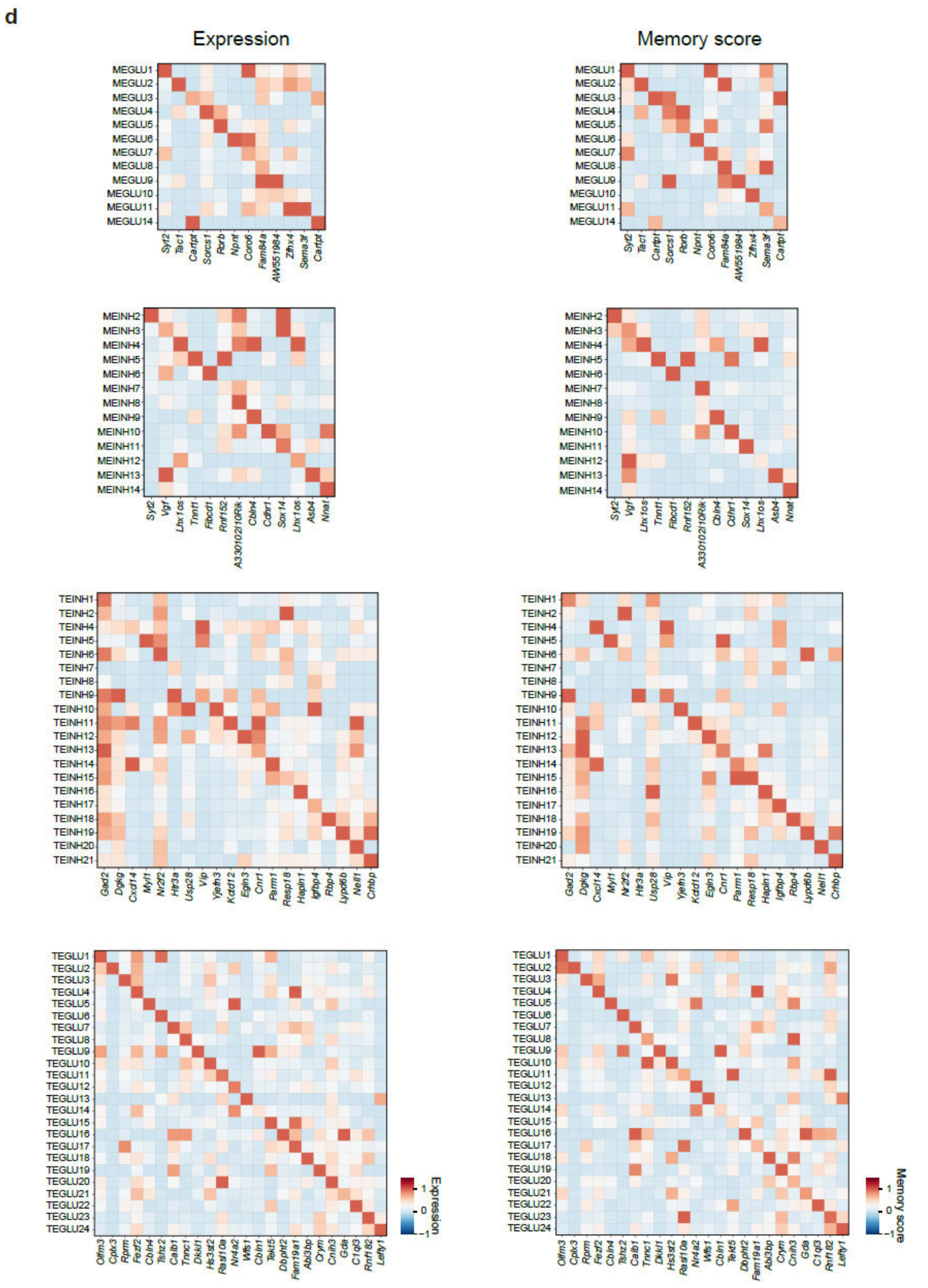


**Supplementary Figure 15. Interpretation of mouse hemibrain Stereo-seq data using CellMemory.** **a.** Adult mouse coronal hemibrain section based on Stereo-seq data. Colors represent brain regions identified through the original research. Left: Lat-ven, lateral-ventral cortex; CAA, cortical amygdalar area; PAN, posterior amygdalar nucleus; FT, fiber tract; SO CA1, stratum oriens area 1; CA1, cornu ammonis area 1; SL/R CA1, stratum lacunosum/raditum cornu ammonis area 1; MLDG, molecular layer of dentate gyrus; DG, dentate gyrus; CA3, cornu ammonis area 3; Mb, midbrain; SN/VTA, substantia nigra/ventral tegmental area.

**b.** Adult mouse coronal hemibrain section based on Stereo-seq data. Cell type annotation of CellMemory for the adult mouse coronal hemibrain section, and UMAP visualization.

**c.** Eight cell types in the mouse hemibrain annotated by CellMemory (left). The normalized signal of cell type-specific genes captured by stereo-seq and their spatial distribution (middle). The gene-specific memory scores are generated by CellMemory (right).

**d.** The scaled mean z-score heatmap showing the specificity of the gene signal and memory score for TAGs associated with the identified excitatory neurons.

### Supplementary Figure 16

**a**

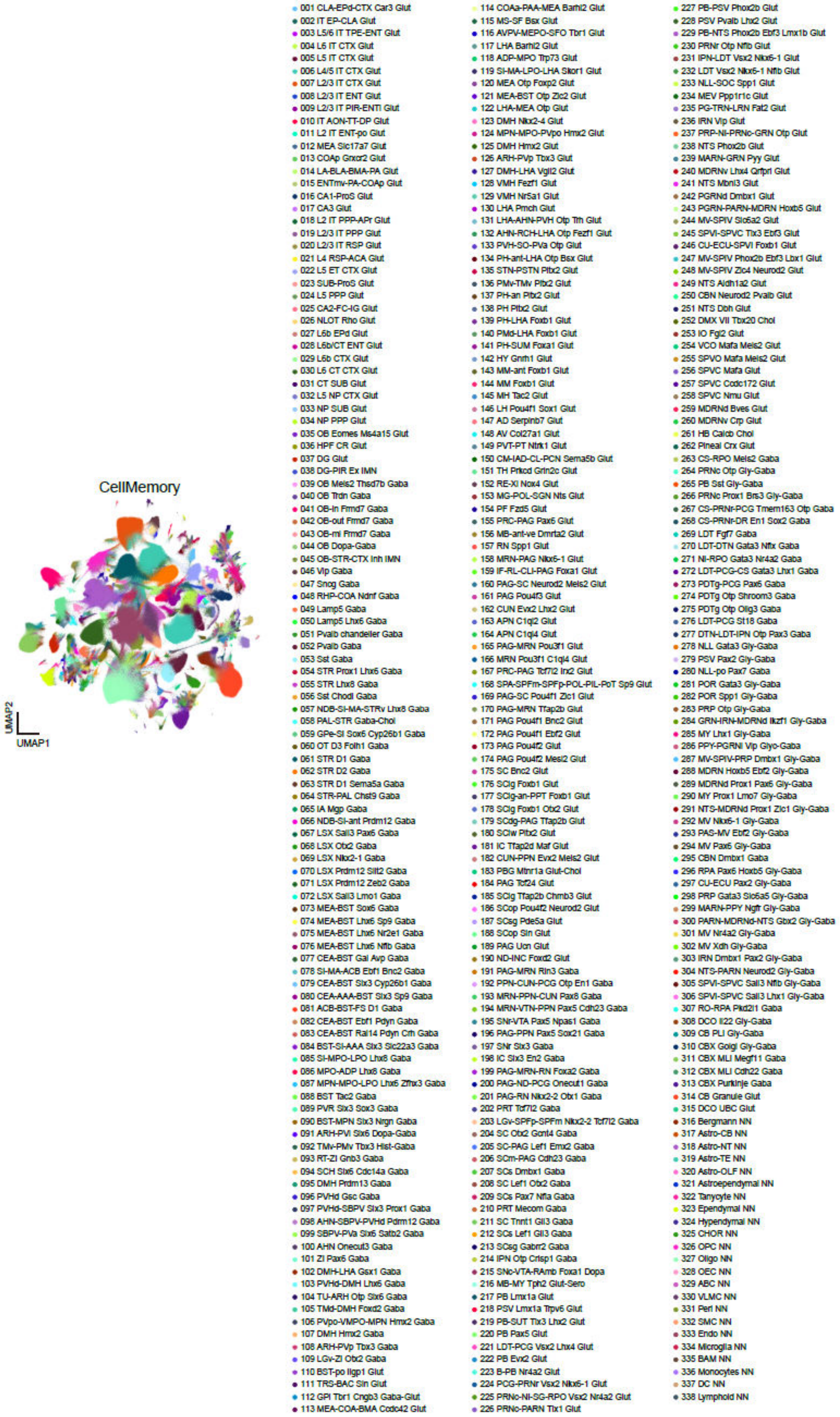


**Supplementary Figure 16.** UMAP representation of 4 million mouse whole brain MERFISH cells, colored by subclasses.

### Supplementary Figure 17

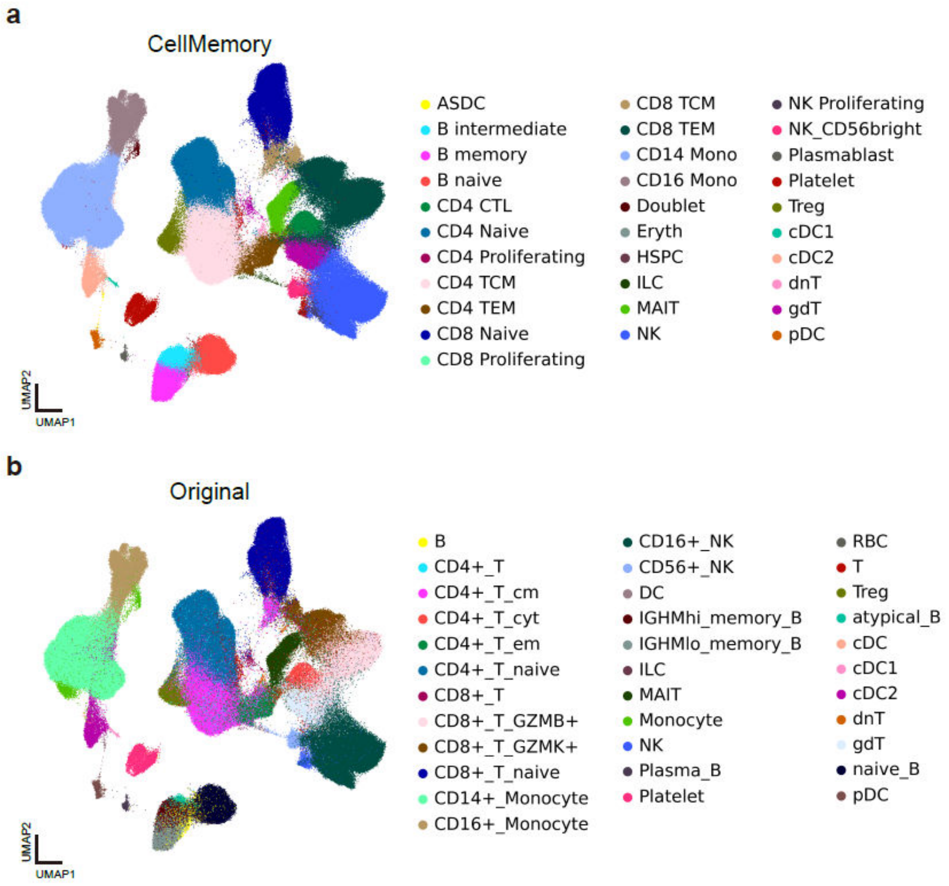


**Supplementary Figure 17. Cross-platform and cross-ethnicity integration of AIDA by CellMemory.**

**a-b.** CLS embeddings generated by CellMemory for AIDA, colored by CellMemory-predicted cell identities (**a**) and original cell types (**b**). To ensure the reliability of the observations, cells from 30 randomly selected donors are retained for each Asian ethnicity.

#### Supplementary Figure 18

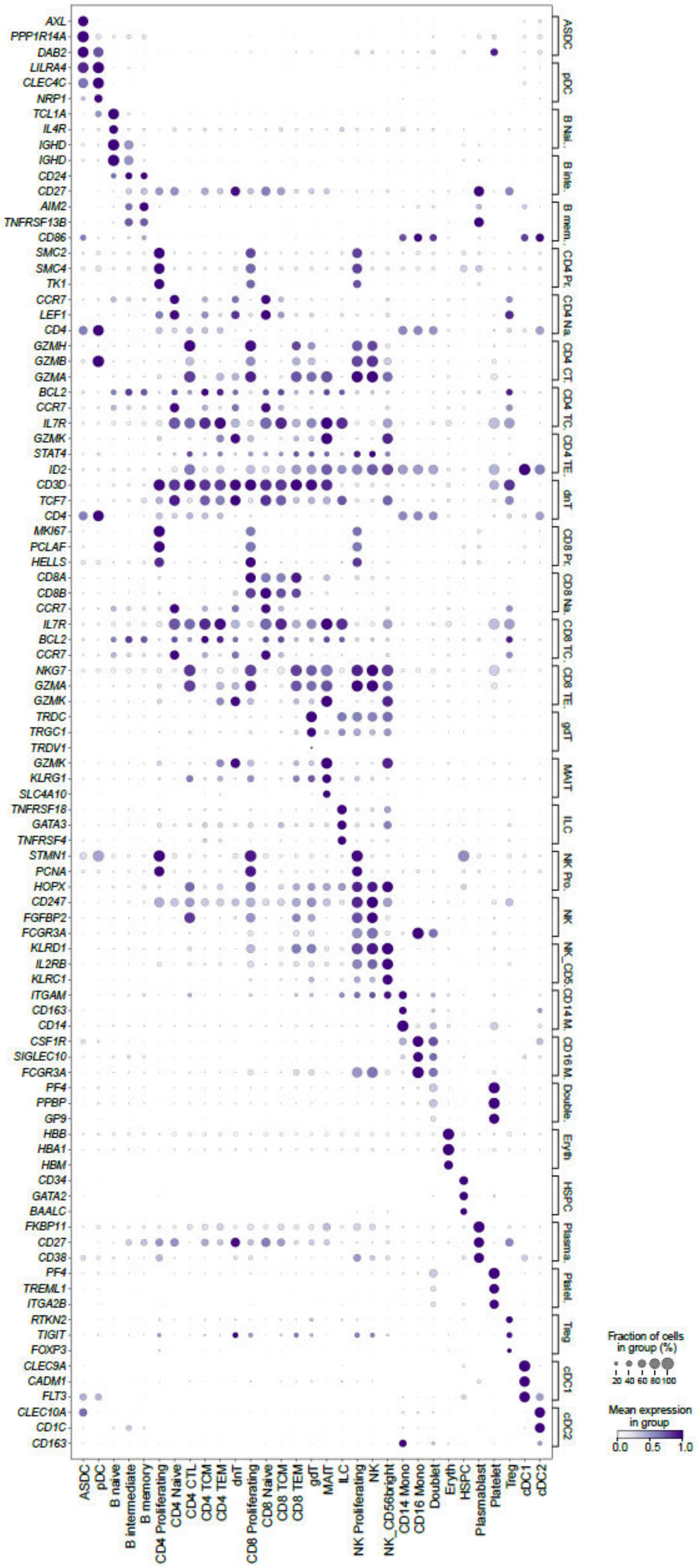


**Supplementary Figure 18. Identification of marker gene expression by CellMemory for each cell type.**

#### Supplementary Figure 19

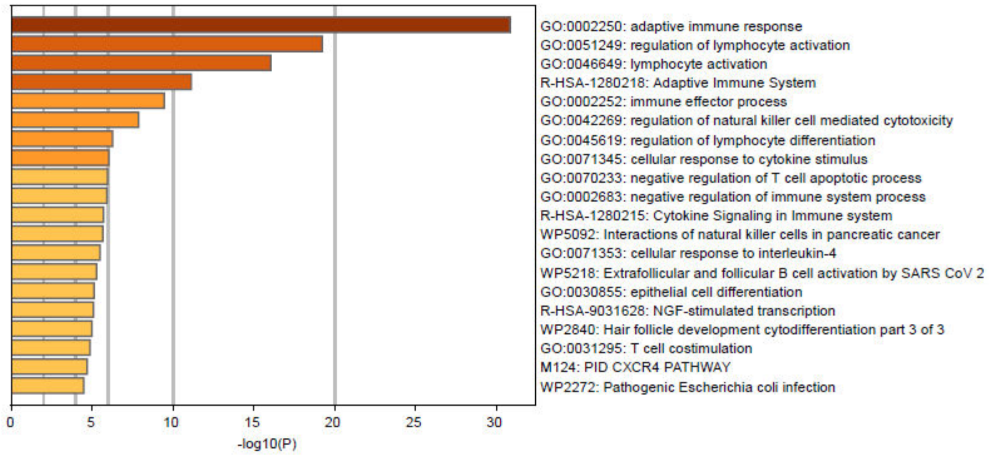


**Supplementary Figure 19. The results of the enrichment analysis for the top 1-100 TAGs of CD8 TCM cells.**

#### Supplementary Figure 20

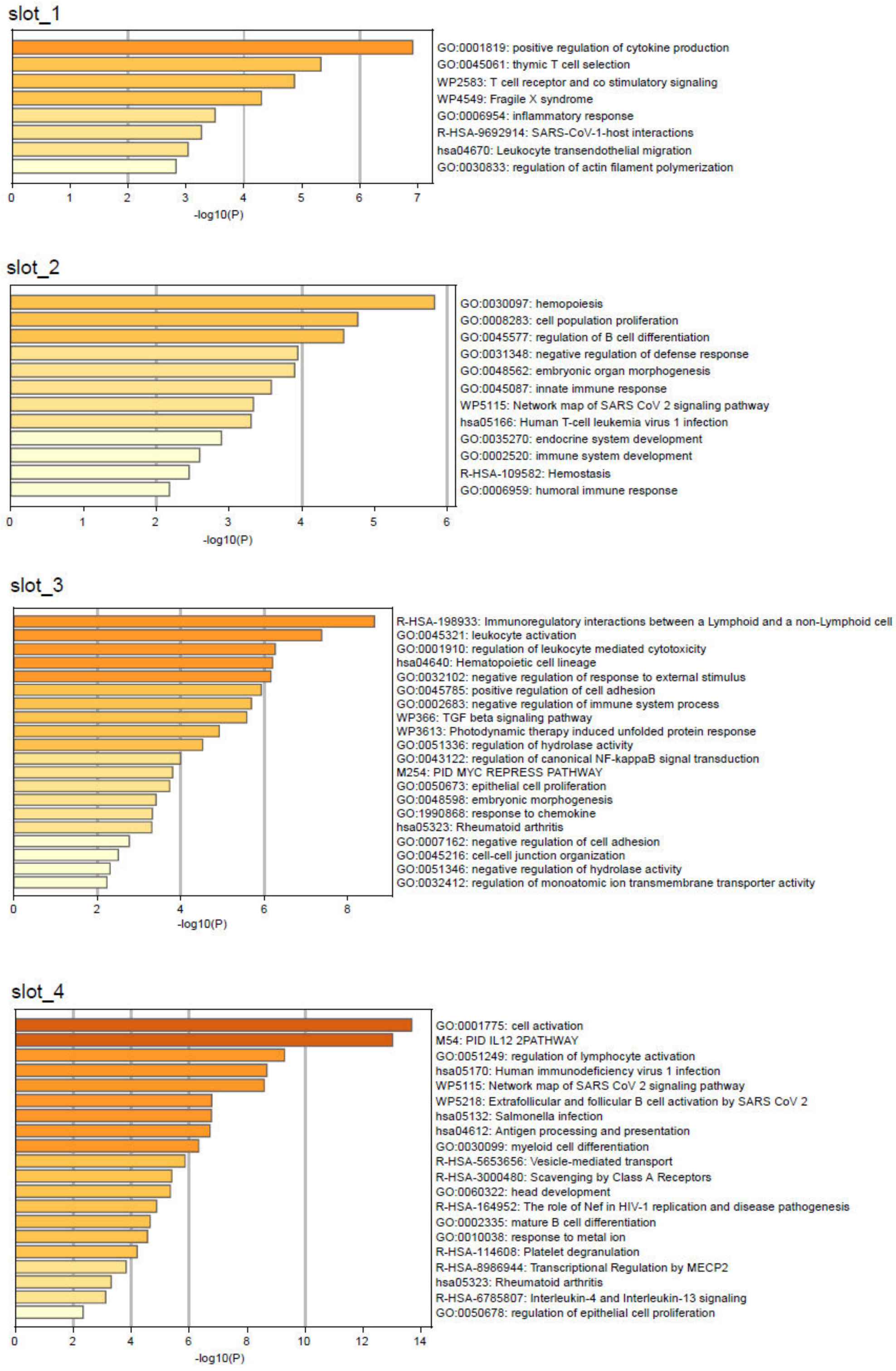

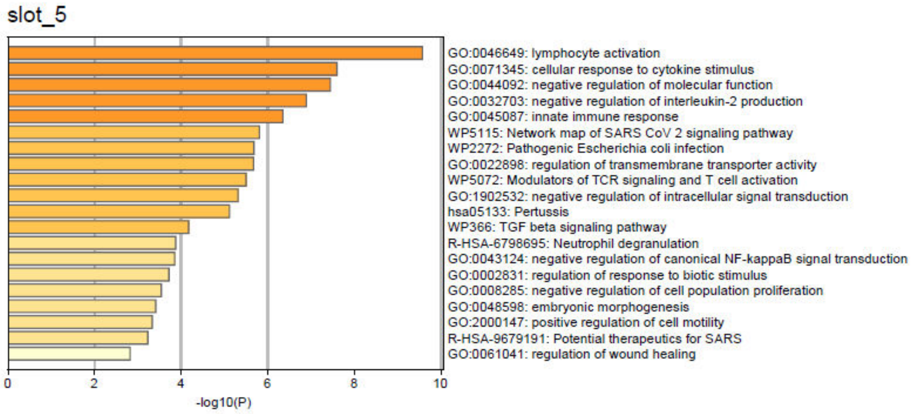
slot_5
GO:0046649: lymphocyte activation
GO:0071345: cellular response to cytokine stimulus
GO:0044092: negative regulation of molecular function
GO:0032703: negative regulation of interleukin-2 production
GO:0045087: innate immune response
WP5115: Network map of SARS CoV 2 signaling pathway
WP2272: Pathogenic Escherichia coli infection
GO:0022898: regulation of transmembrane transporter activity
WP5072: Modulators of TCR signaling and T cell activation
GO:1902532: negative regulation of intracellular signal transduction
hsa05133: Pertussis
WP366: TGF beta signaling pathway
R-HSA-6798695: Neutrophil degranulation
GO:0043124: negative regulation of canonical NF-kappaB signal transduction
GO:0002831: regulation of response to biotic stimulus
GO:0008285: negative regulation of cell population proliferation
GO:0048598: embryonic morphogenesis
GO:2000147: positive regulation of cell motility
R-HSA-9679191: Potential therapeutics for SARS
GO:0061041: regulation of wound healing
0
2
4
6
8
10
-log10(P)

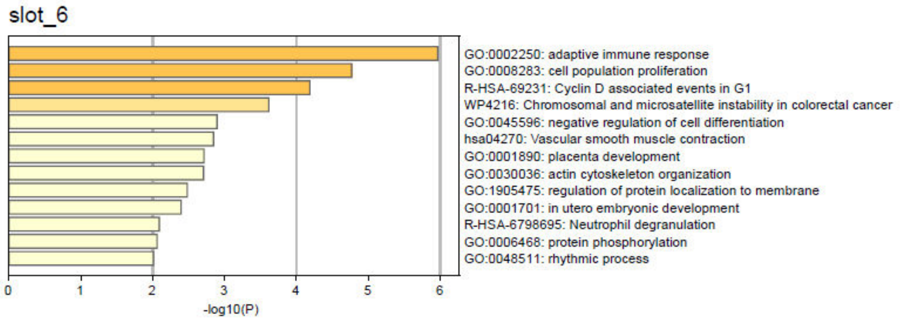
slot_6
GO:0002250: adaptive immune response
GO:0008283: cell population proliferation
R-HSA-69231: Cyclin D associated events in G1
WP4216: Chromosomal and microsatellite instability in colorectal cancer
GO:0045596: negative regulation of cell differentiation
hsa04270: Vascular smooth muscle contraction
GO:0001890: placenta development
GO:0030036: actin cytoskeleton organization
GO:1905475: regulation of protein localization to membrane
GO:0001701: in utero embryonic development
R-HSA-6798695: Neutrophil degranulation
GO:0006468: protein phosphorylation
GO:0048511: rhythmic process
0
1
2
3
4
5
6
-log10(P)

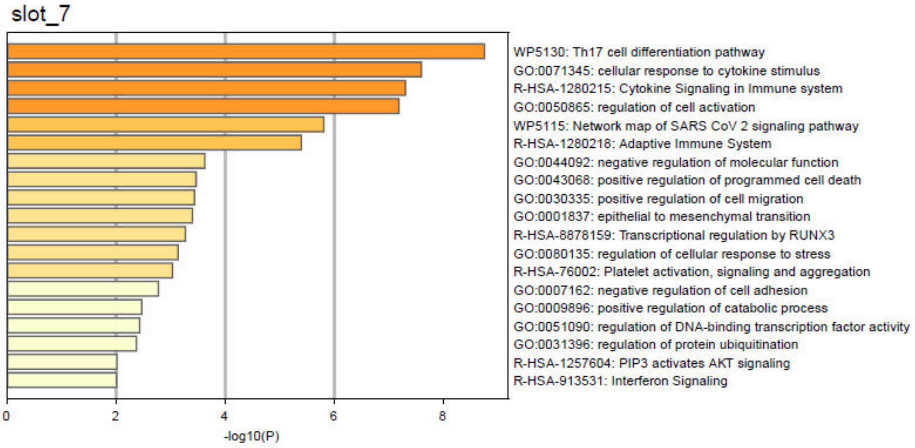
slot_7
WP5130: Th17 cell differentiation pathway
GO:0071345: cellular response to cytokine stimulus
R-HSA-1280215: Cytokine Signaling in Immune system
GO:0050865: regulation of cell activation
WP5115: Network map of SARS CoV 2 signaling pathway
R-HSA-1280218: Adaptive Immune System
GO:0044092: negative regulation of molecular function
GO:0043068: positive regulation of programmed cell death
GO:0030335: positive regulation of cell migration
GO:0001837: epithelial to mesenchymal transition
R-HSA-8878159: Transcriptional regulation by RUNX3
GO:0080135: regulation of cellular response to stress
R-HSA-76002: Platelet activation, signaling and aggregation
GO:0007162: negative regulation of cell adhesion
GO:0009896: positive regulation of catabolic process
GO:0051090: regulation of DNA-binding transcription factor activity
GO:0031396: regulation of protein ubiquitination
R-HSA-1257604: PIP3 activates AKT signaling
R-HSA-913531: Interferon Signaling
0
2
4
6
8
-log10(P)

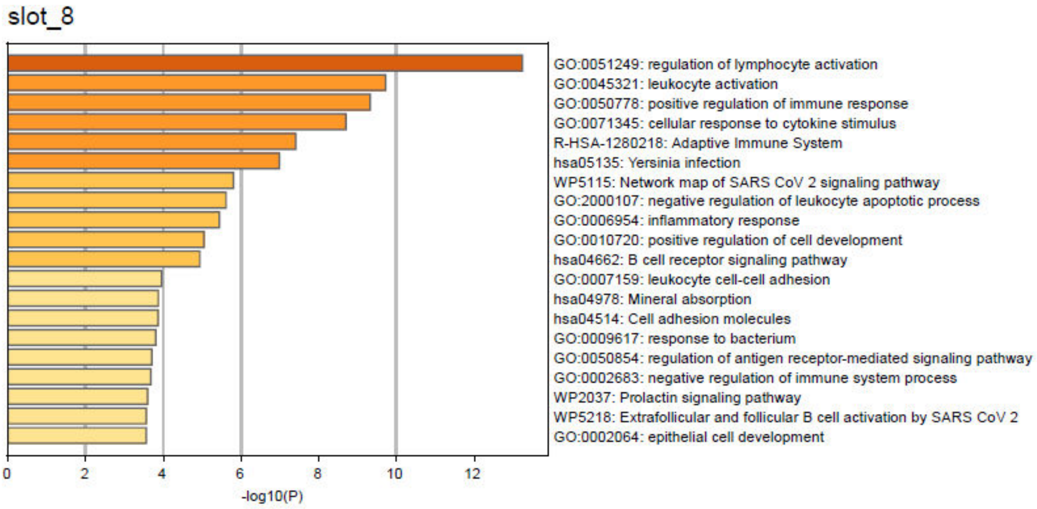


**Supplementary Figure 20. The most significant enrichment terms for each memory slot in the CD8 TCM group.**

#### Supplementary Figure 21

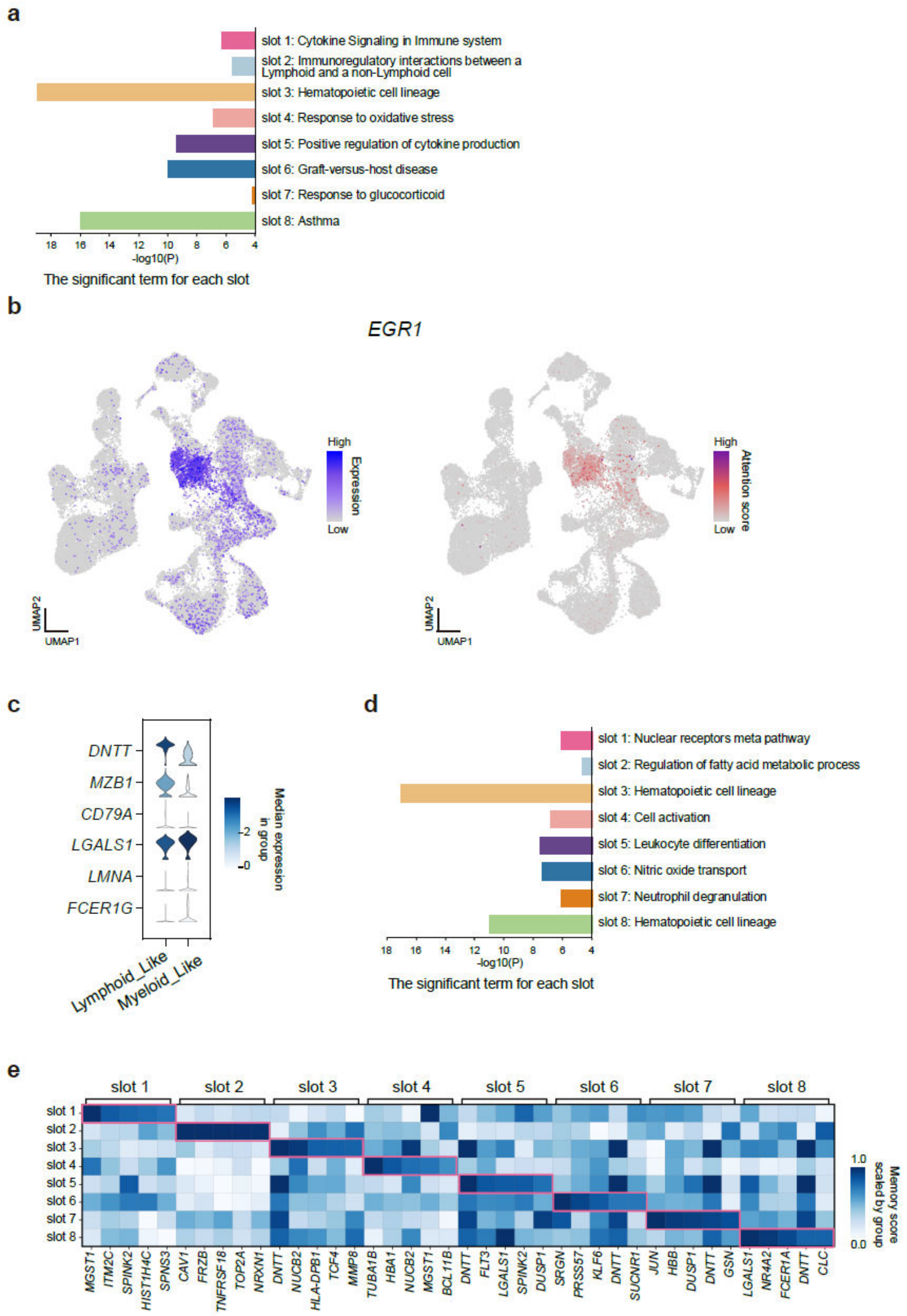


**Supplementary Figure 21. CellMemory's interpretation of complex lineages in MPAL patients.**

**a.** Interpretation of progenitor cells from an MPAL-a patient in memory space, highlighting significant terms from the enrichment results of the top 1-50 TAGs for each memory slot.

**b.** Expression and memory score of *EGR1* in healthy cells and MPAL-b cells.

**c.** Expression of lineage-specific genes in lymphoid and myeloid cells inferred from MPAL-b patient cells.

**d.** Interpretation of progenitor-like cells from MPAL-b in memory space, highlighting significant terms from the enrichment results of the top 1-50 TAGs for each memory slot.

**e.** The heatmap showing the attention level of progenitor-like cells from MPAL-b in memory space.

Eight memory slots are displayed, each showing the top five TAGs that it focuses on.

#### Supplementary Figure 22

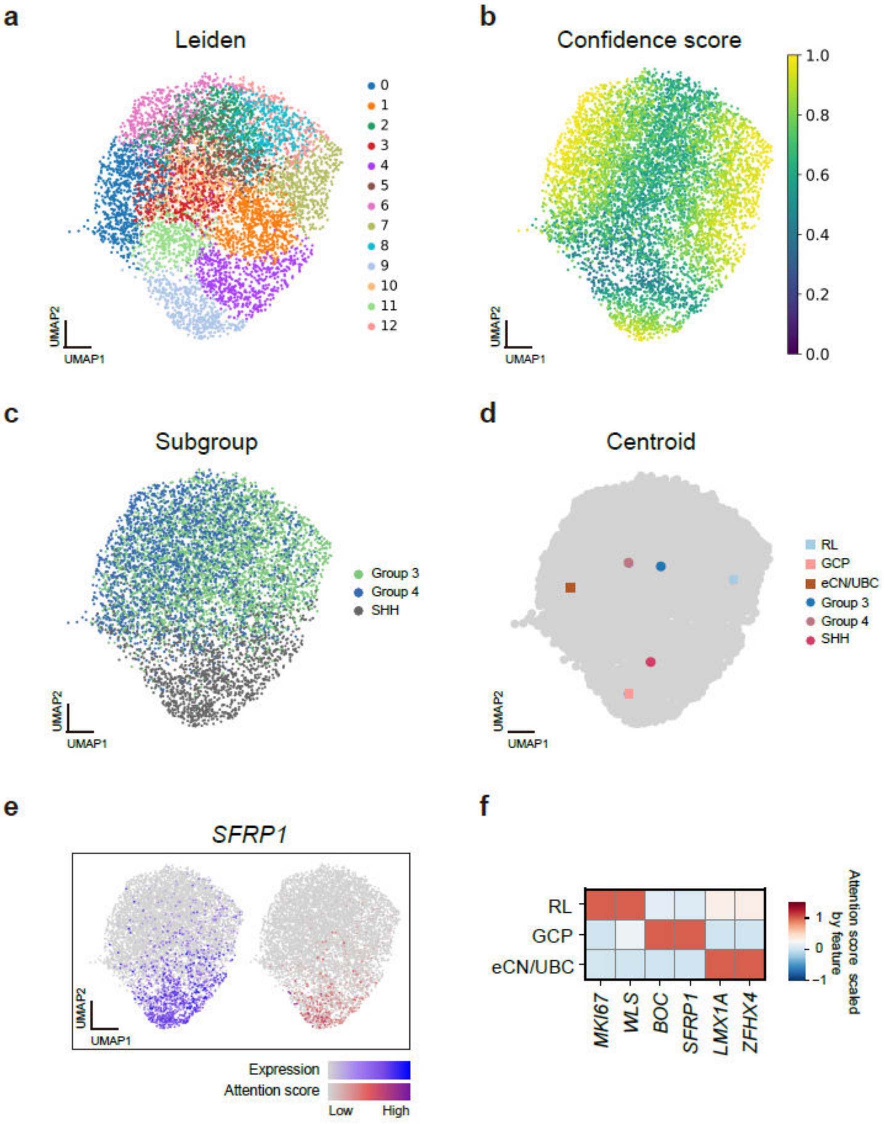


**Supplementary Figure 22. CellMemory's interpretation of heterogeneous origins in different MB subgroups.**

**a.** MB cell embedding with Leiden clustering.

**b.** MB cell embedding with confidence scores.

**c.** MB cells embedding with subgroup labels.

**d.** The centroids of RL, GCP, and eCN/UBC cell types are compared to the centroids of cells from different MB patient subgroups.

**e.** Gene expression and memory score of *SFRP1*.

**f.** Memory scores of *MKI67*, *WLS*, *BOC*, *SFRP1*, *LMX1A*, and *ZFHX4* in MB cells with different inferred origins.

#### Supplementary Figure 23

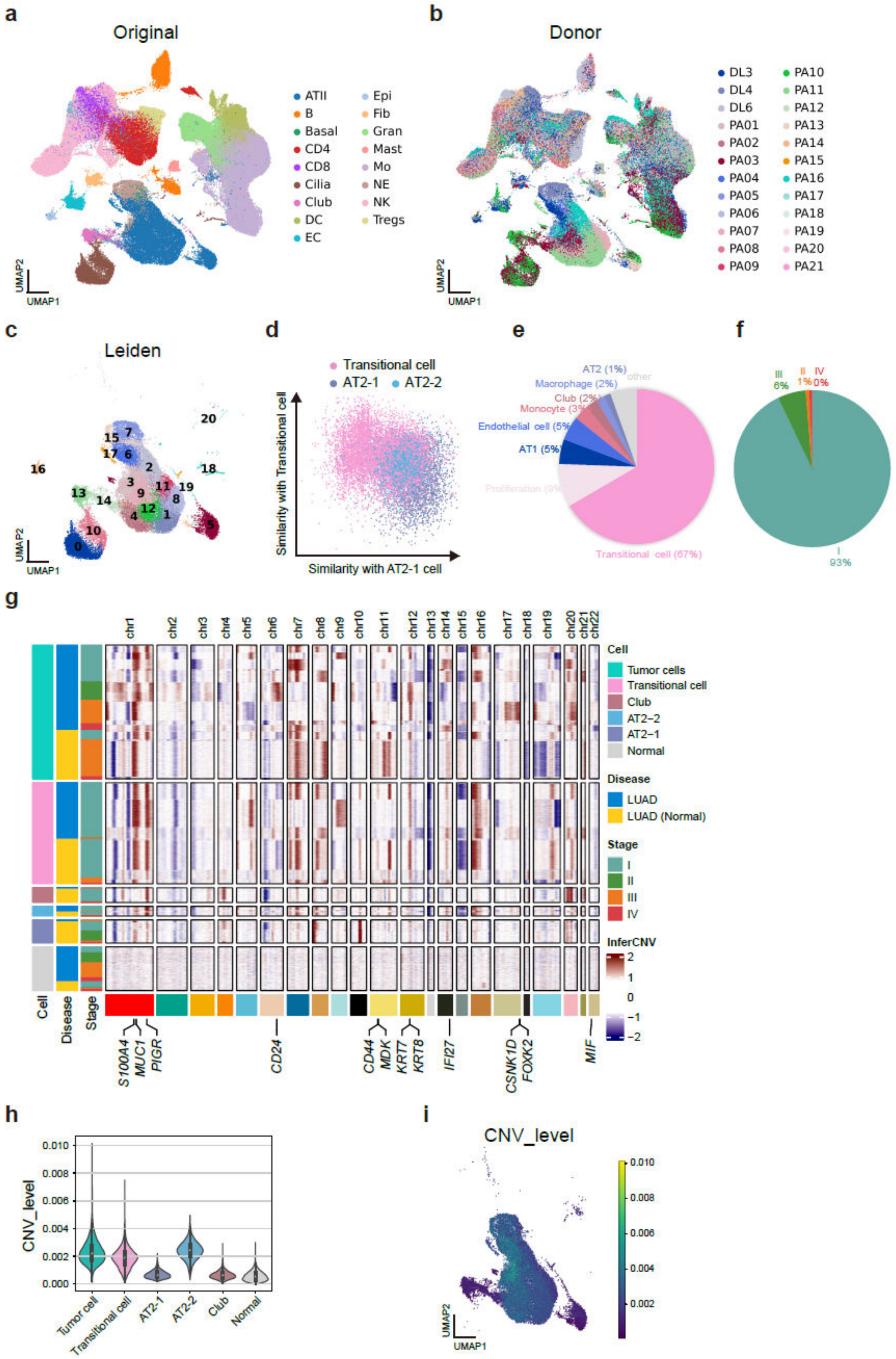


**Supplementary Figure 23. Integration of single-cell data in LUAD.**

**a-b.** UMAP embeddings generated by CellMemory, colored by original label (**a**) and donor label (**b**).

**c.** Leiden clustering of epithelial cell embeddings.

**d.** Similarity of Transitional cells, AT2-1 cells, and AT2-2 cells to Transitional cell features (Y-axis) and AT2-1 cell features (X-axis).

**e.** Construction of CellMemory using the same reference excluding tumor cells, predicting the cell types most similar to tumor cells in LUAD patients.

**f.** Proportion of stages in the Transitional cell.

**g.** CNV patterns of tumor-related epithelial cells, inferred by inferCNV.

**h.** CNV levels in tumor-related epithelial cells.

**i.** UMAP embeddings of tumor-related epithelial cells colored by CNV levels.

#### Supplementary Figure 24

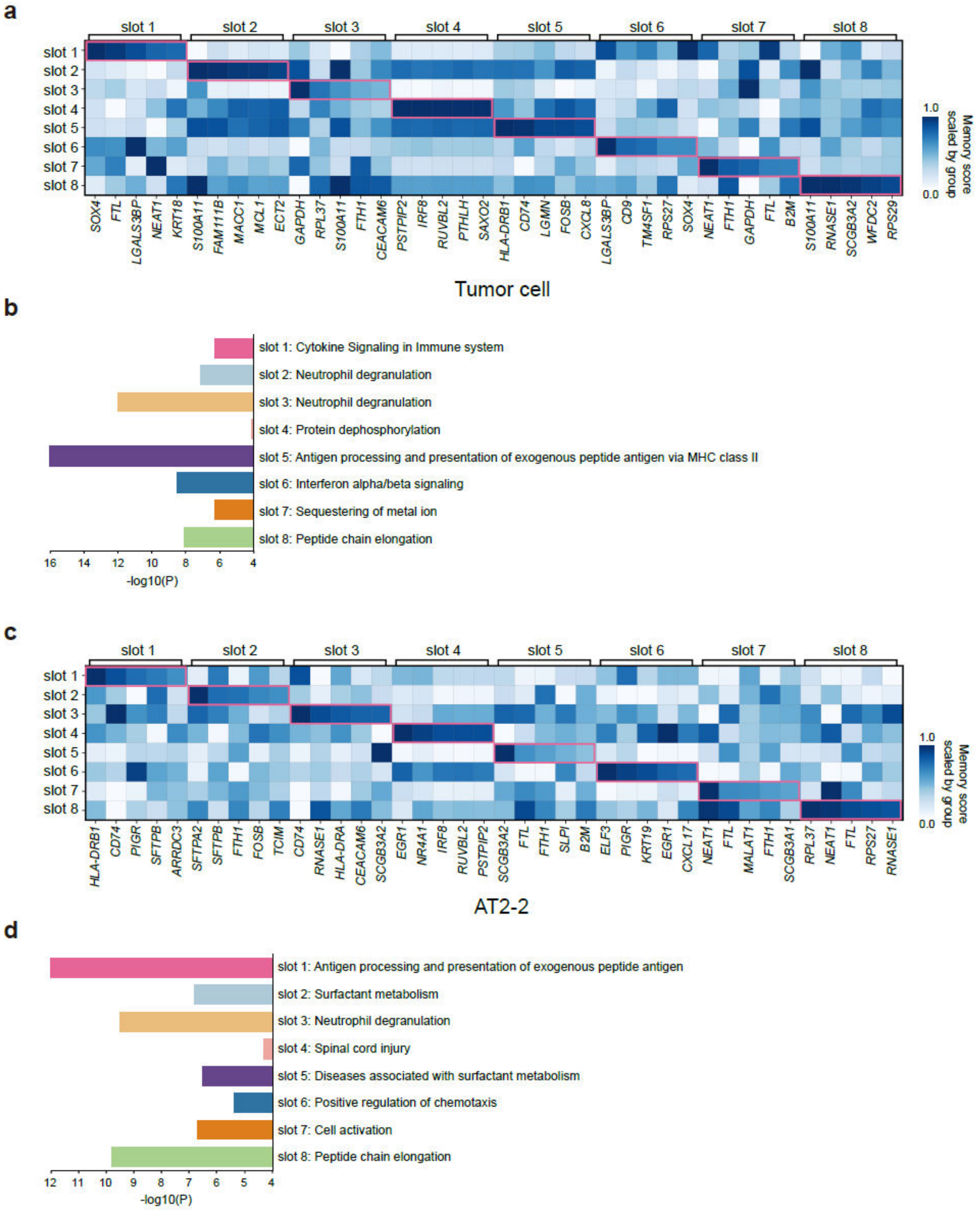


**Supplementary Figure 24.    Hierarchical interpretation of Tumor cell and AT2-2 cell.**

**a.** The heatmap showing the attention level of Tumor cells from the LUAD dataset in memory space. Eight memory slots are displayed, each showing the top five TAGs that it focuses on.

**b.** The top 1-50 TAGs for each slot of Tumor cell were subjected to enrichment analysis, with significant terms displayed.

**c.** The heatmap showing the attention level of AT2-2 cells from the LUAD dataset in memory space. Eight memory slots are displayed, each showing the top five TAGs that it focuses on.

**d.** The top 1-50 TAGs for each slot of AT2-2 cell were subjected to enrichment analysis, with significant terms displayed.

#### Supplementary Figure 25

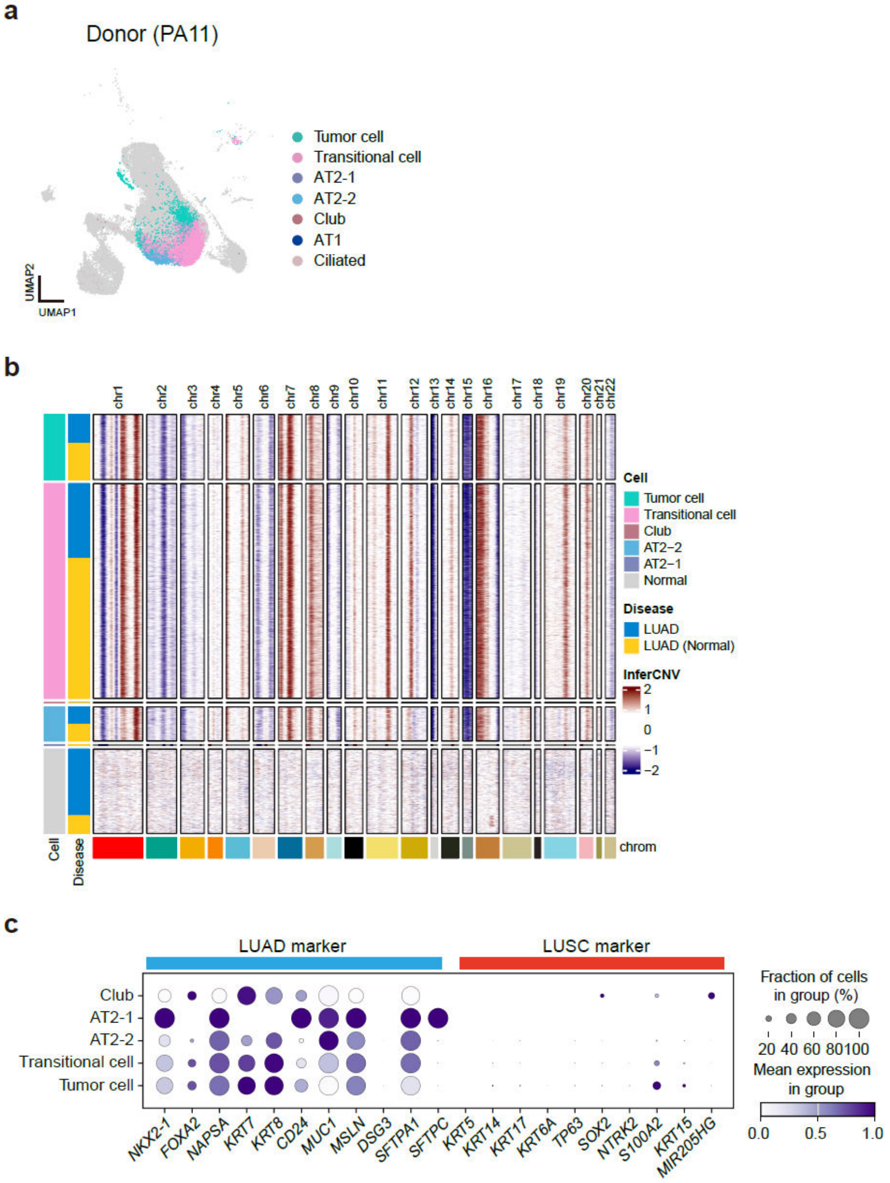


**Supplementary Figure 25. Characterization and CNV of PA11 cells.**

**a.** Epithelial cells of PA11 are highlighted in UMAP.

**b.** CNV patterns in tumor-related epithelial cells of PA11.

**c.** Tumor markers (LUAD and LUSC) expressed in tumor-related epithelial cells of PA11.

#### Supplementary Figure 26

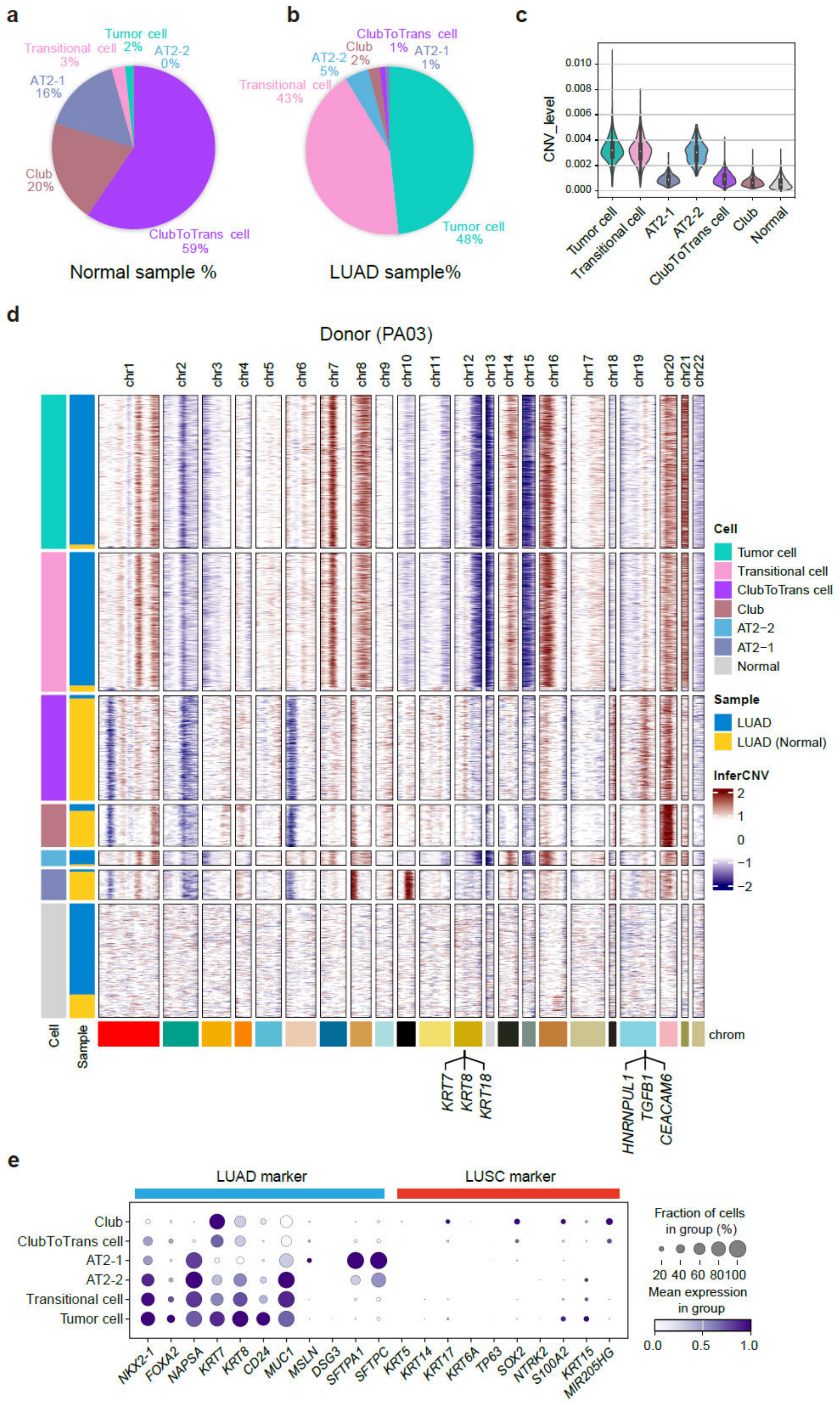


**Supplementary Figure 26. Characterization and CNV of PA03 cells.**

**a-b.** The proportion of tumor-related epithelial cells in the normal sample (**a**) and LUAD sample (**b**) of PA03.

**c.** CNV levels in tumor-related epithelial cells of PA03 patients.

**d.** CNV patterns in tumor-related epithelial cells of PA03.

**e.** Tumor markers (LUAD and LUSC) expressed in tumor-related epithelial cells of PA03.

#### Supplementary Figure 27

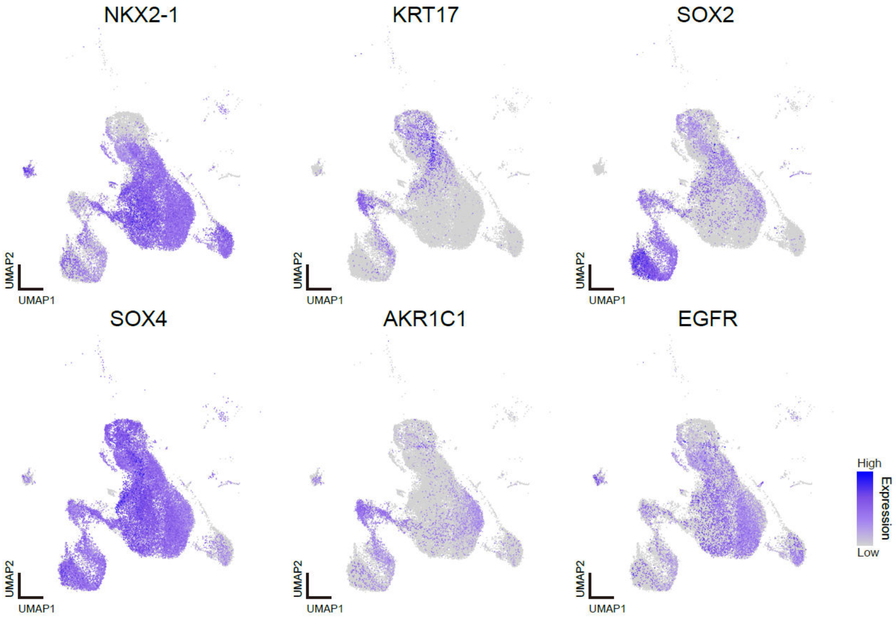


**Supplementary Figure 27. Expression of genes related to LUAD and LUSC in epithelial cells of LUAD patients.**

### Supplementary Figure 28

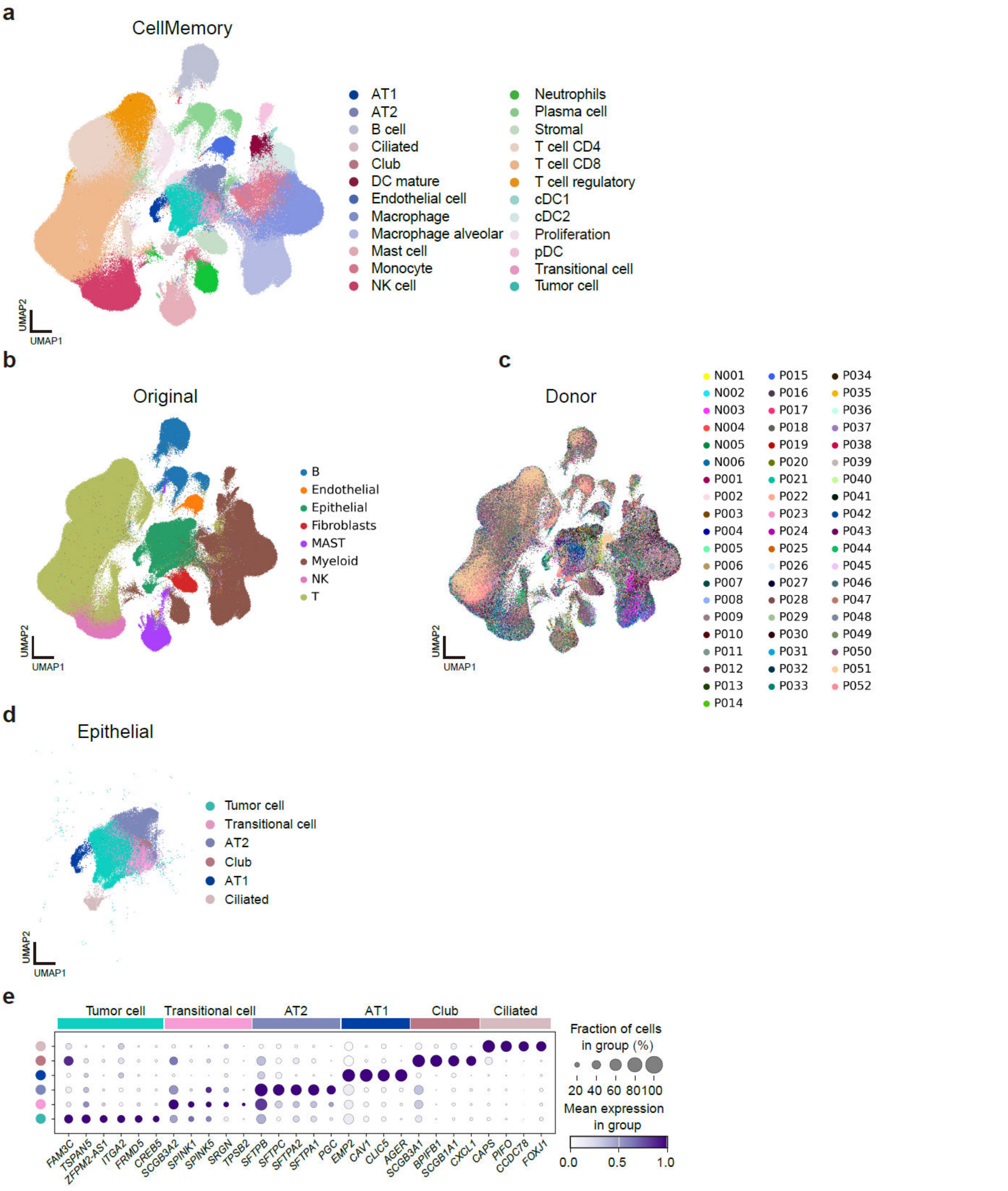


**Supplementary Figure 28. Integration of LUAD dataset 2.**

**a.** UMAP generated by CLS embedding, integrating, and annotating 488,236 cells from LUAD donors.

**b-c.** UMAP embeddings colored by original label (**b**) and donor label (**c**).

**d.** UMAP of epithelial cells in the LUAD dataset 2.

**e.** Dot plot demonstrating specific gene expression in epithelial cells.

#### Supplementary Figure 29

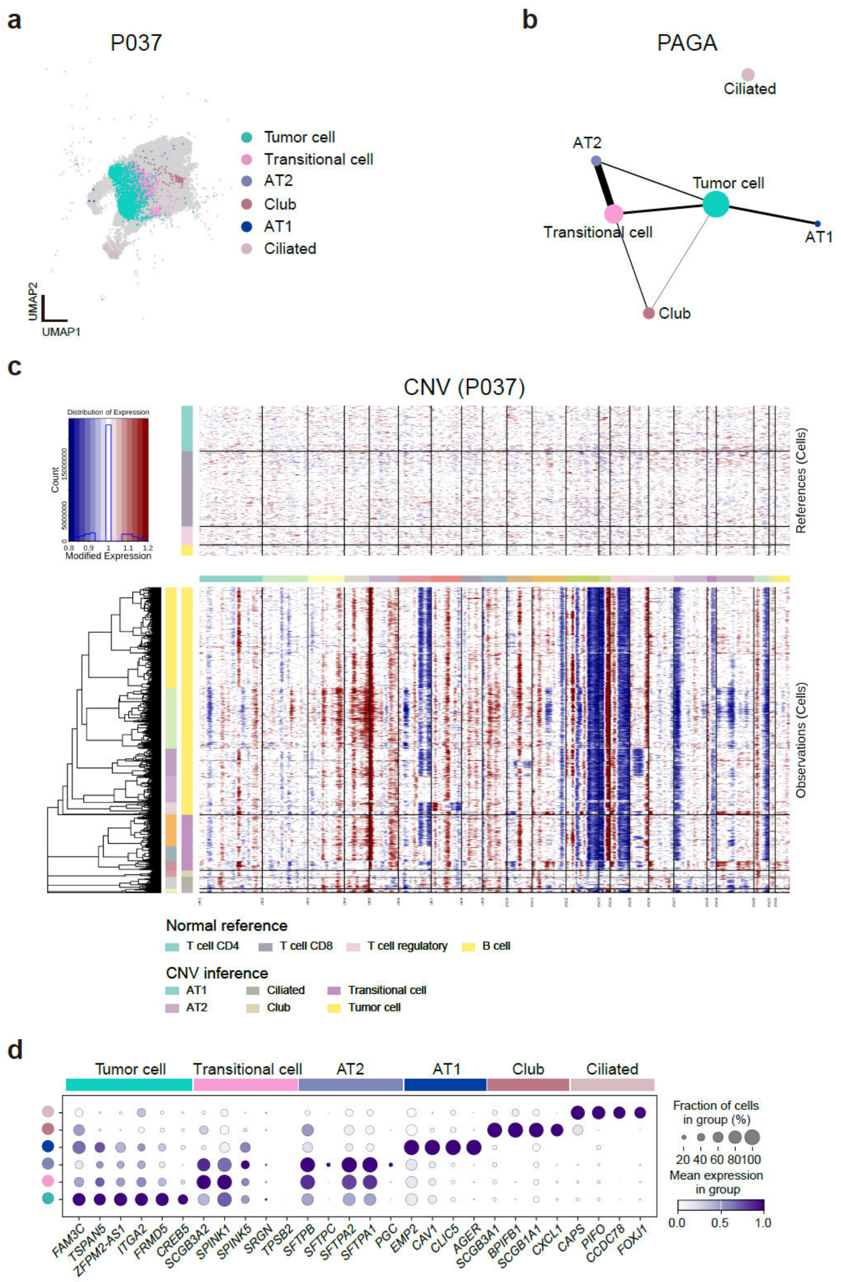


**Supplementary Figure 29. Characterization and CNV of P037 cells in LUAD dataset 2.**

**a.** Epithelial cells of P037 are highlighted in UMAP.

**b.** Partition-based graph abstraction (PAGA) illustrates connections between separated cells, with edge weights representing confidence in the connections between epithelial cells. The size of the dots represents the number of cells in each cluster. The similarity between AT2 and Transitional cells in the epithelial cells of P037.

**c.** CNV patterns in all epithelial cells of P037.

**d.** Dot plot demonstrating specific gene expression in epithelial cells of P037. AT2 cells show gene expression similarities with Transitional cells and Tumor cells.

#### Supplementary Figure 30

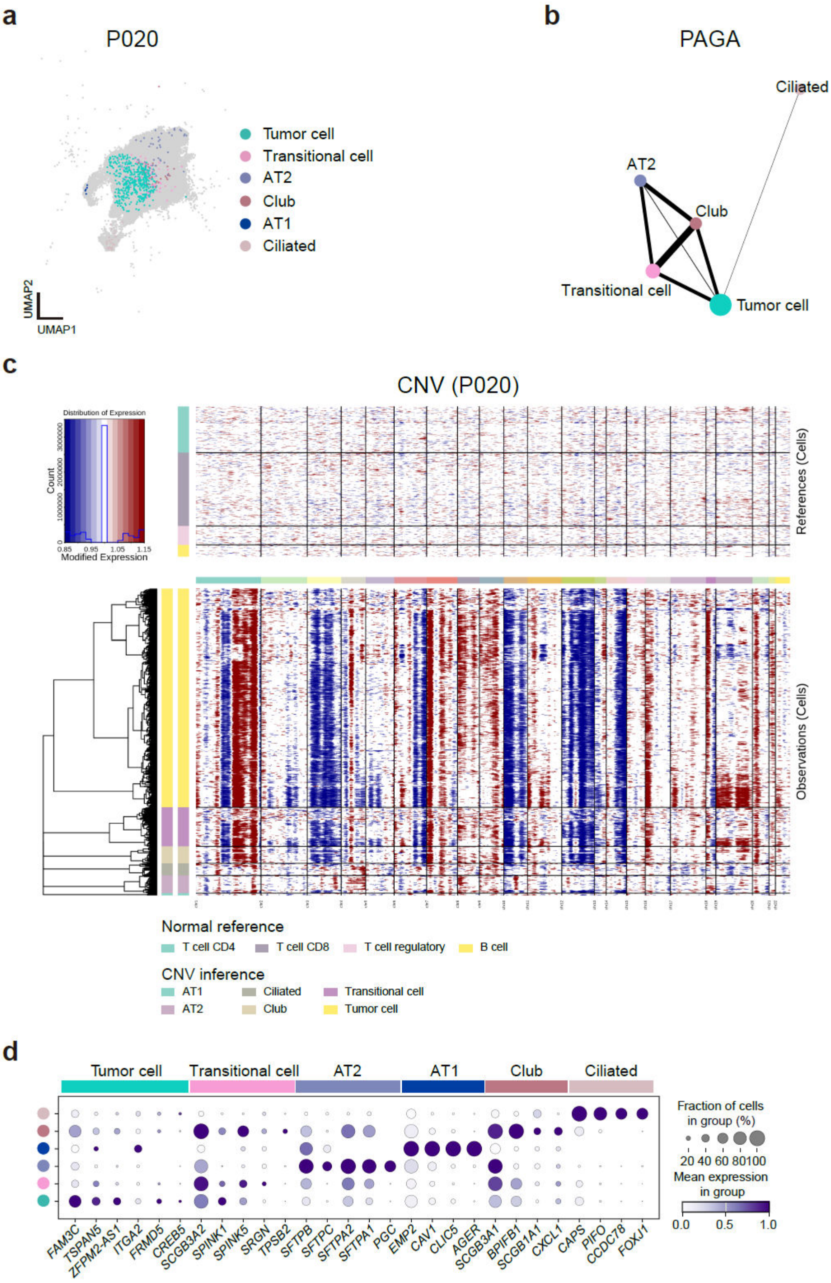


**Supplementary Figure 30. Characterization and CNV of P020 cells in LUAD dataset 2.**

**a.** Epithelial cells of P020 are highlighted in UMAP.

**b.** Partition-based graph abstraction (PAGA) illustrating connections between epithelial cells. The significant similarity between Club cells, Transitional cells, and Tumor cells in the epithelial cells of P020, indicates a relationship between club cells and tumor cells.

**c.** CNV pattern in epithelial cells of P020. Club cells exhibit a CNV Pattern very similar to Tumor cells. The CNV patterns of AT2 cells do not appear to be very similar.

**d.** Dot plot demonstrating specific gene expression in epithelial cells of P020. Club cells show gene expression similarities with Transitional cells and Tumor cells.

#### Supplementary Figure 31

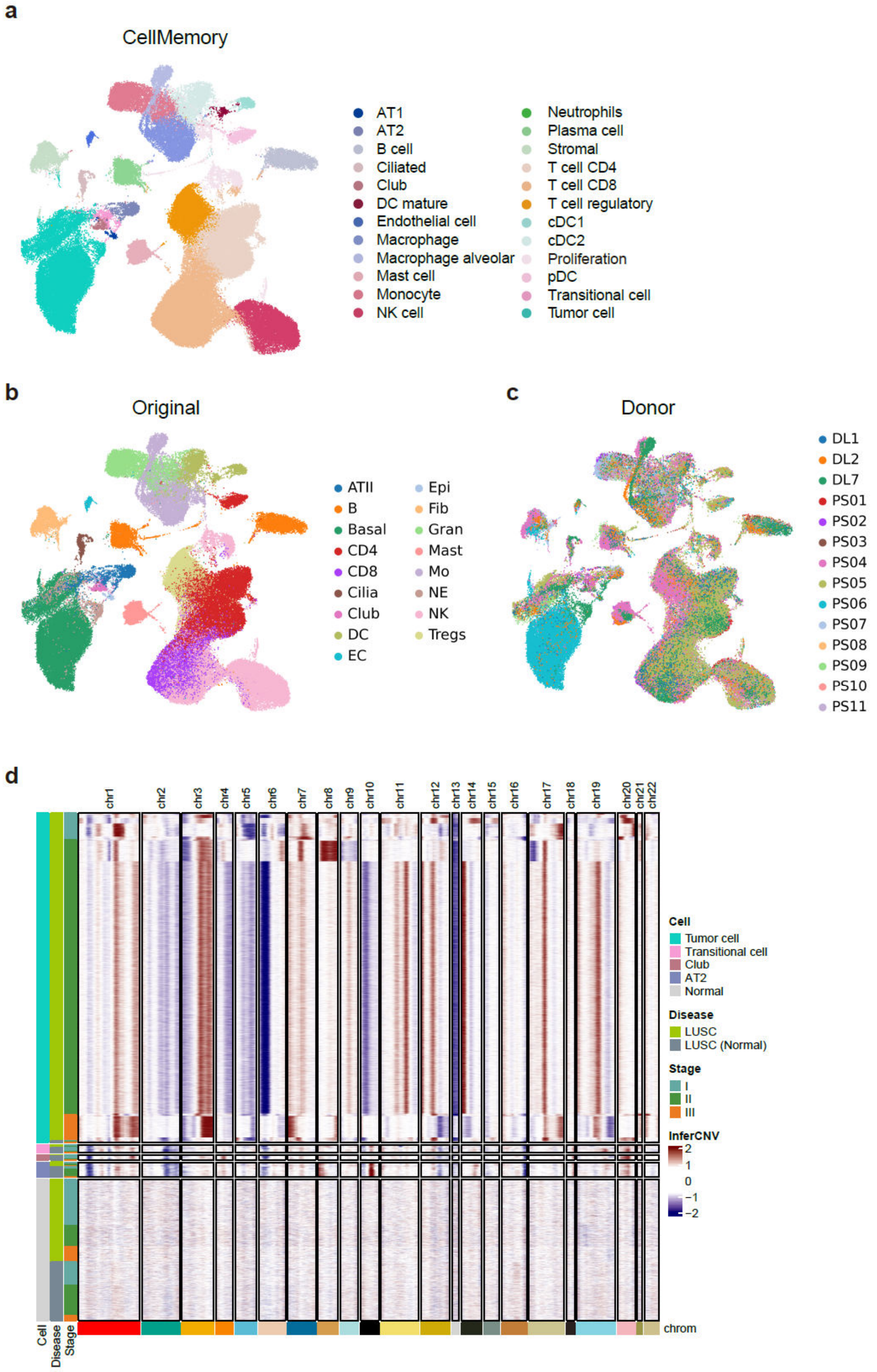

e

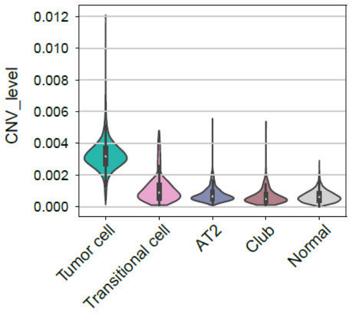


f

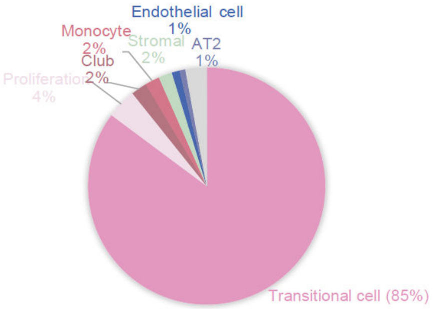


g

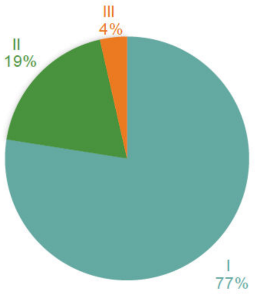


**Supplementary Figure 31. CellMemory analysis of LUSC heterogeneity.**

**a-c.** UMAP embeddings of LUSC cells generated by CellMemory, with predicted label (**a**), original label (**b**), and donor information (**c**).

**d.** CNV in tumor-related epithelial cells of all LUSC patients.

**e.** CNV levels in tumor-related epithelial cells of LUSC patients.

**f.** Construction of a model using the same reference excluding tumor cells, predicting the epithelial cell types most similar to tumor cells in LUSC patients. Transitional cell has the highest percentage.

**g.** Proportion of stages in the Transitional cell.

#### Supplementary Figure 32

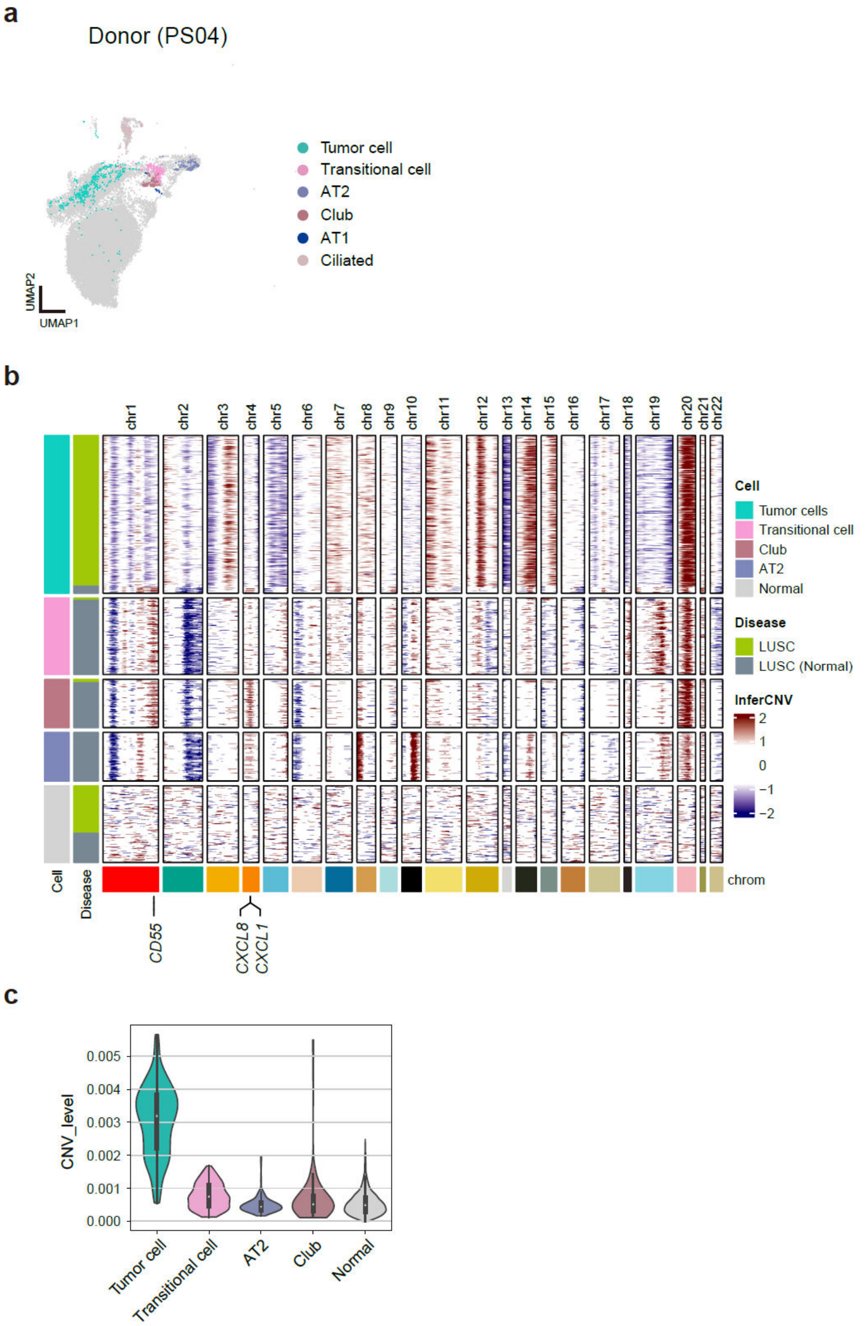


**Supplementary Figure 32. Characterization and CNV of PS04 cells in LUSC.**

**a.** Representation of individual PS04 in cell embeddings.

**b.** CNV results in tumor-related epithelial cells of PS04.

**c.** CNV levels in tumor-related epithelial cells of PS04.

### Supplementary Figure 33

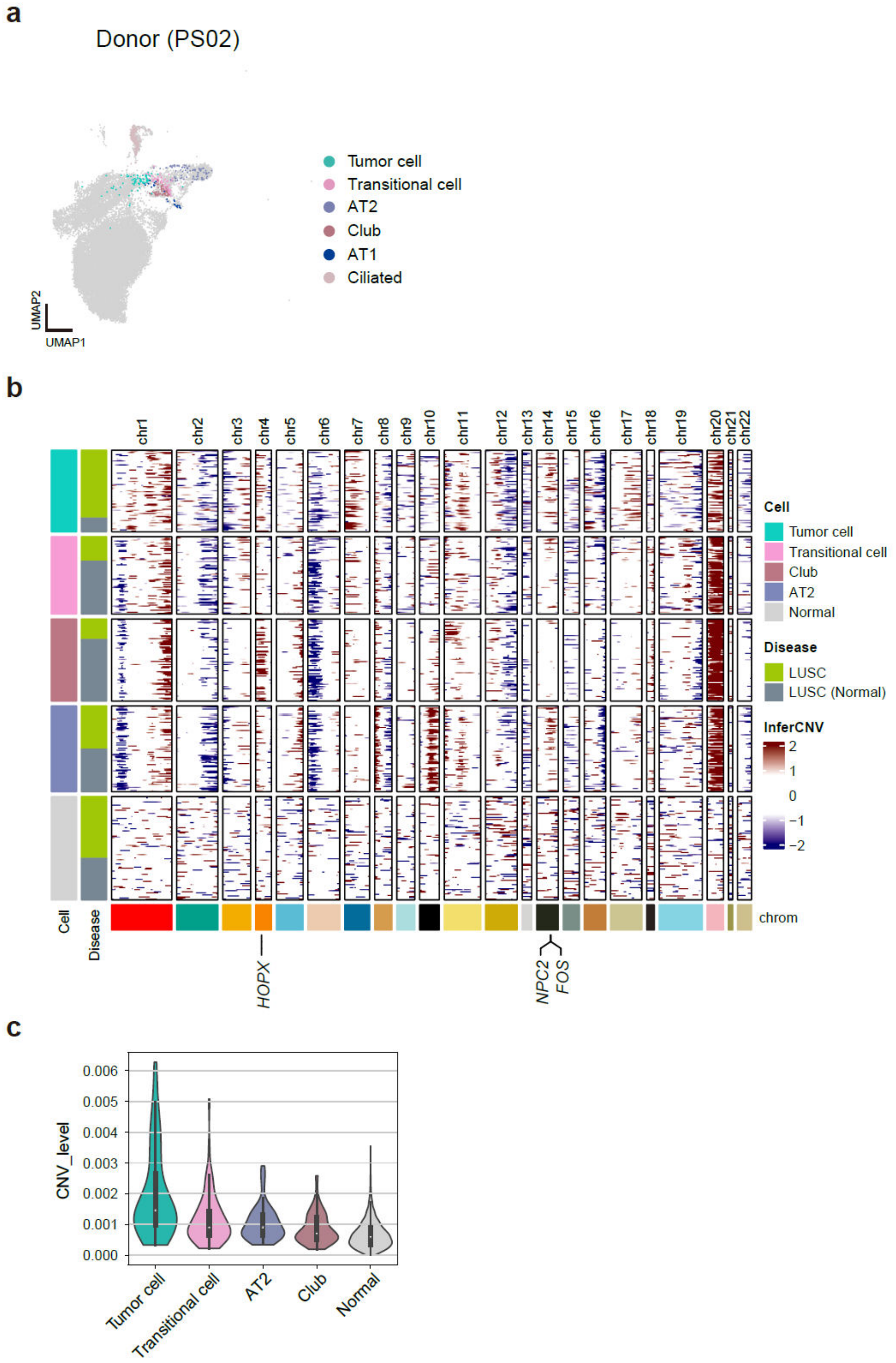


**Supplementary Figure 33. Characterization and CNV of PS02 cells in LUSC.**

**a.** Representation of individual PS02 in cell embeddings.

**b.** CNV results in tumor-related epithelial cells of PS02.

**c.** CNV levels in tumor-related epithelial cells of PS02.

#### Supplementary Table 1

**Hyperparameters list**

| Parameter | Value |
|---|---|
| Batch size | 140 |
| Learning rate | 0.0003 |
| Learning rate scheduler | CosineAnnealingLR (T_max=200) |
| Optimizer | Adam |
| Loss | CrossEntropyLoss |
| Dropout | 0.1 |
| Top k | 20 |
| Embedding dims | 256 |
| Head dims | 256 |
| Forward dims | 512 |
| Heads num | 4 |
| Memory slots | 8 |
| Number of layers | 4 |
| Early stopping (patience) | 5 |
| Shared attention layer | True |

We used 1 GPU (RTX4090) to benchmark.

**Supplementary Table 2**

Data availability list.

**Supplementary Table 3**

The annotation performance of single-cell datasets.

**Supplementary Table 4**

The integration performance of single-cell datasets.

**Supplementary Table 5**

The integration performance under different token processing strategies.

**Supplementary Table 6**

The annotation performance of spatial datasets.

**Supplementary Table 7**

The TAGs of the AIDA dataset.

**Supplementary Table 8**

The TAGs of the MPAL dataset.

**Supplementary Table 9**

The TAGs of the LUAD dataset.
